# Tauopathy in the *APP*_swe_/*PS1*_ΔE9_ mouse model of familial Alzheimer’s disease

**DOI:** 10.1101/405647

**Authors:** Athanasios Metaxas, Camilla Thygesen, Stefan J. Kempf, Marco Anzalone, Ramanan Vaitheeswaran, Sussanne Petersen, Anne M. Landau, Hélène Audrain, Jessica L. Teeling, Sultan Darvesh, David J. Brooks, Martin R. Larsen, Bente Finsen

## Abstract

Despite compelling evidence that the accumulation of amyloid-beta (Aβ) promotes cortical MAPT (tau) aggregation in familial and idiopathic Alzheimer’s disease (AD), murine models of cerebral amyloidosis are not considered to develop tau-associated pathology. The absence of neurofibrillary lesions in amyloidosis mice remains a challenge for the amyloidocentric paradigm of AD pathogenesis. It has resulted in the generation of transgenic mice harboring mutations in their *tau* gene, which may be inappropriate for studying a disease with no known *TAU* mutations, such as AD. Here, we have used *APP*_*swe*_/*PS1*_*ΔE9*_ mice to show that tau pathology can develop spontaneously in murine models of familial AD. Tauopathy was abundant around Aβ deposits, with Gallyas- and thioflavin-S-positive perinuclear inclusions accumulating in the *APP*_*swe*_/*PS1*_*ΔE9*_ cortex by 18 months of age. Age-dependent increases in Gallyas signal correlated positively with binding levels of the paired helical filament (PHF) ligand [^18^F]Flortaucipir, in all brain areas examined. Sarkosyl-insoluble PHFs were visualized by electron microscopy. Tandem mass tag proteomics identified sequences of hyperphosphorylated tau in transgenic mice, along with signs of RNA missplicing, ribosomal dysregulation and disturbed energy metabolism. Human frontal gyrus tissue was used to validate these findings, revealing primarily quantitative differences between the tauopathy observed in AD patient vs. transgenic mouse tissue. Levels of *tau* mRNA were not different between *APP*_*swe*_/*PS1*_*ΔE9*_ and littermate control animals. As physiological levels of endogenous, ‘wild-type’ tau aggregate secondarily to Aβ in transgenic mice, this study demonstrates that amyloidosis is both necessary and sufficient to drive tauopathy in experimental models of familial AD.

## Introduction

Genetically-inherited and sporadic forms of Alzheimer’s disease (AD) are characterized by a common set of hallmark brain lesions, which include the accumulation of amyloid-β (Aβ) peptides into plaques, neuroinflammation, aggregation of hyperphosphorylated MAPT (tau) into neurofibrillary tangles (NFTs), and neurodegeneration. Transgenic mouse models that reproduce aspects of the aforementioned lesions have been generated based on mutations in the amyloid precursor protein (*APP*) and presenilin 1 (*PSEN1*) and 2 (*PSEN2*) genes, which are known to cause familial AD (*1*). Despite playing important roles in evaluating APP processing, Aβ toxicity, and amyloid-targeting therapeutic strategies, transgenic mice are not being regarded as models that can replicate the full spectrum of AD histopathology (*2*). In particular, while the overexpression of mutant *APP* and *APP*/*PSEN1* has been shown to yield amyloidosis (*3*), neuroinflammation (*4*) and neurodegeneration (*5*) in mice, it is generally not considered to promote the conversion of endogenous tau into neurofibrillary structures (*6*).

To address the *in vivo* role of tau hyperphosphorylation and NFT formation in AD pathogenesis, human *MAPT* (*TAU*) has been introduced into the mouse genome, either mutated or non-mutated, on a *Tau*-knockout background (*7, 8*). *TAU* overexpressing mice demonstrate progressive neurofibrillary pathology, albeit in the marked absence of cerebral amyloidosis, which is required for a neuropathological diagnosis of AD. Moreover, mutations in *TAU* have been linked to non-AD tauopathies, most commonly frontotemporal lobar degeneration [FTLD; (*9*)], a condition with neuropathological hallmarks distinct from AD. Thus, murine models of amyloidosis and combined amyloidosis-tauopathy models have been widely criticized for their translational relevance to the human condition. It has been argued that virtually all existing murine models would be considered as ‘not’ AD (*10*) according to the ABC scoring system of neuropathology (*11*). The inability of amyloidosis mice to develop neurofibrillary lesions is thought to contribute to the poor translation of preclinical research into clinical benefits (*12*), and has raised concern about the amyloidocentric model of AD pathogenesis (*13*).

Two principal explanations have been put forward for the lack of tau-associated pathology in amyloidosis mice (*14*). First, adult mice express fewer isoforms of the tau protein than humans (three vs. six), which might render them less liable to the post-translational modifications (PTMs) that are associated with the accumulation of tau into NFTs, such as phosphorylation (*15*). However, murine tau has been shown to readily fibrillize *in vitro* upon treatment with polyanionic factors, including RNA (*16*), and there is ample evidence of tau hyperphosphorylation in the transgenic mouse brain [(*17*), Table S1], indicating that no differences exist in the propensity of murine and human tau for aggregation and PTMs. A second reason that is often cited for the absence of tauopathy in amyloidosis models is that the murine lifespan may be too short for the complete sequence of neurofibrillary pathology to unfold in transgenic mice. Although age scaling studies suggest otherwise (*18*), the aging factor has been neglected in the design of preclinical studies.

Transgenic Fischer rats (TgF344-AD), expressing human *APP* harboring the Swedish double mutations (KM670/671NL) and *PSEN1* lacking exon 9 (*APP*_swe_/*PS1*_ΔE9_), both under control of the mouse prion protein promoter, develop progressive neurofibrillary pathology (*19*). In this study, transgenic *APP*_swe_/*PS1*_ΔE9_ mice that were constructed in an identical manner as TgF344-AD rats were used to demonstrate neurofibrillary pathology in aging amyloidosis mice.

## Results

### Neurofibrillary pathology in aging APP_swe_/PS1_ΔE9_ mice

Fresh-frozen brain sections from 3-, 6-, 12-, 18- and 24-month-old *APP*_swe_*/PS1*_ΔE9_ transgenic (TG) mice and their wild-type (WT) counterparts were processed for the detection of neurofibrillary alterations with the Gallyas silver stain (n=6/group). Thioflavin-S and DAPI (4′,6-diamidino-2-phenylindole) were used to detect perinuclear β-pleated structures. Co-staining for amyloid and Gallyas was used to probe the relationship between amyloidosis and tau-associated pathology in aging TG animals. Fresh-frozen sections of the middle frontal gyrus from a patient with definite AD were processed in parallel with sections from *APP*_swe_*/PS1*_ΔE9_ mice, to compare Gallyas-positive structures in mouse vs. human tissue.

Aβ deposition was the predominant lesion in the 6-month-old *APP*_swe_*/PS1*_ΔE9_ brain (Fig. 1A&B), with age-dependent increases in argyrophilic density observed exclusively in TG mice (Fig. 1C-F). Only mild and diffuse silver staining was observed in the neocortex of 6-month-old animals (Fig. 1G). Densely-labeled, round structures, surrounded by a halo of argyrophilic staining, constituted the majority of Gallyas-positive signal in the neuropil of the neocortex and hippocampus at 12-24 months of age (Fig. 1H&I). In addition, diffuse and compact argyrophilic staining was observed surrounding red-stained nuclei in the neocortex of 18- and 24-month-old *APP*_swe_*/PS1*_ΔE9_ mice (Fig. 1J&K). The perinuclear structures were positive for thioflavin-S (Fig. 1L-N), which colocalized with nuclear DAPI (Fig. 1O) and was further detected in cell-sized structures lacking a stainable nucleus (Fig. 1P). There were no apparent differences in morphology between the argyrophilic structures in brain tissue from 24-month-old TG mice (Fig 1Q-U) and AD-confirmed patient material (Fig. 1V-Z), although neuropil threads were detected exclusively in AD tissue (Fig. 1Q-Z). Coronal brain sections of 20-month-old Tg2576 mice, harboring the Swedish double mutations, were used to examine 6E10- and Gallyas-positive pathology in a second mouse model of amyloidosis (Fig. 1AA-AD). Amorphous argyrophilic signal (AC) and perinuclear lesions (AD) were also present in the Tg2576 mouse brain, albeit at lower levels compared to 18-month-old *APP*_swe_/*PS1*_ΔE9_ mice.

**Fig. 1.**
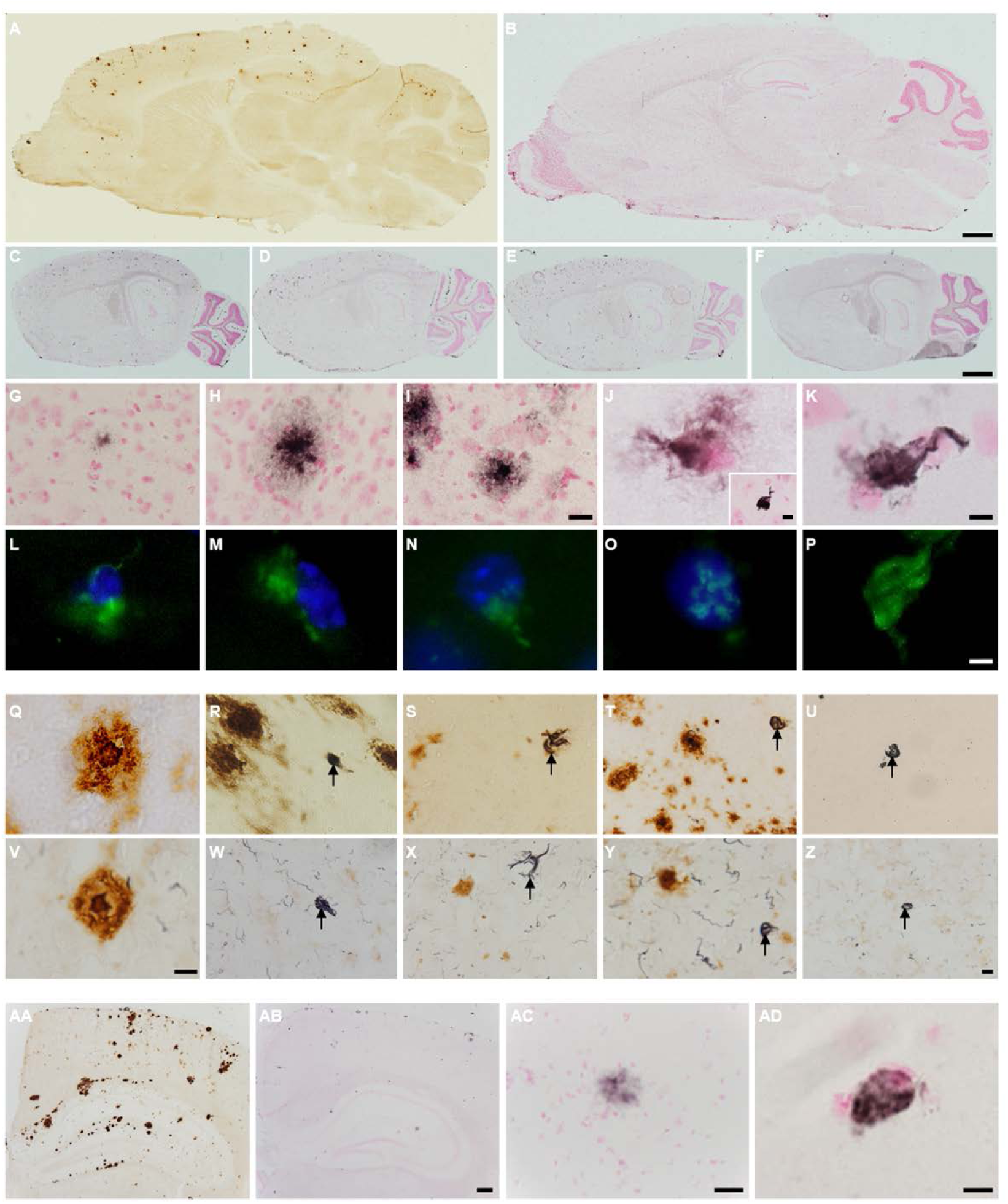
Neurofibrillary alterations in amyloidosis mice. (A&B) Sagittal brain sections of 6-month-old *APP*_swe_/*PS1*_ΔE9_ mice, processed for 6E10 immunohistochemistry (A) and the Gallyas silver stain (B). Silver-labeled sections were counterstained with nuclear fast red. β-amyloidosis dominates over argyrophilic pathology in 6-month-old *APP*_swe_/*PS1*_ΔE9_ mice. **(C-F)** Progressive increase in Gallyas-positive signal in 12- (C), 18- (D), and 24-month-old transgenic mice (E). Wild-type animals showed no silver deposition up to 24 months of age (F). **(G-P)** All photomicrographs are from the neocortex of *APP*_swe_/*PS1*_ΔE9_ miceArgyrophilic signal was scarce in 6-month-old TG animals (G). Gallyas-positive structures in 18- (H) and 24-month-old animals (I), likely of neuritic nature. Gallyas silver (J & K) and thioflavin-S stainings (L-P), showing perinuclear and intranuclear signal in 18- and 24- month-old transgenic mice. The insert in J shows compact Gallyas staining in the absence of nuclear fast red. Note potential fragmented nuclei in (M) and (N), intranuclear signal in (O), and absence of DAPI signal in (P). **(Q-Z)** Gallyas/6E10 doubly-labeled sections from a 24-month-old transgenic mouse (Q-U) and an AD patient (V-Z), showing dense-core plaques (Q & V), teardrop-shaped structures (R & W, arrows), tuft-shaped filaments (S & X, arrows), and globose structures in close proximity (T & Y) and over 200 µm afar from Aβ deposits (U & Z). **(AA- AD)** 6E10/Gallyas- AA) and Gallyas-labeled (AB-AD) sections of 20-month-old Tg2576 mice. Scale bar is 2 mm for A&B, 1 mm for C-F, 10 µm for G-I, 5 µm for J-K, L-P, 10 µm for Q & V, 20 µm for R-U & W-Z, 200 µm for AA&AB, 20 µm for AC, and 5 µm for AD.

The fraction of brain tissue occupied by Gallyas-positive staining in aging *APP*_swe_/*PS1*_ΔE9_ mice is shown in Fig. S1A. Conformationally-altered tau was detected with the MC-1 monoclonal antibody (Fig. S1B). Vascular and meningeal lesions were present in 18- and 24-month-old animals (Fig. S1C).

### [^18^F]Flortaucipir autoradiography

The paired helical filament (PHF) ligand [^18^F]Flortaucipir ([^18^F]AV-1451, [^18^F]T807) was used to quantify tau pathology in aging *APP*_swe_/*PS1*_ΔE9_ TG mice by autoradiography (Table 1). Increased binding was observed in the neocortex, hippocampus, amygdala and the cerebellum of 12-month-old *APP*_swe_/*PS1*_ΔE9_ mice, compared to age-matched WT, 3- and 6-month-old TG animals (*P*<0.001 for all regions; Bonferroni *post-hoc* tests). [^18^F]Flortaucipir binding was further elevated in the visual (*P*<0.001), somatosensory (*P*<0.001), motor cortex (*P*<0.001), and the amygdala (*P*<0.05) of 18- vs. 12-month-old *APP*_swe_/*PS1*_ΔE9_ TG mice. Increased binding over age-matched WT mice was first observed in the thalamus of TG animals at 18 months of age (*P*<0.001). In 24-month-old *APP*_swe_/*PS1*_ΔE9_ mice, [^18^F]Flortaucipir signal had increased in all brain regions examined compared to age-matched controls. Three-way ANOVA confirmed genotype-[F_(1,476)_=2603.1, *P*<0.001], age-[F_(4,476)_=457.3, *P*<0.001] and brain region-specific increases in the binding levels of [^18^F]Flortaucipir [F_(9,476)_=42.9, *P*<0.001], as well as significant age x genotype x region interaction effects [F_(36,476)_=5.5, *P*<0.001].

Within each brain area analyzed, there was a positive correlation between the age-dependent increase in the binding levels of [^18^F]Flortaucipir and the progressive increases in the density of Gallyas-positive lesions (Pearson r for all brain regions: 0.92, *P*<0.001; Fig. S2).

**Table 1.**
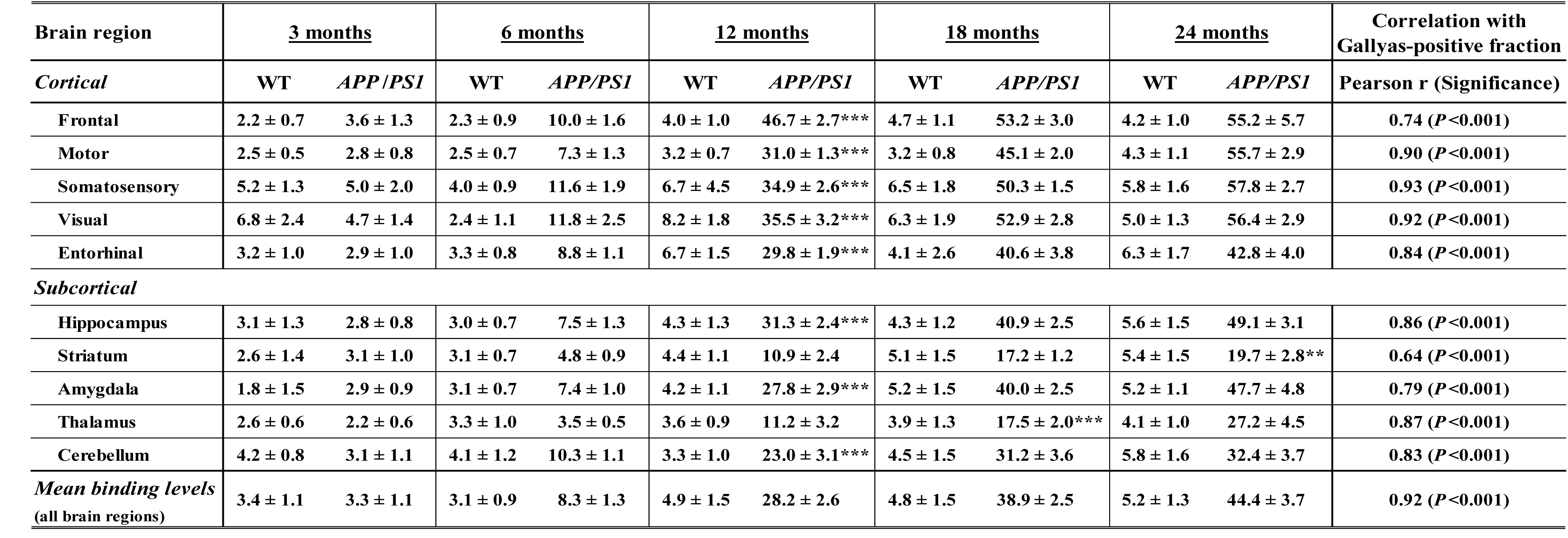
Autoradiography of [^18^F]Flortaucipir binding sites in aging *APP*swe/*PS1*ΔE9 mice. Fresh-frozen brain sections from *APP*_swe_/*PS1*_ΔE9_ and age-matched wild-type (WT) animals were incubated with 38.4±9.6 MBq [_18_F]Flortaucipir for a period of 60 min (specific activity: 145±68 GBq/µmol). Autoradiography data are presented as the mean specific binding of [_18_F]Flortaucipir (kBq/mL) ± standard error of the mean in brain regions of 5-6 animals/group. By 24 months of age, [^18^F]Flortaucipir binding in *APP*_swe_/*PS1*_ΔE9_ mice had increased across all brain areas examined vs. age-matched WT animals. The age-dependent increase in [_18_F]Flortaucipir binding levels was positively correlated with the progressive increase in Gallyas-positive argyrophilic signal, in all TG brain areas examined. \*\**P*<0.01, \*\*\**P*<0.001 vs. age-matched littermate control mice, Bonferroni *post-hoc* tests. Symbols of significant differences between groups of 24 & 18 vs. 3-, 6- and 12-month-old-mice were omitted from the table for clarity of presentation.

Representative autoradiograms of [^18^F]Flortaucipir binding sites are shown in Fig. 2. Binding was decreased in the presence of 50 µM unlabeled flortaucipir but was not reversed by 1 µM of the amyloid-preferring Pittsburgh compound B (PIB).

**Fig. 2.**
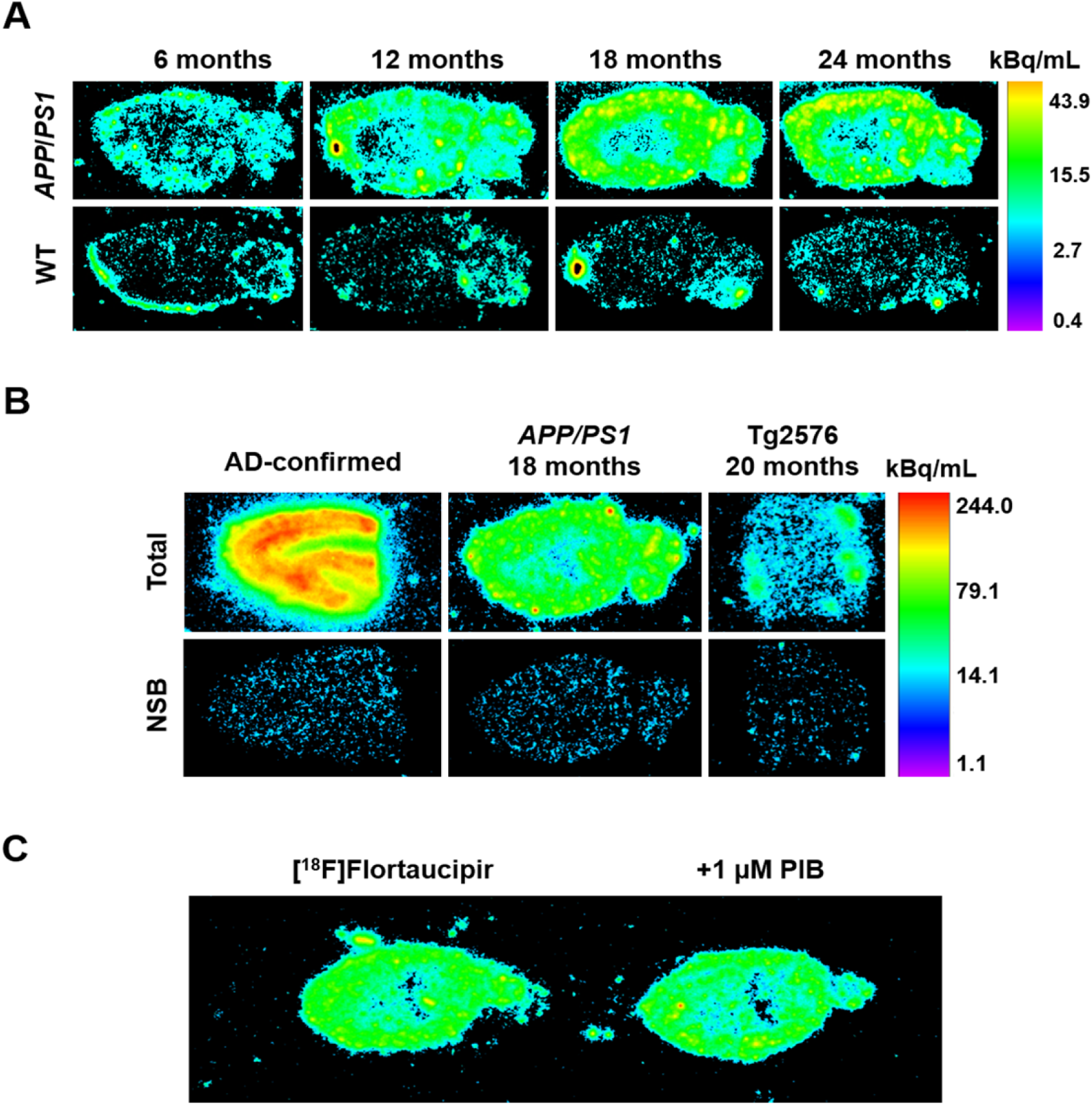
Representative autoradiograms of [18F]Flortaucipir binding sites. **(A)** Sagittal brain sections of aging transgenic (top panel) and wild-type mice (WT, lower panel), taken at the level of the entorhinal cortex [lateral 2.88±0.12 mm of the Paxinos and Franklin mouse atlas (74)]. Images were analyzed on a black & white display mode, and presented as a pseudocolor interpretation of black & white pixel intensity, calibrated in kBq/mL of [^18^F]Flortaucipir solution. Age-dependent increases in binding levels were observed exclusively in *APP*_swe_/*PS1*_ΔE9_ mice. **(B)** [_18_F]Flortaucipir binding in sections from the middle frontal gyrus of AD-confirmed patients, 18-month-old *APP*_swe_/*PS1*_ΔE9_ mice and 20-month-old Tg2576 animals, showing the magnitude of tau pathology in patient vs. transgenic mouse tissue. Non-specific binding (NSB) was assessed in the presence of 50 µM ‘cold’ flortaucipir. **(C)** Binding was not blocked by co-incubating sections with [_18_F]Flortaucipir and 1 µM of the amyloid-targeting agent Pittsburgh Compound B (PIB).

### Mapt expression

Relative expression of total *Mapt* mRNA was determined by RT-qPCR (Fig. 3). There were no age [F_(4,50)_=0.29, *P*>0.05], genotype [F_(1,50)_=0.93, *P*>0.05], or age x genotype interaction effects on the expression levels of *Mapt* [F_(4,50)_=1.21, *P*>0.05].

**Fig. 3.**
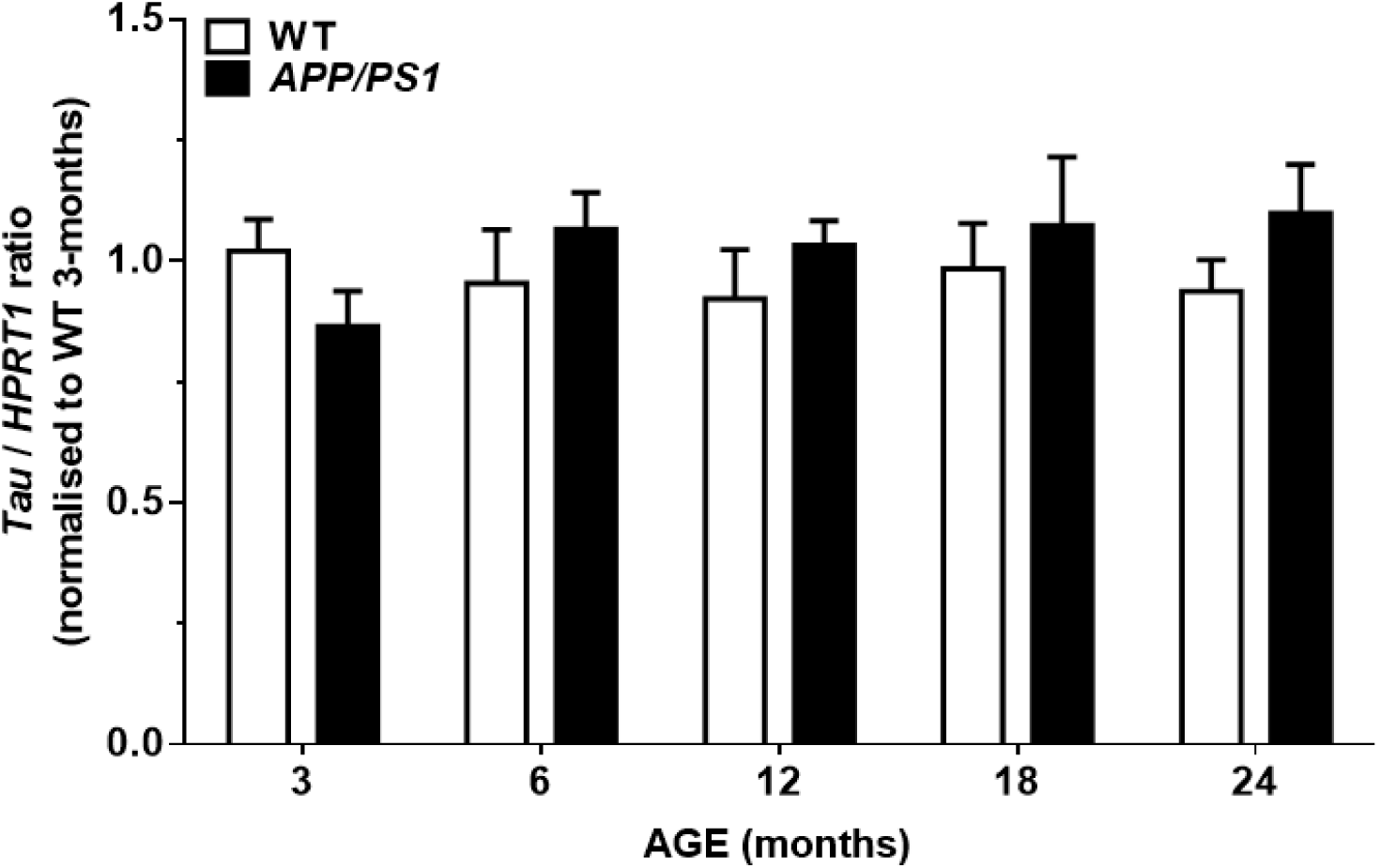
Regulation of *Mapt* mRNA in aging mice. Levels of endogenous murine *tau* mRNA were not altered by age or genotype. PCR products of x4 diluted *tau* cDNA were determined after 24 cycles. A single peak was obtained by melt-curve analysis, and no signal was detected in the genomic DNA and buffer controls. The efficiency of amplification was 99.2±0.2% for *Hprt1* and 100.3±2.1% for *Mapt*.

### Isolation and Transmission Electron Microscopy (TEM) of sarkosyl-insoluble tau

The general methods of Sahara et al. (*20*) and Greenberg and Davies (*21*) were evaluated for the extraction of PHFs from the 24-month-old *APP*_swe_/*PS1*_ΔE9_ TG brain (Fig. S3). Although longer filaments were isolated by the procedure of Sahara et al., the Greenberg and Davies method was chosen for the isolation of sarkosyl-insoluble tau from 3- and 24-month-old mice, based on immunoblotting experiments, solubility considerations, and to allow for comparisons with literature data (*22*). Soluble and insoluble tau levels were measured in mouse brain homogenates by using the mouse Total Tau Meso Scale kit (Meso Scale Diagnostics LLC). TEM was used to visualize filaments in the sarkosyl-insoluble extracts from the TG mouse and AD patient brains by negative staining.

Tau protein levels increased with age in the pellet obtained by centrifuging WT and *APP*_swe_/*PS1*_ΔE9_ homogenates at 27,000 × g [Fig. 4A; age effect: F_(1,18)_=50.0, *P*<0.001; genotype effect: F_(1,18)_=2.4, *P*>0.05]. Levels of tau in the supernatant fraction were not different between 3- and 24-month old, WT and *APP*_swe_/*PS1*_ΔE9_ TG animals [age: F_(1,16)_=0.6, *P*>0.05; genotype: F_(1,16)_=0.0, *P*>0.05]. Treatment of the supernatant with 1% sarkosyl for 2 h at 37°C increased the concentration of tau in the detergent-soluble fraction by >3-fold. Sarkosyl-soluble tau levels were lower in the 24- vs. 3-month-old mouse brain [F_(1,16)_=12.5, *P*<0.01], irrespective of genotype [F_(1,16)_=0.5, *P*>0.05]. Sarkosyl-insoluble tau was not detected in 3-month-old animals, and its levels were not different between 24-month-old *APP*_swe_/*PS1*_ΔE9_ and WT mice [t_(8)_=0.7, *P*>0.05; independent two-tailed Student’s t-test].

**Fig. 4.**
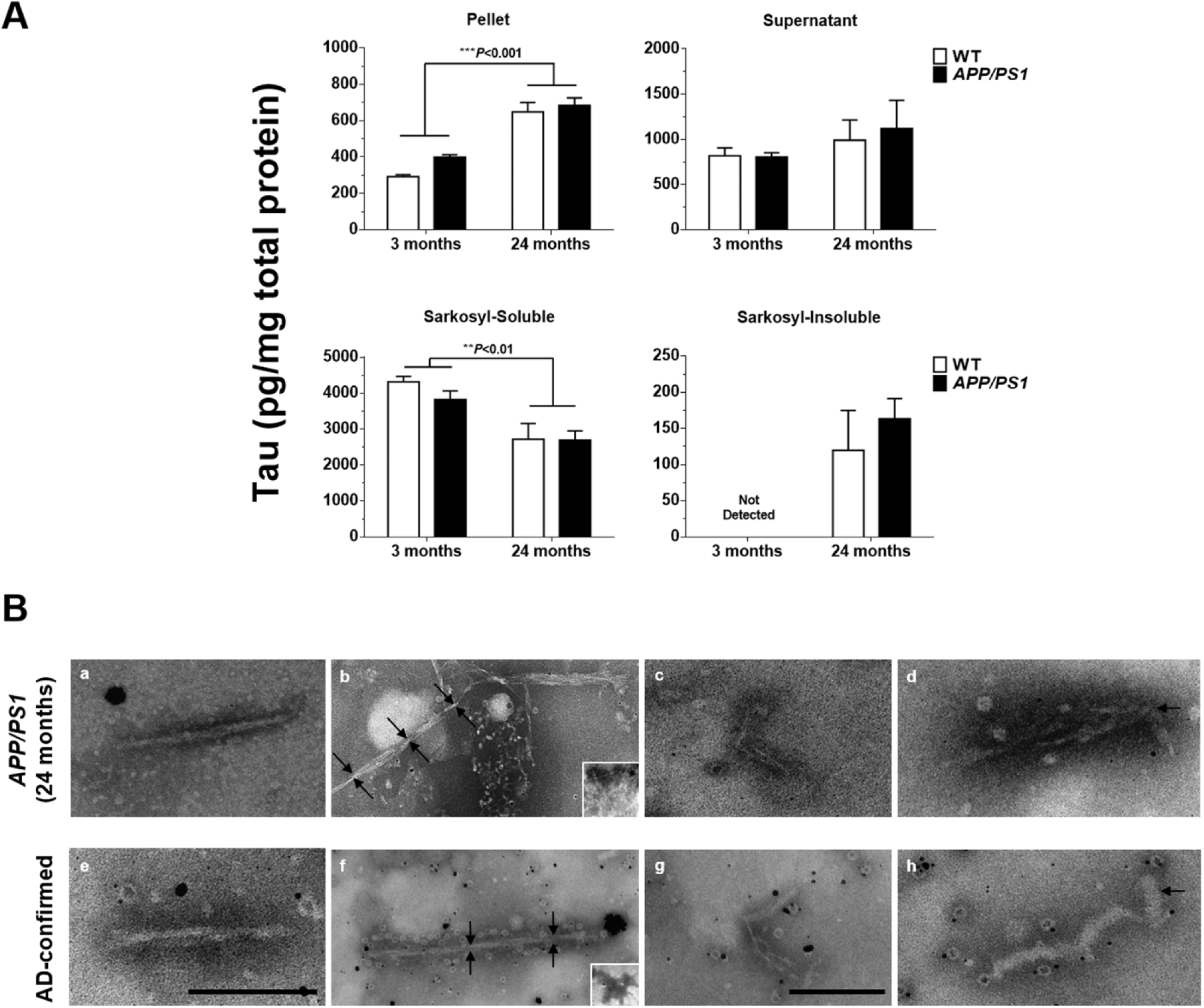
Isolation and electron microscopy of sarkosyl-insoluble tau. **(A)** Levels of soluble and insoluble tau were determined with the mouse Total Tau Meso Scale kit. Tau levels increased with age in the pellet obtained by centrifuging brain homogenates at 27,000 × g. The resulting supernatant was treated with 1% sarkosyl and centrifuged at 200,000 × g. The solubility of tau in sarkosyl was decreased with age, irrespective of genotype. **(B)** Overview of negatively-stained filament types in the sarkosyl-insoluble fraction from 24-month-old *APP*_swe_/*PS1*_ΔE9_ and AD brain tissue. Fibrils of ∼20 nm in width, appearing as straight filaments (a) or as two intertwined fibrils (e), each with a diameter of ∼10 nm. PHFs with axial periodicities of ∼80 nm (b & f; arrows) were present in *APP*_swe_/*PS1*_ΔE9_ mice, and more frequently observed in AD patient material. The inserts show ‘stacked’ PHFs, which were denser in the AD preparation. Structures commonly identified in the detergent-insoluble fractions of the mouse and human brain included bent fibrils of ∼7 nm in width (c & g), and rod-shaped particles (d & h; arrows). Scale bars: 200 nm (a, b, e, f), 100 nm (c, d, g, h).

Fig. 4B shows negatively-stained filaments in the sarkosyl-insoluble extract from the 24-month-old *APP*_swe_/*PS1*_ΔE9_ mouse and AD patient brain. Fibrils of mean length 104.9±8.3 nm and width 10.1±0.5 nm were isolated from TG mice. Wider fibrils (∼20 nm), with or without a pronounced twist, were readily detected (a & e). Longer filaments (271.7±11.3 nm), with axial periodicities of 78.7±9.8 nm, constituted ∼8% and ∼34% of the fibril population analyzed in the *APP*_swe_/*PS1*_ΔE9_ and AD brains, respectively (b & f). Clusters of long filaments, which were denser in AD patient material, were present in the insoluble preparation from *APP*_swe_/*PS1*_ΔE9_ mice (b & f, inserts). Thin, bent fibrils (c & g) and rod-shaped particles (d & h) were observed in both 24-month-old *APP*_swe_/*PS1*_ΔE9_ and AD brains. There were no between-species differences in the dimensions of the isolated filaments [short filaments, length: t_(82)=_0.1, *P*>0.05, width: t_(82)_=1.2, *P*>0.05; long filaments, length: t_(16)=_0.3, *P*>0.05, width: t_(16)=_0.8, *P*>0.05; independent two-tailed t-tests].

### Proteomics of sarkosyl-insoluble tau

The sarkosyl-insoluble fractions extracted from 3- and 24-month-old mouse brain, AD and non-AD individuals, were pooled and digested with trypsin & Lys-C. The peptides were labeled with Tandem Mass Tags (TMT), fractionated, and analyzed by nanoLiquid Chromatography-Electrospray Ionization Mass Spectrometry (LC-ESI MS/MS).

A list of tau-associated proteins quantified in the sarkosyl-insoluble proteome is shown in Table 2. Lists of between-group abundance ratios for all regulated proteins are shown in Data File S1. There were 583 proteins identified in the sarkosyl-insoluble mouse proteome, of which 456 were also present in the human samples. Isoforms of tau with three (3R) and four (4R) microtubule-binding repeats were extracted from both human and the murine brain. In mice, all isoforms collapsed under the term microtubule associated protein (MAP; UniProt accession number: B1AQW2). Mouse MAP was regulated by age, rather than genotype. The protein was enriched 2.1-fold in 24- vs. 3-month-old TG mice, and 1.8-fold in 24- vs. 3-month-old WT mice. Genotype-specific enrichment was observed for mouse tau isoform-B (UniProt accession number: P10637-3), a 3R isoform of tau with an extended C-terminal domain, which was identified by the sequence ^205^KVQIVYKPVDLSKV^218^. Tau isoform-B increased 3.2-fold in 24-month-old TG vs. WT mice, and 4.5-fold in 24- vs. 3-month-old TG animals. Human MAP (UniProt accession number: A0A0G2JMX7), containing tau isoforms P10637-2, -4, -6 & -8, was 37-fold enriched in the sarkosyl-insoluble fraction of AD compared to non-AD brain.

**Table 2.**
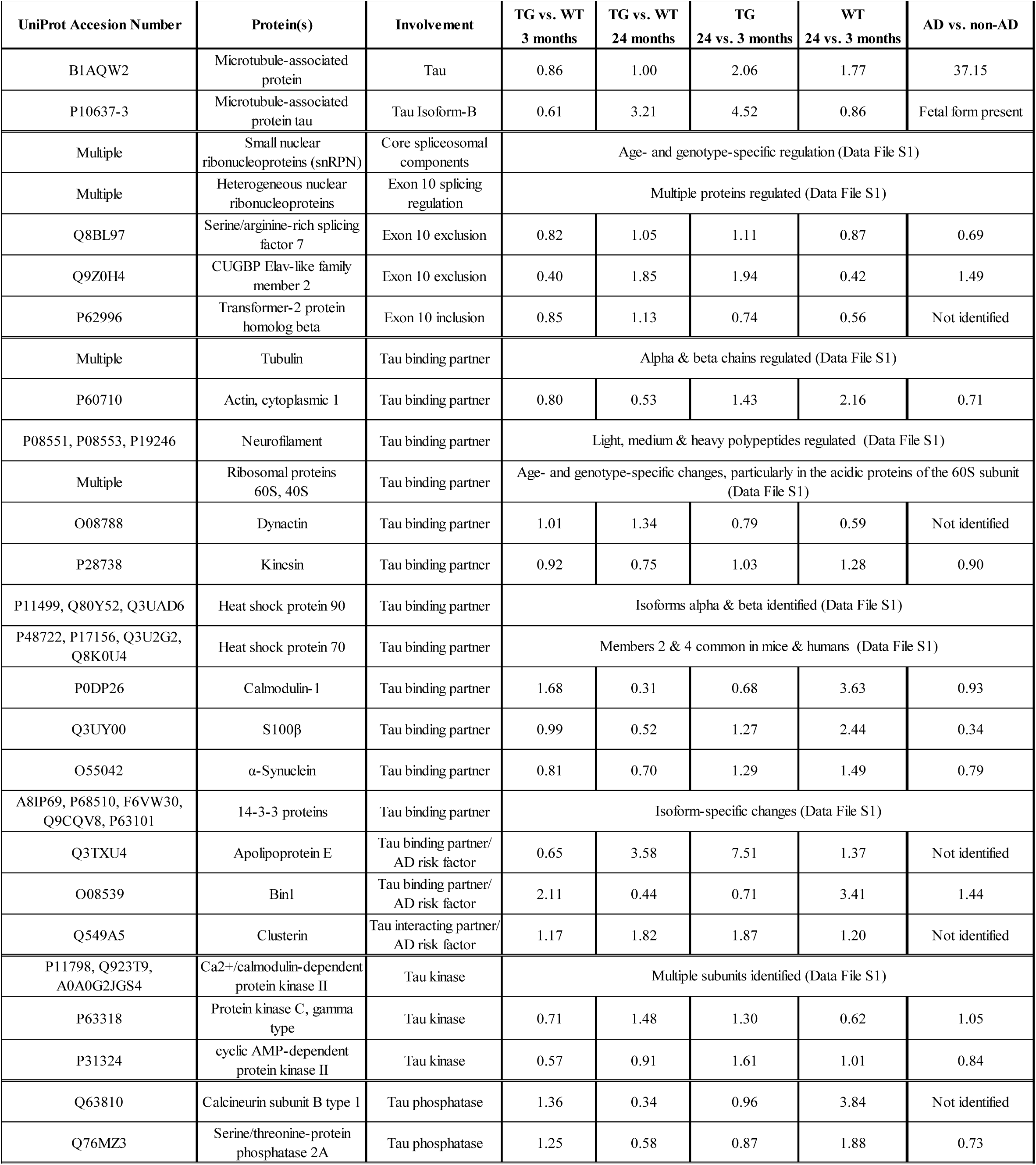
Proteomics of sarkosyl-insoluble tau. Tau-associated proteins quantified in the detergent-insoluble fractions of the mouse and human brain. The presented proteins have been selected for their documented roles in the regulation and binding of tau.

The mouse MAP sequence ^174^KVAVVR**<u>T</u>**PPKSPSA**<u>S</u>**KS^190^, phosphorylated at Threonine (T) 180 and Serine (S) 188, was more than 20-fold enriched in 24-month-old TG, compared to age-matched WT and 3-month-old *APP*_swe_/*PS1*_ΔE9_ mice. The peptide sequence was not regulated in aging WT animals. An orthologous sequence of the human MAP was phosphorylated at Threonine (T) 566 and Serine (S) 573 (^560^KVAVVR**<u>T</u>**PPKSPS**<u>S</u>**AKS^576^). The reported phosphorylation sites correspond to amino acids (aa) T231 and S238 of the human tau isoform with 441 aa. Indications of additional phosphorylation sites were obtained by searching modified peptides against tau isoform- and species-specific databases. Phosphorylated S396, S400 and S404 on the conserved sequence ^396^**<u>S</u>**PVV**<u>S</u>**GDT**<u>S</u>**PR^406^ of the human 441 aa isoform were identified in the sarkosyl-insoluble mouse proteome. Phosphorylation at S404 in 24-month-old TG mice was confirmed by immunoblotting (Fig. S4). In addition to phosphorylation, murine MAP was deamidated at Asparagine (N) 44, a site on the N-terminal domain of tau that is not conserved in humans (^34^AEEAGIGDTP**<u>N</u>**QEDQAAGHVTQAR^57^). Human MAP was sdeamidated at position N484, corresponding to N167 of the 441 aa tau isoform (^473^GAAPPGQKGQA**<u>N</u>**ATRIPAK^491^).

The database for annotation, visualization and integrated discovery (DAVID, v6.8) was used for gene ontology (GO) enrichment analysis of the sarkosyl-insoluble proteome (*23, 24*). RNA splicing, mRNA processing and translation were among the 10 most enriched biological processes associated with protein upregulation in 24-month-old *APP*_swe_/*PS1*_ΔE9_ vs. WT mice and AD vs. non-AD subjects. Ribonucleoprotein complexes, ribosomes, and exosomes were among the 10 most enriched cellular components in the insoluble extracts from the mouse and human brain (Fig. 5A). The top 10 molecular functions of the enriched proteins were associated with poly(A) RNA binding, as well as binding of molecules contributing to the structural integrity of ribosomes and the cytoskeleton (Fig. 5B). Pathway-based enrichment analysis of upregulated proteins in 24-month-old *APP*_swe_/*PS1*_ΔE9_ vs. WT mice involved GO terms such as Alzheimer’s and Huntington’s disease, long-term depression, cholinergic, serotonergic and glutamatergic synapse (Fig. 5C). Glycolysis/gluconeogenesis and the Krebs cycle were among the top 10 pathways for downregulated proteins (Fig. 5D).

**Fig. 5.**
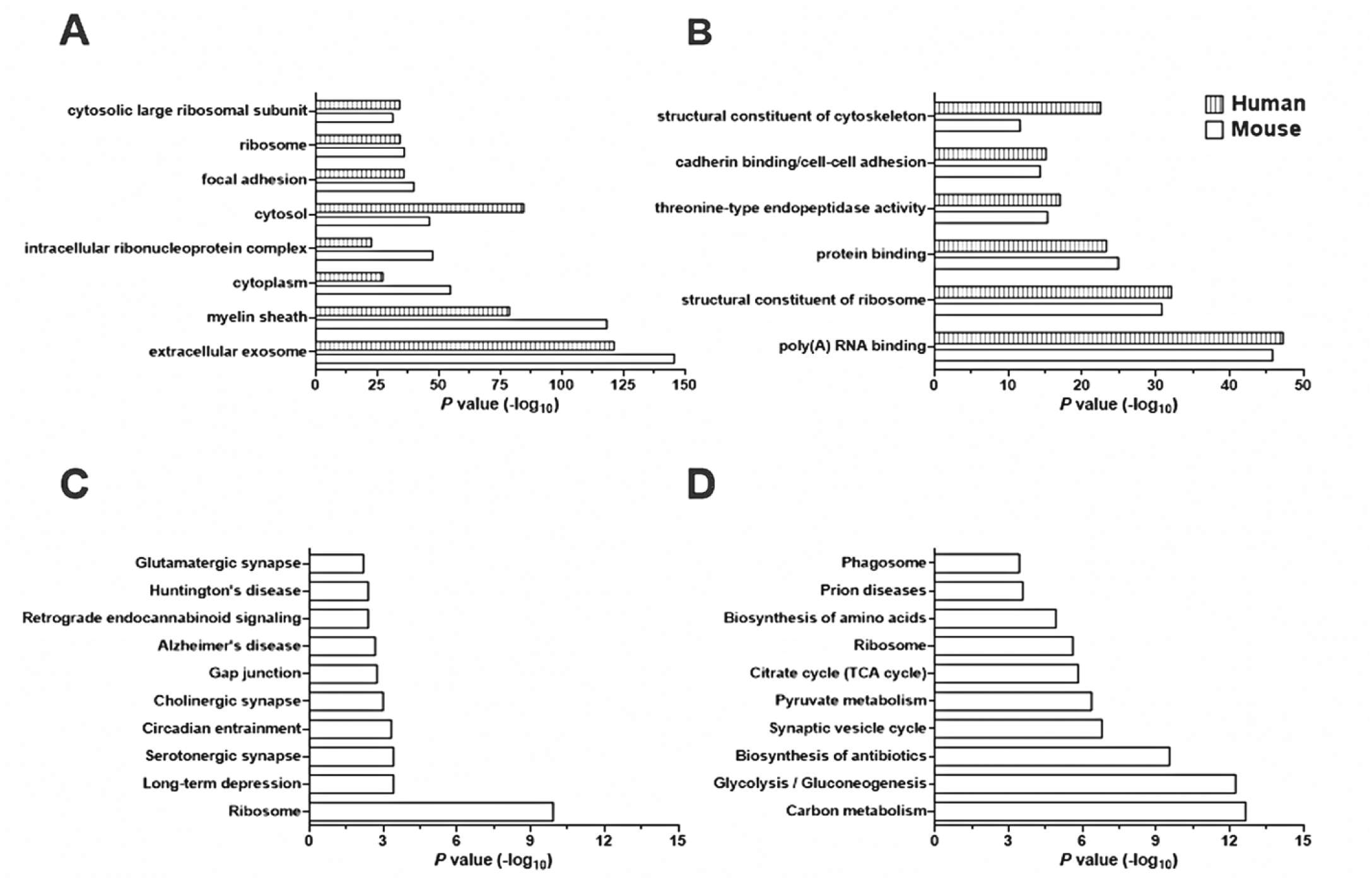
Gene Ontology (GO) enrichment analysis of the sarkosyl-insoluble proteome. **(A)** Enriched cellular components; **(B)** Enriched molecular functions; **(C)** Top 10 enriched pathways based on protein upregulation in 24-month-old TG vs. WT mice, according to the Kyoto Encyclopedia of Genes and Genomes (KEGG); **(D)** Top 10 enriched KEGG pathways based on protein downregulation in 24-month-old TG vs. WT mice. Functional annotation clustering was generated by using DAVID software. Maximum enrichment probability (*P* value) was based on an EASE score threshold value of 0.05.

## Discussion

The present study describes tauopathy in murine models of familial AD. Neurofibrillary alterations in *APP*_*swe*_/*PS1*_*ΔE9*_ and Tg2576 mice were demonstrated by a set of tools that are currently used for the evaluation of pathological tau clinically, such as the Gallyas silver stain and [^18^F]Flortaucipir. The presence of PHF tau was confirmed by TEM of sarkosyl-insoluble preparations from the *APP*_*swe*_/*PS1*_*ΔE9*_ mouse brain. As murine tau possesses a remarkably high number of 76 potential serine/threonine and 4 tyrosine phosphorylation sites, an antibody-free proteomics approach was used for the detection of tauopathy-related epitopes. Of the five hyperphosphorylated sites identified, S404 has been associated with the intraneuronal and extracellular deposition of NFTs in AD (*25*). The pathology observed in the present study occurred at physiological levels of endogenous tau, as there was no difference in total tau mRNA and protein between *APP*_*swe*_/*PS1*_*ΔE9*_ and WT mice. Hence, in addition to progressive amyloidosis (*3*), neuroinflammation (*4*) and neurodegeneration (*5*), *APP*_*swe*_/*PS1*_*ΔE9*_ mice develop progressive neurofibrillary pathology of the AD type, mimicking a range of AD pathologies, in a translationally-relevant manner. The observation that endogenous tau accumulates secondarily to Aβ in models of cerebral amyloidosis is entirely consistent with *post-mortem* (*26*) and *in vivo* imaging data (*27*), showing that the development of cortical tau pathology in AD patients is associated with, and may depend on, pre-existing amyloid pathology.

Current approaches to induce tauopathy in mice have been criticized for generating models that poorly recapitulate the situation in the AD brain, as *TAU* in AD is neither overexpressed, nor mutated (*28*). FLTD-linked mutations, in particular, induce tauopathy that is not only morphologically different than that of AD (e.g. Pick bodies), but further characterized by distinct neurodegenerative processes. For example, cholinergic neurons are extensively lost in AD, but not in FTLD (*29*). Acetylcholinesterase inhibitors, which are prescribed for the symptomatic relief of cognitive impairment in AD, are largely ineffective in FTLD and may even worsen its symptoms (*30*). Thus, the pathophysiology that differentiates AD from primary tauopathies is unlikely to be modeled in mutant *TAU* models. Moreover, neurofibrillary alterations in *TAU* overexpressing mice occur in the absence of Aβ deposition, which is a defining feature of AD histopathology. The present results indicate that amyloidosis models may overcome these limitations, by reproducing both the neurofibrillary pathology of familial AD and the molecular heterogeneity that is associated with it. In addition to the spontaneous aggregation of tau in *APP*_*swe*_/*PS1*_*ΔE9*_ and Tg2576 mice, analysis of the sarkosyl-insoluble *APP*_*swe*_/*PS1*_*ΔE9*_ proteome identified proteins that have been strongly linked to AD pathogenesis, in general, and tau pathology in particular. Among them, APOE and BIN1 are encoded by genes whose variants are known to increase the risk of late-onset AD, through pathways involving interactions with both APP (*31, 32*) and tau (*33-35*). Core components of the spliceosome, on the other hand, particularly Sm-D1 and Sm-D2, are closely related to the deposition of NFTs, but not plaques in familial AD (*36*). This literature implicates multiple mechanisms in AD tauopathy, which occur downstream of Aβ processing in cases of autosomal dominant AD (ADAD) and, as shown here, *APP*_*swe*_/*PS1*_*ΔE9*_ mice.

Although the sporadic and familial forms of AD share common clinical and histopathological features, it is becoming increasingly recognized that they are not precisely equivalent (*37*). Positron emission tomography (PET) with [^11^C]PIB demonstrates accumulation of Aβ in the cerebellum of familial AD cases, which is not typical of sporadic AD (*38*). Cerebellar deposition of hyperphosphorylated tau has been observed in ADAD cases harboring the *PSEN1* E280A mutation, but not in sporadic AD (*39*). Thus, the pronounced cerebellar involvement in *APP*_*swe*_/*PS1*_*ΔE9*_ mice, which are known to accumulate Aβ in this region (*40*), suggests that the model mimics familial, rather than the sporadic forms of AD. Reports of cerebellar pathology in ADAD cases and *APP*_*swe*_/*PS1*_*ΔE9*_ mice warrant caution in using the cerebellum as a reference region for the quantification of [^11^C]PIB and [^18^F]Flortaucipir PET (*41, 42*), as this is likely to underestimate cortical Aβ and tau pathology, respectively.

Unlike the imbalance in Aβ homeostasis, which is thought to be central in the pathogenesis of AD, gross changes in tau production and clearance were not observed in this study. On the one hand, Gallyas-, [^18^]Flortaucipir- and thioflavin-S-positive signal was observed in the vasculature of 18-24-month-old *APP*_*swe*_/*PS1*_*ΔE9*_ mice. Moreover, there were age- and genotype-specific changes in multiple components of the phagosome and proteasome in TG vs. WT animals (Data File S1). On the other hand, total tau mRNA and protein levels were not different between 24-month-old WT and TG mice, as evidenced by tau mesoscale, proteomics and PCR. Moreover, age-dependent decreases in the solubility of tau were equally observed in *APP*_*swe*_/*PS1*_*ΔE9*_ and control animals. Notwithstanding that the contribution of individual pathways to tau degradation was not assessed in this study, these findings suggest that the neurofibrillary alterations observed in *APP*_*swe*_/*PS1*_*ΔE9*_ mice are not mediated by an imbalance between the production and clearance of tau. It is important to note that, unlike in TG mice, sarkosyl-insoluble tau was increased in AD vs. non-AD tissue, a finding that is consistent with literature data on the regulation of human tau in AD (*43*). It might be that the increased concentration of brain tau in late-stage AD is associated with heavily impaired clearance pathways or pronounced neuronal damage, processes that may not be modeled in 24-month-old *APP*_*swe*_/*PS1*_*ΔE9*_ mice. Alternatively, the present data may highlight the involvement of transcriptional and translational mechanisms, rather than production and clearance pathways, in the assembly of PHF tau.

A prevalence of 3R isoforms in the composition of NFTs has been observed in the AD hippocampus by immunohistochemical and biochemical methods (*44*). Moreover, a shift from 4R to 3R isoforms has been associated with the morphological evolution of tau-positive neurons from a pre-tangle to the NFT state (*45*). Although the literature on the regulation of tau isoforms in AD remains scarce, the present results support the notion that an imbalance in tau isoform ratio is involved in the neurofibrillary alterations of AD, with 3R isoforms being preferentially sequestered into the insoluble tau fraction. The identification of tau isoform-B, a 3R isoform that is predominantly expressed in the fetal mouse brain, supports the suggestion that immature tau isoforms participate in AD tauopathy (*46*), and implicates aberrant transcription and translation mechanisms in the disease process. A re-induction of fetal tau may be attributed to the deregulation of core splicing machinery, which was marked in this study and considered to occur early and selectively in AD (*47*). Moreover, as the selection of splice sites is determined by canonical sequences encoded into the genome, the re-expression of fetal isoforms might be a consequence of aberrant DNA replication during cell cycle re-entry (*48*). Cell cycle proteins that were deregulated in an age- and genotype-specific manner in this study include Sub1, cdc42, CEND1, Histone H3 and nucleolin (Data File S1). Clearly, the exact mechanisms underlying tauopathy in AD cannot be resolved by the present set of experiments. The data demonstrate, however, that the formation of PHF tau is associated with loss of regulatory control over *tau* splicing *in vivo*, which may have important implications for the origins and management of tauopathy in AD. It is tempting to speculate that tau hyperphosphorylation may partly be due to the re-emergence of fetal isoforms, which are known to be over-phosphorylated compared to adult tau (*49*). Moreover, it is plausible that an imbalance in tau isoform ratio mediates protein mislocalization from the axonal to the somatodendritic compartment, as distinct tau isoforms are differentially sorted across the cell (*50*). Of note, cofilin-dependent, ‘classical’ pathways of tau missorting (*51*) may also be involved in the pathology observed in this study, as cofilin was reduced in the sarkosyl-insoluble proteome of 24-month-old *APP*_*swe*_/*PS1*_*ΔE9*_ mice. Collectively, these data highlight the relevance of amyloidosis models for studying the diverse macroscopic and molecular aspects of AD tauopathy.

The limitations associated with models overexpressing *APP* and *PSEN* mutations have been discussed previously (*2*). To exclude the possibility that tauopathy is an artefact of *APP* or *PSEN* overexpression, it would be important to determine whether it develops in second-generation amyloidosis models, carrying AD-related mutations in endogenous genes. Moreover, as there is evidence of [^18^F]Flortaucipir binding to monoamine oxidases (MAO; *52*), signal quantification in the presence of MAO inhibitors is warranted to determine the extent of off-target binding, if any (*53*). Practical considerations in using amyloidosis mice to study tauopathy include long waiting times for the accumulation of endogenous murine tau, mouse-on-mouse antibody issues, and the low abundance of pathology as compared to human AD. While ∼30% of all Nissl-positive cells in the prefrontal cortex of Braak stage V-VI brains may contain NFTs (*54*), Gallyas-positive signal in this study occupied ∼1% of the frontal cortex of 24-month-old *APP*_*swe*_/*PS1*_*ΔE9*_ mice, neuritic structures included. Nevertheless, cognitive impairment in AD is known to correlate with the spread of tau pathology, and the number of brain areas containing at least one NFT has been shown to be the best explanatory variable of intellectual status in AD (*55*). In this context, it is worth evaluating whether measurements of tauopathy in aging *APP*_*swe*_/*PS1*_*ΔE9*_ mice correlate with the progressive cognitive impairment that these animals exhibit in the Barnes maze assay (*56*).

## Materials and Methods

### Study design

Mice were grouped according to age and genotype. Sample numbers were based on preliminary studies, showing absence of Gallyas and [^18^F]Flortaucipir signal in 18-month-old WT vs. TG animals. (Immuno)histochemistry, autoradiography, the isolation of sarkosyl-insoluble tau, and electron microscopy studies were repeated at least three independent times. Tau Meso Scale was performed two independent times. Remaining samples were pooled and subjected to proteomics. To avoid cross-contamination during the isolation of sarkosyl-insoluble tau, glassware was washed in ultrapure de-ionized H_2_O (dH_2_O, Ultra Clear™, Siemens), followed by rinses in formic acid (FA, 98-100%; Merck Millipore), dH_2_O, ethanol (99%; VWR International) and dH_2_O. No samples were excluded from data analysis, which was performed in an unblinded manner. To compare tau pathology in transgenic mouse vs. human brain, tissue from the middle frontal gyrus of an AD-confirmed patient [BB08-002, Female, 80 years old, *post-mortem* interval (PMI): 9 h] and a non-AD subject (BB16-023, Female, 83 years old, PMI unknown) were processed along with the murine samples for the Gallyas silver stain, autoradiography, electron microscopy and proteomics experiments. The AD and non-AD samples were chosen for their abundance and complete lack of tau pathology respectively, as assessed by Gallyas silver staining and [^18^F]Flortaucipir autoradiography. To avoid confounding effects of anesthesia on tau phosphorylation, mice were euthanized by cervical dislocation.

### Ethical statement

Mouse tissue: All procedures complied with Danish law (Dyreværnsloven-Protection of Animals Act, nr 344/2005) and European Union directive 2010/63/EU, regulating animal research. Ethical permission was granted by the Animal Ethics Inspectorate of Denmark (nr 2011/561-1950).

Human tissue: Fresh-frozen samples from the middle frontal gyrus were obtained from the Maritime Brain Tissue Bank, Department of Medical Neuroscience, Faculty of Medicine, Dalhousie University, Sir Charles Tupper Building, 5850 College Street, Halifax Nova Scotia B3H 4R2. Ethical approval was obtained from the Nova Scotia Health Authority Research Board in Halifax, Canada, and the Danish Biomedical Research Ethical Committee of the region of Southern Denmark (Project ID: S-20070047). Informed, written consent forms were obtained for all subjects.

### Animals and tissue sectioning

*APP*_swe_/*PS1*_ΔE9_ mice (*57*), originally purchased from the Jackson Laboratories (MMRRC Stock No: 34832-JAX), were bred and maintained on a C57BL/6J background. The animals were group-housed (4-8/cage) in a temperature (21±1°C) and humidity controlled environment (45-65%), under a 12:12 h light:dark cycle (lights on: 7 am). Food and water were available *ad libitum*. Female *APP*_swe_/*PS1*_ΔE9_ mice were used at 3, 6, 12, and 18 months of age. Sex- and age-matched WT littermates were used as control. Both male and female mice were used in the 24-month-old groups (n=6/genotype & age-group, total animal number: 60). The animals were euthanized by cervical dislocation, and brains immediately removed and bisected along the midline. Right hemispheres were frozen in isopentane on dry-ice (−30°C). The olfactory bulb, striatum, cortex, hippocampus, diencephalon, brainstem and cerebellum from the left hemisphere were dissected on a petri dish on ice, collected in Eppendorf^®^ tubes, and frozen on dry-ice. The tissue was stored at −80°C until use.

Sectioning was carried out at −17°C using a Leica CM3050S cryostat (Leica Biosystems GmbH). Series of 20 µm-thick sagittal sections were collected at 300 µm intervals. The sections were mounted onto ice-cold Superfrost™. Plus slides (Thermo Fisher Scientific), dried at 4°C in a box containing silica gel for at least 2 h, and stored at −80°C for future experiments. Every 13^th^ and 14^th^ section was collected in Eppendorf^®^ tubes for RNA extraction with Trizol™.

Fresh-frozen coronal brain sections of male and female, 20-month-old Tg2576 and WT mice were provided by the Centre for Biological Sciences, University of Southampton, U.K.

### (Immuno)histochemistry, autoradiography and proteomics

The Gallyas silver stain was performed according to Kuninaka et al. (*58*), thioflavin-S according to Sun et al. (*59*), [^18^F]Flortaucipir autoradiography according to Marquié et al. (*60*), proteomics according to Kempf et al. (*61*). Protocol details are provided in Supplementary Materials and Methods.

### Statistical analysis

Parametric testing was employed following inspection of the data for normality with the Kolmogorov-Smirnov test in Prism (v6.01; GraphPad Software). Data sets were analyzed by Statistica ™ v10 (TIBCO Software Inc., USA). The effects of age, genotype and brain region on the binding levels of [^18^F]Flortaucipir were analyzed by three-way ANOVA. Gallyas-positive area fraction and tau gene/protein levels were analyzed by two-way ANOVA for the independent factors age and brain region or genotype, respectively. Where ANOVA yielded significant effects, Bonferroni *post-hoc* comparisons were used to detect between-group regional and age-dependent differences. Levels of sarkosyl-insoluble tau between 24-month-old TG and WT mice, and PHF dimensions extracted from TG vs. AD brain were compared by two-tailed independent Student’s t-tests. Significance was set at α=0.05. A 1.3-fold change cut-off value for all TMT ratios was used to rank proteins as up- or down-regulated in the proteomics study (*62*).

## Acknowledgments

We thank Andrew Reid, senior technician and manager of the Maritime Brain Tissue Bank, for organizing the transportation of human tissue. We acknowledge the Core Facility for Integrated Microscopy, Faculty of Health and Medical Sciences, University of Copenhagen, for assisting with TEM. The Villum Center for Bioanalytical Sciences at SDU is acknowledged for supporting the proteomics part of the study. Precursor material for the synthesis of [^18^F]Flortaucipir was generously supplied by AVID Radiopharmaceuticals, Philadelphia, PA, USA.

## Author contributions

AM designed the project and wrote the manuscript, performed (immuno)histochemistry, autoradiography, tau isolation studies, and assisted with TEM. CT and SJK performed proteomics and immunoblotting. MA and RV performed tissue sectioning, (immuno)histochemistry, autoradiography and tau filament analysis. SP performed PCR and tau Meso Scale. AML organized and performed autoradiography. HA synthesized [^18^F]Flortaucipir. JLT performed (immuno)histochemistry and provided Tg2576 tissue, SD provided human tissues and critically reviewed the manuscript. DJB, MRL and BF supervised the project and contributed to its design. All authors assisted with data interpretation, participated in drafting the manuscript and approved its final version.

## Data and materials availability

The proteomics data have been deposited to the ProteomeXchange Consortium (*63*) via the PRIDE partner repository with the dataset identifier PXD009306 [username; password: HdJxxVxU (*64*)]. The MC-1 antibody, and material for the synthesis of [^18^F]Flortaucipir were obtained through an MTA.

## Supplementary Materials

### Materials and Methods

#### Gallyas silver staining

The Kuninaka et al. simplification of the modified Gallyas method was used to examine neurofibrillary pathology (*58, 65*). Fresh-frozen sections were directly immersed in 4% neutral buffered formalin (NBF) at 4°C for 24 h. The sections were thoroughly washed in ultrapure deionized H_2_O (dH_2_O; Ultra Clear™, Siemens), dried at room temperature (RT) for 10 min and defatted for 1 h in a solution of chloroform/99% ethanol (1:1) in the dark. Following hydration through a series of graded ethanols into dH_2_O (2 × 1 min: 99%, 96%, 70%), slides were immersed into an aqueous solution of 0.25% potassium permanganate (20 min), washed in dH_2_O (1 min), and incubated in 1% oxalic acid (2 min). After washing in dH_2_O (2 × 5 min), sections were transferred into the alkaline silver iodide solution (1 min), washed with 0.5% acetic acid (2 × 5 min), and developed in a water bath at 15°C, until the appearance of a brownish shade (12-14 min). The developed sections were washed in 0.5% acetic acid (3 min), toned with 0.1% gold chloride (10 min), fixed with 1% sodium thiosulfate (5 min), and counterstained with 0.1% nuclear fast red (3 min). Ethanol and chloroform were from VWR International. The remaining chemicals were from Sigma-Aldrich Co.

#### Aβ immunohistochemistry on silver-stained sections

The biotinylated 6E10 antibody (SIG-39340, Nordic BioSite) was used to investigate the association between neurofibrillary pathology and Aβ load in *APP*_swe_/*PS1*_ΔE9_ mice. Clone 6E10 is raised against amino acids 1-16 of human Aβ, recognizing multiple amyloid peptides and precursor forms (manufacturer information). Silver-stained sections were immersed in 70% formic acid for 30 min, rinsed for 10 min in 50 mM Tris-buffered saline (TBS, pH 7.4), and further washed/permeabilised in TBS containing 1% Triton X-100 (3 × 15 min). Sections were subsequently blocked for 30 min in TBS, containing 10% fetal bovine serum (FBS). Incubation with the 6E10 anti-β-amyloid antibody was carried out overnight at 4°C, in TBS containing 10% FBS (1:500 dilution of stock). Adjacent, negative control sections were incubated with biotin-labeled mouse IgG1 (MG115, Thermo Fisher Scientific), diluted to the same protein concentration as the primary antibody. Following incubation with 6E10, the sections were adjusted to RT for 30 min and washed in TBS+1% Triton X-100 (3 × 15 min). Endogenous peroxidase activity was quenched for 20 min in a solution of TBS/methanol/H_2_O_2_ (8:1:1). After washing in TBS+1% Triton X-100 (3 × 15 min), sections were incubated for 3 h with HRP-streptavidin in TBS/10% FBS (1:200; GE Healthcare Life Sciences). After a final wash in TBS (3 × 10 min), peroxidase activity was visualised with 0.05% 3,3′diaminobenzidine (DAB) in TBS buffer, containing 0.01% H_2_O_2_ (Sigma Aldrich Co.).

#### MC-1 immunohistochemistry

Fresh-frozen sections were directly immersed in 4% NBF at 4°C for 24 h. The sections were thoroughly washed in dH_2_O, dried at RT for 10 min, and defatted for 1 h in a solution of chloroform/99% ethanol (1:1) in the dark. Following hydration through a series of graded ethanols into dH_2_O (2 × 1 min: 99%, 96%, 70%), endogenous peroxidase activity was quenched for 30 min with 1.5% H_2_O_2_ in TBS. The sections were washed and blocked for 1 h at RT using 10% FBS in 0.05% Triton X-100 (TBS-Tx; blocking buffer). The mouse anti-human MC-1 antibody, a generous gift from Dr. Peter Davies, was diluted in blocking buffer (1:10 dilution of stock). Negative control sections were incubated in blocking buffer with mouse IgG1 (1:100; MG100, Thermo Fisher Scientific). Following overnight incubation at 4°C, the sections were washed in TBS-Tx (4 × 5 min) and incubated at RT for 3 h with biotin-labeled goat anti-mouse IgG1 (abcam, ab97238), diluted 1:300 in TBS-Tx, containing 50% Superblock (Thermo Fisher Scientific; #37580). After washing in TBS-Tx (4 × 5 min), sections were incubated for 1 h with HRP-streptavidin, diluted 1:200 in TBS-Tx containing 50% Superblock. The sections were washed in TBS (4 × 5 min) and developed with 50 mg DAB and 0.01% H_2_O_2_, in TBS containing 10 mM imidazole and 0.5% nickel ammonium hexahydrate (pH 7.4).

For all light microscopy studies, the developed sections were thoroughly washed in dH_2_O, dehydrated in graded alcohols, cleared in xylene, and cover-slipped with PERTEX^®^ (Histolab Products AB).

#### Thioflavin-S staining

Thioflavin-S staining was performed according to Sun et al. (*59*). Fresh-frozen sections were directly immersed in 4% NBF at 4°C for 24 h. The sections were thoroughly washed in ultrapure dH_2_O, dried at RT for 10 min, and defatted for 1 h in a solution of chloroform/99% ethanol (1:1) in the dark. Following hydration through a series of graded ethanols into dH_2_O (2 × 1 min: 99%, 96%, 70%), slides were immersed into an aqueous solution of 0.25% potassium permanganate (5 min), washed in dH_2_O (1 min), and incubated in 1% oxalic acid (2 min). After washing in dH_2_O (2 × 2.5 min), the sections were incubated with freshly-prepared 0.25% NaBH_4_ (2 × 5 min), washed in dH_2_O (5 × 2 min), and transferred into a 0.1% thioflavin-S solution (8 min; dark incubation). The sections were differentiated in 80% ethanol (2 × 10 s), washed in dH_2_O (3 × 5 dips), and incubated for 30 min at 4°C in the dark with high-concentration phosphate buffer, to prevent photobleaching (411 mM NaCl, 8.1 mM KCl, 30 mM Na_2_HPO_4_; 5.2 mM KH_2_PO_4_). Following a dip in dH_2_O, the sections were counterstained with DAPI (30 µM) for 10 min and mounted with Aquatex^®^ mounting medium (Sigma Aldrich Co.).

Immuno(histochemistry) images were acquired with an Olympus DP71 digital camera, mounted on an Olympus BX51 microscope equipped for Epi-Fluorescence illumination, or an Olympus DP80 Dual Monochrome CCD camera, mounted on a motorized BX63 Olympus microscope.

#### [^18^F]Flortaucipir autoradiography

Autoradiography experiments were conducted at the Department of Nuclear Medicine and PET-centre, Aarhus University, Denmark. [^18^F]Flortaucipir was synthesised in Aarhus using previously detailed methods (*66*).

[^18^F]Flortaucipir autoradiography was performed as described previously (*60*), with minor modifications. Sections were thawed to RT for 20 min and fixed/permeabilised in 100% methanol for 20 min. The sections were incubated for a period of 60 min in a 160 mL bath of 10 mM phosphate buffered saline (PBS, pH 7.4), containing 38.4±9.6 MBq [^18^F]Flortaucipir (specific activity: 145±68 GBq/µmol). A series of adjacent brain sections was incubated with identical amounts of radioligand in the presence of 50 µM ‘cold’ flortaucipir, to assess non-specific binding (NSB). Following incubation, sections were serially washed in PBS (1 min), 70% ethanol in PBS (2 × 1min), 30% ethanol in PBS (1 min) and PBS (1 min). After rapid drying under a stream of cold air, the sections were placed in light-tight cassettes and exposed against FUJI multi-sensitive phosphor screens for 30 min (BAS-IP SR2025, GE Healthcare Life Sciences). To allow quantification, standards of known radioactive concentration were prepared by serial dilution of the [^18^F]Flortaucipir incubation solution, and exposed along with the sections. Images were developed in a BAS-5000 phosphor-imager at 25 µm resolution.

For image analysis, the intensity values produced by the ^18^F standards were entered with their corresponding radioactivity values (kBq/mL) into a calibration table, and the relationship between radioactivity and intensity determined by using ImageJ software (v. 1.51c; National Institutes of Health, USA). Adjustments were undertaken to allow for the radioactive decay of [^18^F]Flortaucipir. Values of specific binding were derived after subtraction of NSB from total binding images.

#### Isolation of sarkosyl-insoluble Tau

The general procedure by Greenberg and Davies (*21*) was used to isolate PHFs. Left brain hemispheres from two mice per group were pooled and weighed (467.1±3.1 mg). The tissue was thoroughly homogenised with a motor driven Potter*-*Elvehjem tissue grinder (WHEATON), in a 5-fold excess (v/w) of 10 mM Tris-HCl buffer (pH 7.4), containing 800 mM NaCl, 1 mM EGTA, 10% sucrose, protease inhibitors (cOmplete™Protease Inhibitor; Roche Diagnostics) and phosphatase inhibitors (PhosSTOP™; Roche Diagnostics; H buffer). The homogenate was centrifuged at 4°C for 20 min in a refrigerated ultracentrifuge (27,000 × g; Sorvall RC M150 GX). The supernatant was decanted and kept on ice, the pellet (P1) suspended in 5 vol of H-buffer and re-centrifuged at 27,000 × g for 20 min (4°C). The combined supernatants (S1) were brought to 1% sarkosyl in H buffer and incubated for 2 h at 37°C in a C24 incubator shaker (100 rpm; New Brunswick Scientific). Following centrifugation at 200,000 × g for 40 min (4°C), the sarkosyl-soluble fraction was decanted and kept on ice, and the sarkosyl-insoluble pellet suspended in H buffer, containing 1% CHAPS hydrate (Sigma Aldrich Co.). After filtering through acetate cellulose filters (0.45 µm; VWR International), the filtrates were centrifuged at 200,000 g for 70 min, and the final pellet suspended in 250 µL dH_2_O. Aliquots of 150 µL from the P1, S1, sarkosyl soluble and insoluble fractions were kept for determining Tau protein levels. Samples were stored at −80°C until further processed.

#### Tau Meso Scale

Tau protein concentration in soluble and insoluble fractions was measured with the mouse Total Tau kit (K151DSD-1; Mesoscale Diagnostics LLC). The anti-mouse monoclonal antibody used for detection binds between amino acids 150-200 of Tau, but the clone number and exact epitope recognition site(s) are proprietary. Plates were processed in a SECTOR^®^ Imager 6000 plate reader (Meso Scale Diagnostics LLC), and data acquired with Discovery Workbench software (v.4.0; Meso Scale Diagnostics LLC). Results are presented as pg of Tau/mg of total protein, the latter measured at A562 nm with a Pierce™. BCA protein kit and bovine serum albumin as standard (Thermo Fisher Scientific).

#### Transmission electron microscopy (TEM)

Electron microscopy was performed in the Core Facility for Integrated Microscopy, Faculty of Health and Medical Sciences, University of Copenhagen, Denmark. Carbon-coated copper grids (200 mesh; Ted Pella Inc.) were glow-discharged with a Leica EM ACE 200 (Leica Biosystems Nussloch GmbH), and loaded with 6 µL of sarkosyl-insoluble sample. The sample was adsorbed for 1 min, blotted and stained with 2% phosphotungstic acid in dH_2_0 for 2 min. After blotting and a quick wash with dH_2_0, the samples were examined with a Philips CM 100 TEM (Koninklijke Philips N.V.), operated at an accelerating voltage of 80 kV. Digital images were acquired at a nominal magnification of x180,000, by using an OSIS Veleta digital slow scan 2k × 2k CCD camera and the iTEM software package (Olympus Corporation). Filaments shorter and longer than 200 µm were analyzed in at least two fields of view with ImageJ software. Reported values are mean ± SEM of 70 and 32 PHFs for *APP*_swe_/*PS1*_ΔE9_ and AD tissue, respectively.

#### Mass spectrometry-based proteomics

##### Reduction, alkylation and enzymatic digestion

Sarkosyl-insoluble samples were dried down and denatured at RT with 6 M Urea, 2 M Thiourea and 10 mM dithiothreithol (DTT; Sigma-Aldrich Co.) in dH_2_O (pH 7.5), supplemented with cOmplete™ protease inhibitors and PhosSTOP™ phosphatase inhibitors (Roche Diagnostics). After vortexing and sonication, 100 µg total protein/condition was alkylated in 20 mM iodoacetamide (IAA) for 20 min in the dark. A total of 2 µL of endoproteinase Lys-C was added to the protein sample (6 µg/µL; Wako Chemicals GmbH), and the solution incubated for 2 h at RT. The sample was then diluted 10 times with 20 mM TEAB, pH 7.5, and digested with trypsin overnight (1:50 w/w trypsin:protein). The enzymatic digestion was stopped with 5% formic acid (FA) and the peptide sample cleared by centrifugation (14,000 × g, 15 min). Protein and peptide quantification was performed by Qubit™ fluorometric quantitation (Invitrogen™).

##### Tandem Mass Tag (TMT) labeling

A total of 100 µg tryptic peptides were dried and desalted with R2/R3 columns, before TMT-10plex labeling (AB Sciex Pte. Ltd.). The desalting procedure is detailed below. Labeling was performed according to manufacturer instructions: TMT-126 for WT-3 months(1), TMT-127N WT-3 months(2), TMT-127C Tg-3 months(1), TMT-128N Tg-3 months(2), TMT-128C for WT-24 months(1), TMT-129N for WT-24 months(2), TMT-129C for Tg-24 months(1), 130N for Tg-24 months(2), TMT-130C for human non-AD and TMT-131 for human AD. The labeled peptides from all groups were mixed 1:1, dried down and stored for further enrichment and analysis. Human AD and non-AD samples were included for validation, as well as for taking advantage of the stacking effect of the TMT10-plex, in order to increase identification rates.

##### Enrichment of phosphopeptides and formerly sialylated N-linked glycopeptides (deglycosylated)

Multi- and mono-phosphorylated peptides, as well as sialylated N-linked glycopeptides, were separated from unmodified peptides by using a TiO_2_ workflow (*67*). Modified peptides bind to TiO_2_ beads because the phospho and sialic groups are acidic and retained on the column, whereas unmodified peptides flow-through. The eluted modified peptides were deglycosylated to remove N-linked glycans (*68*). Hydrophilic interaction chromatography (HILIC) was used for sample fractionation prior to nano liquid chromatography-tandem mass spectrometry [LC-MS/MS; (*67*)].

Briefly, the combined labelled peptides (∼1000 µg) were dissolved in TiO_2_ loading buffer [80% acetonitrile (ACN), 5% trifluoroacetic acid (TFA), 1 M glycolic acid] and incubated for 30 min at RT with 6 mg of TiO_2_ beads (5020 Titansphere™ TiO_2_, 5 µm; gift from GL Sciences). The beads were sequentially washed with TiO_2_ loading buffer, 80% ACN/1% TFA and 10% ACN/0.1% TFA. Modified peptides were eluted with 1.5% ammonium hydroxide solution, pH 11.3, and dried-down in a vacuum centrifuge. The unbound TiO_2_ fraction and the combined washing fractions contain unmodified peptides. The dried modified peptides were deglycosylated at 37°C overnight in 20 mM TEAB buffer, pH 8.0, containing N-glycosidase F (P0705L, New England Biolabs Inc.) and Sialidase A™ (GK80046, ProZyme). Unmodified and modified peptides were dried and desalted on micro-columns before capillary HILIC fractionation.

##### Sample desalting with R2/R3 micro-column

Samples were desalted by using home-made P200 tip-based columns, packed with equal ratios of reversed-phase resin material Poros R2 (Oligo R2 Reversed Phase Resin 1-1112-46, Thermo Fisher Scientific) and Poros R3 (OligoR3 Reversed Phase Resin 1-1339-03, Thermo Fisher Scientific). The end of the tip was blocked with C8 material (Model 2314, 3m Empore™ C8). The column was prepared by short centrifugation (1000 × g) of the R3 reversed-phase resin (100% ACN), equilibrated with 0.1% TFA, and centrifuged again. The acidified samples were loaded onto the columns and washed three times with 0.1% TFA. Peptides were eluted with 50% ACN, 0.1% TFA and dried by vacuum centrifugation.

##### HILIC fractionation

The fractions containing unmodified peptides were fractionated prior to nanoLC-MS/MS analysis using HILIC, as described previously (*69, 70*). Peptides were dissolved in 90% ACN, 0.1% TFA (solvent B), and loaded onto a 450 μm OD × 320 μm ID × 17 cm micro-capillary column packed with TSK Amide-80 (3 μm; Tosoh Bioscience LLC). The peptides were separated on a 1200 Series HPLC (Agilent Technologies) over 30 min, by using a gradient from 100–60% solvent B (A = 0.1% TFA), at a flow-rate of 6 μL/min. Fractions were collected every 1 min based on the UV chromatogram. Subsequently, the peptide fractions were dried by vacuum centrifugation.

##### Reversed-phase nanoLC-ESI-MS/MS

The peptides were resuspended in 0.1% FA, and automatically injected on a ReproSil-Pur C18 AQ, in-house packed-trap column (Dr. Maisch GmbH; 2 cm × 100 µm inner diameter; 5 µm). The peptides were separated by reversed phase chromatography at 250 nL/min on an analytical ReproSil-Pur C18 AQ column (Dr. Maisch GmbH), packed in-house (17 cm × 75 µm; 3 µm), which was operated on an EASY-nanoLC system (Thermo Fisher Scientific). Mobile phase was 95% ACN/ 0.1% FA (B) and water/0.1% FA (A). The gradient was from 1% to 30% B over 80 min for mono-phosphorylated, deglycosylated and unmodified peptides, and 1% to 30% solvent B over 110 min for multi-phosphorylated peptides, followed by 30 - 50% B over 10 min, 50 - 100% B over 5 min, and 8 min at 100% B. The nano-LC was connected online to a Q Exactive HF Hybrid Quadrupole-Orbitrap mass spectrometer (Thermo Fisher Scientific), operating in positive ion mode and using data-dependent acquisition. The eluent was directed toward the ion transfer tube of the Orbitrap instrument by dynamic electrospray ionization. The Orbitrap acquired the full MS scan with an automatic gain control target value of 3×10^6^ ions and a maximum fill time of 100 ms. Each MS scan was acquired in the Orbitrap at high-resolution [120,000 full-width half maximum (FWHM)] at m/z 200 with a mass range of 400-1400 Da. The 12 most abundant peptide ions were selected from the MS for higher energy collision-induced dissociation fragmentation (collision energy: 34 V) if they were at least doubly-charged. Fragmentation was performed at high resolution (60,000 FWHM) for a target of 1×10^5^ and a maximum injection time of 60 ms using an isolation window of 1.2 m/z and a dynamic exclusion of 20 s.

##### Data search & analysis

Raw data were searched against the mus musculus or homo sapiens reference databases from swissprot and uniprot via Mascot (v2.3.02, Matrix Science) and Sequest HT search engines, respectively, using Proteome Discoverer (v2.1, Thermo Fisher Scientific). A precursor mass tolerance of 20 ppm and a product ion mass tolerance of 0.05 Da was applied, allowing two missed cleavages for trypsin. Fixed modifications included carbamidomethylation of Cys/Arg and TMT-10plex labeling for Lys and N-termini. Variable modifications contained phosphorylation on Ser/Thr/Tyr, acetylation on the N-termini, oxidation of Met and deamidation of Asn. The TMT datasets were quantified using the centroid peak intensity with the ‘reporter ions quantifier’ node. To ensure high-confident identification, the Mascot percolator algorithm (q value filter set to 0.01), Mascot and Sequest HT peptide rank 1, a cut-off Mascot score value of ≥18, and a Sequest HT?Cn of 0.1 were used. Only high confident peptides were used for further analysis. The peptides were filtered against a Decoy database resulting into a false discovery rate (FDR) of <0.01. Two murine biological replicates per group without missing values were considered for the analysis, and normalization was performed on the protein median. Based on the mean technical variation from repetitive measurements of murine brain samples (*61, 62*), the threshold for determining regulated proteins was set to 1.3-fold, i.e. proteins with TMT ratios >1.30 were considered upregulated and <0.77 downregulated. Tau isoform-specific searches were performed by creating a Uniprot database of isoform sequences for mouse Tau (Uniprot ID: P10637-1, -2, -3. -4, -5, -6) and human Tau (Uniprot ID: P10636-1, -2, -3, -4, -5, -6, -7, -8, -9), as performed by Morris et al. (*71*). Moderate confidence peptides (FDR<0.05) were included in the isoform-specific search.

#### RT-qPCR

For reverse transcription, quantitative polymerase chain reaction (RT-qPCR), Trizol™-isolated RNA (2 μg) from brain sections of WT and TG mice was reverse-transcribed to cDNA, by using the Applied Biosystems™ high-capacity cDNA transcription kit (Thermo Fisher Scientific). Samples were analyzed in triplicate on a StepOnePlus™ Real-Time PCR system (Applied Biosystems™, Thermo Fisher Scientific). Each 20 µL sample contained nuclease-free H_2_O (Thermo Fisher Scientific), 1x Maxima SYBR^®^ green/probe master mix (Thermo Fisher Scientific), 500 nM forward and reverse primers (TAG Copenhagen A/S), 4x diluted cDNA for *Mapt* and 10x diluted cDNA for hypoxanthine phosphoribosyltransferase (*Hprt1*), which was used as a reference gene. *Hprt1* sequences *(72)* and mouse-specific *Mapt* primers spanning exon 10 have been described previously (*73*). Conventional PCR cycling conditions were used [95°C (10 min), followed by 40 cycles of 95°C (15s)/60°C (1 min)], followed by a melt curve. After normalization to *Hprt1*, data were expressed as fold change from the mean value of the 3-month-old WT samples. Nuclease-free H_2_O and genomic DNA were used as controls.

#### Immunoblotting of sarkosyl-insoluble Tau

Ten µg of lysed, denatured protein were separated on 4-12% Bolt Bis-Tris gradient gels (NW04125Box, Novex^®^), and transferred to polyvinylidene fluoride (PVDF) membranes using the Trans-Blot SD Semi-Dry Transfer Cell system (Bio-Rad Laboratories Inc.). Protein content on the PVDF membranes was visualized with PonceauS. Following washing (10 min) and blocking for 1 h in Roti^®-^Block (Carl Roth GmbH), the membranes were incubated overnight at 4°C in blocking solution, containing rabbit anti-tau (1:1000; A0024, Dako Agilent) or rabbit anti-phopshoSer404 primary antibodies (1:200; OAAF07796, Aviva Systems Biology). The blots were washed in TBS+1% Triton X-100 (3 × 15 min; TBS-Tx) and incubated for 2 h with horseradish peroxidase-conjugated secondary antibody (anti-rabbit IgG, HRP-linked antibody, #7074; Cell Signaling Technology^®^). After a final wash in TBS-Tx (3 × 15 min), the blots were developed with enhanced *chemiluminescent* substrate (ECL), according to manufacturer instructions (Luminata™ Forte Western HRP Substrate, WBLUF0100, Merck Millipore).

**Fig. S1.**
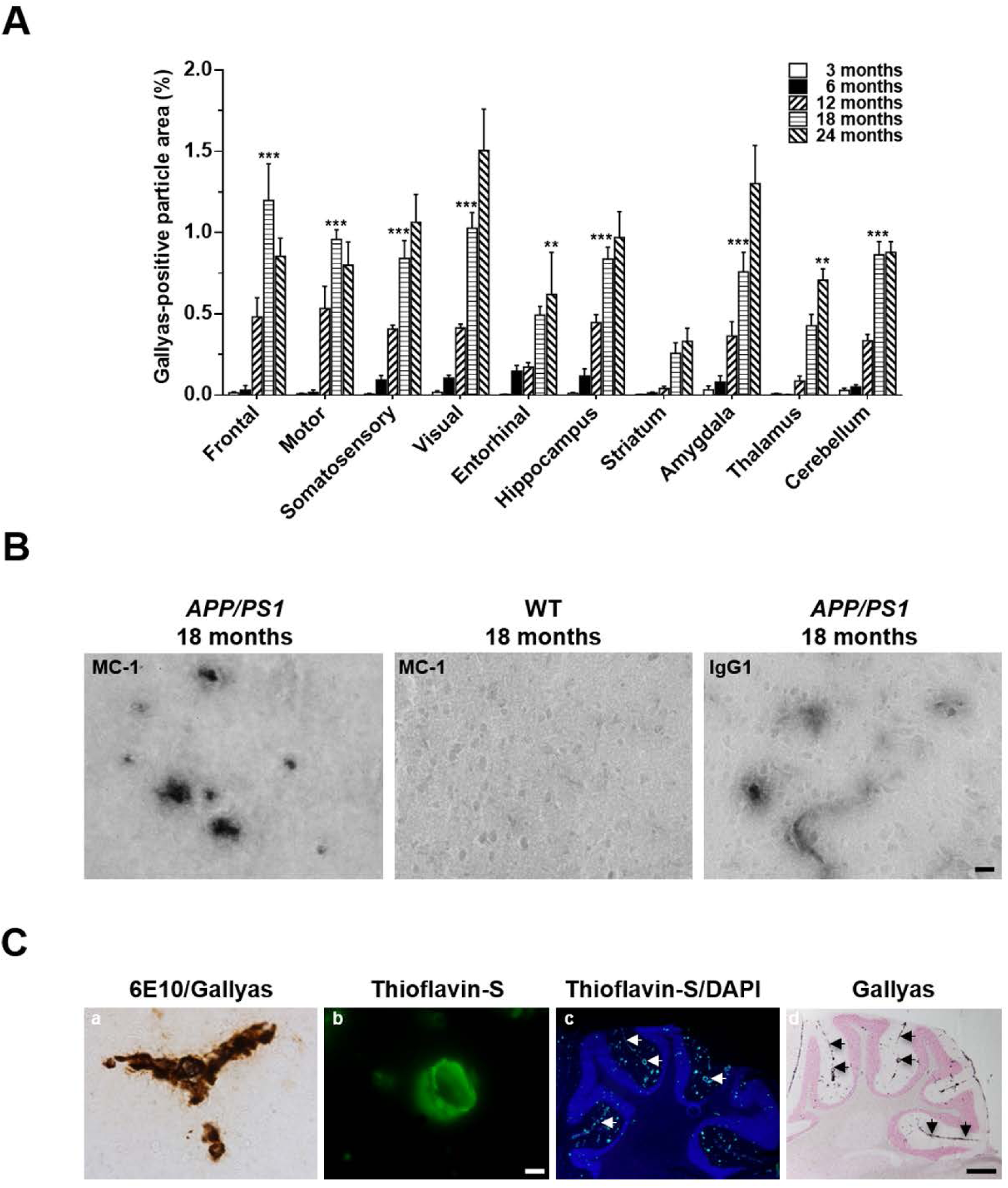
**(A)** Quantification of Gallyas-positive signal in aging *APP*_swe_/*PS1*_ΔE9_ mice. Regions of interest were manually drawn by reference to the mouse brain atlas of Paxinos and Franklin (*74*). Gallyas-positive particles were measured with the particle analysis plugin in ImageJ, after thresholding ROIs on a black and white image display mode, by using default software settings. Data are presented as the mean area fraction occupied by Gallyas-positive particles ± standard error of the mean (SEM), in brain regions of 5-6 animals/group. Asterisks denote the age when Gallyas signal was first increased compared to 3-month old *APP*_swe_/*PS1*_ΔE9_ mice (\*\**P*<0.01, \*\*\**P*<0.001, Bonferroni *post-hoc* tests). Increased silver deposition across all brain areas analyzed was detected in 12- vs. 3- and 6-month-old *APP*_swe_*/PS1*_ΔE9_ mice (*P*<0.001), with additional accumulation occurring in 18- vs. 12- (*P*<0.001), and 24- vs. 18-month-old TG animals (*P*<0.05, Bonferroni *post hoc* tests). Two-way ANOVA confirmed significant main effects of age [F_(4,245)_=169.9, *P*<0.001] and brain region [F_(9,245)_=11.4, *P*<0.001], as well as significant age × region interaction effects on the fraction of brain tissue bearing Gallyas-positive signal [F_(36,245)_=3.2, *P*<0.001]. **(B)** MC-1 immunoreactivity in the neocortex of 24-month-old TG and WT mice. Indications of conformationally modified tau were obtained by using the conformation-dependent MC-1 antibody. In mice, part of the MC-1 signal may be derived from non-specific binding to mouse immunoglobulin 1 (IgG1). Scale bar: 20 µm. **(C)** Vascular and meningeal lesions in 18-month-old *APP*_swe_*/PS1*_ΔE9_ mice. Gallyas/6E10- (a) and thioflavin-S-positive vascular pathology (b). The arrows in (c) & (d) respectively point to thioflavin-S and Gallyas signal in the meninges of the cerebellum. Scale bars: 10 µm (a & b), 200 µm (c & d).

**Fig. S2.**
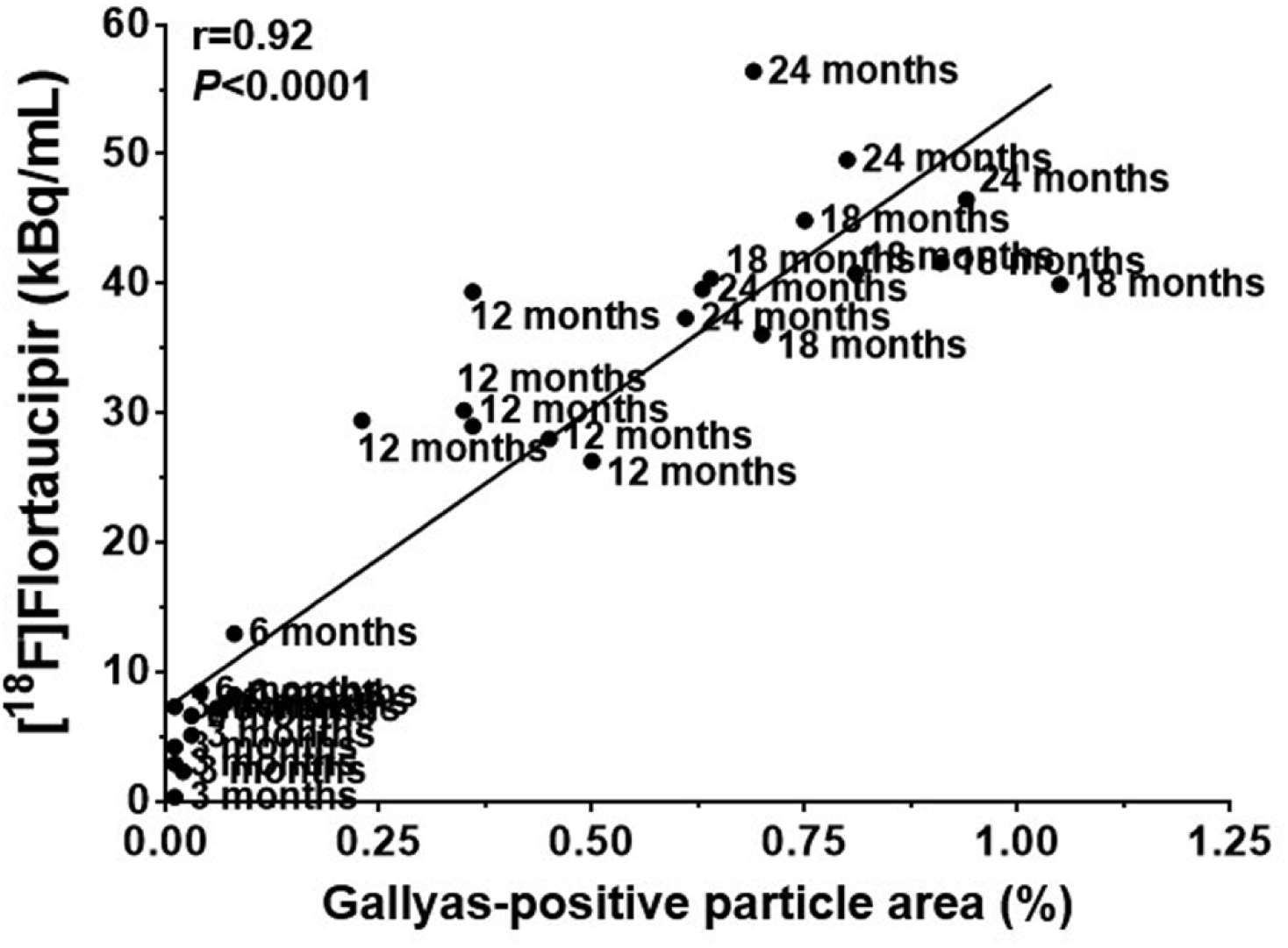
Correlation between Gallyas-positive area fraction and [^18^F]Flortaucipir binding levels. Signals from the Gallyas silver stain and [^18^F]Flortaucipir autoradiography monitor the propagation of identical pathology, most likely tau-associated lesions. Each dot represents values from a single TG animal.

**Fig. S3.**
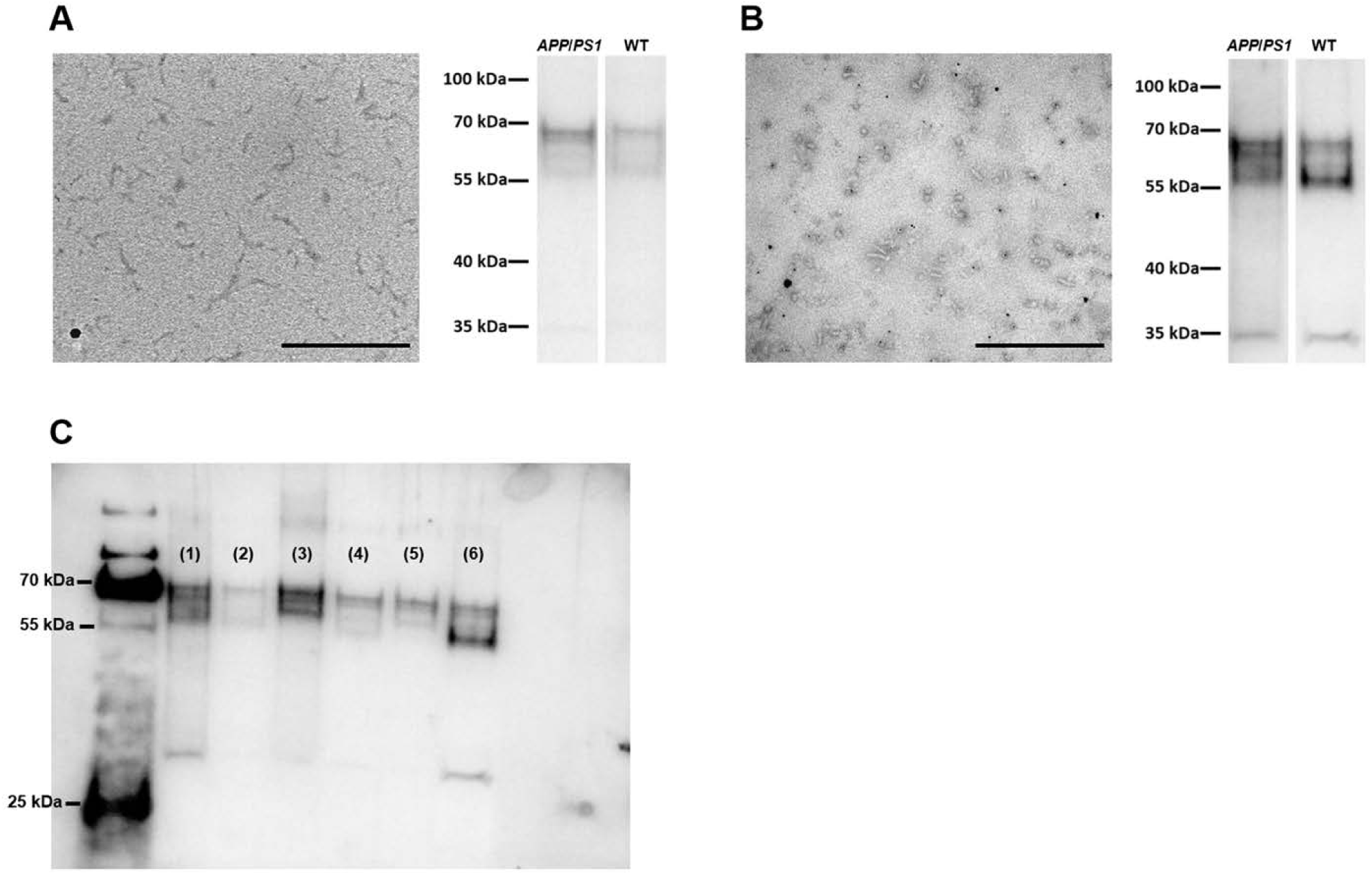
Evaluation of methods for extracting sarkosyl-insoluble tau. TEM and immunoblotting of sarkosyl-insoluble tau, extracted according to **(A)** Sahara et al. (*20*) and **(B)** Greenberg and Davies (*21*). A triplet of immunoreactive bands in the 55-70 kDa range was detected by both methods, by using a rabbit antibody directed to the C-terminal domain of unmodified tau (aa 243-441; A0024, Dako Agilent). **(C)** Bands in A & B were spliced from gel in C, showing total tau immunoreactivity in the following groups: (1) TG 24 months, (6) WT 24 months (Greenberg and Davies method). (2) WT 24 months, (3) Human AD, (4) TG 18 months, (5) TG 24 months (Sarhara et al. method).

**Fig. S4.**
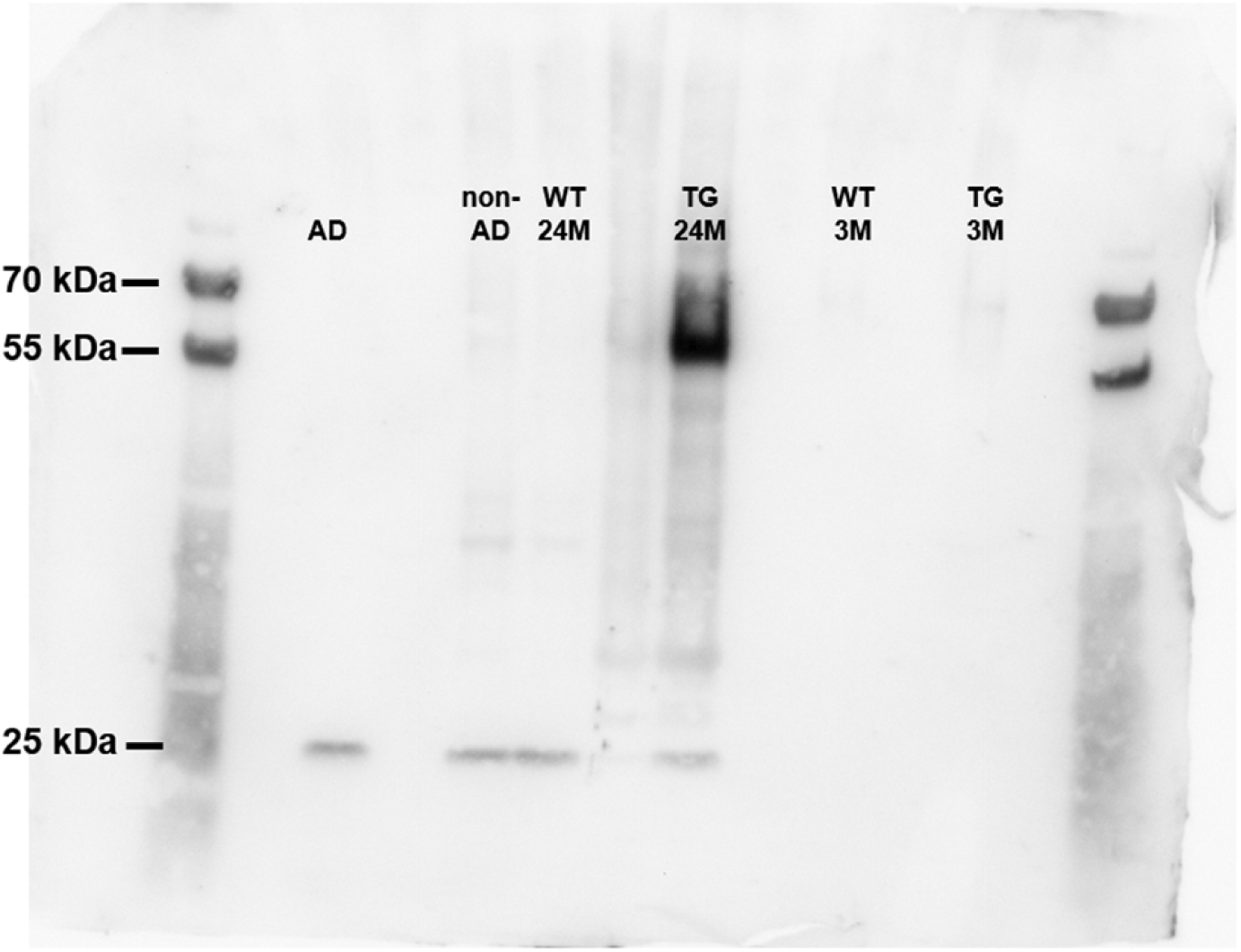
pS404 immunoblot. The accumulation of sarkosyl-insoluble tau hyperphosphorylated at pS404 in 24-month-old transgenic (TG) mice was confirmed with a rabbit anti-phopshoSer404 antibody (1:200; OAAF07796, Aviva Systems Biology).

**Table S1.** Evidence of tau hyperphosphorylation in *APP*_swe_*/PS1*_ΔE9_ mice. Abbreviations: IHC: Immunohistochemistry; WB: Western Blot; IF: Immunofluorescence; NCx: Neocortex; Fr Cx: Frotnal Cortex; Hip: Hippocampus.

| Background strain | Age (Months) | Gender | Method of Euthanasia | Method of Tau Evaluation | Antibody/Epitope | Brain Region | Reference |
| --- | --- | --- | --- | --- | --- | --- | --- |
| B6.C3 | 12 | Female | Anesthesia | IHC | pS262 | NCx | (75) |
| B6.C3 | ~9 | Male | Not reported | IF & WB | pS396 | NCx & Hip | (76) |
| B6.C3 | ~7.5 | Not reported | Anesthesia | IF | AT8 (pS202/pT205) | NCx & Hip | (77) |
| B6.C3 | 6 to >24 | Male & Female | Anesthesia | IHC | PHF1, CP13, pT231, p262, pS396, pS422 | Entire Brain | (78) |
| B6.C3 | 8 | Not reported | Anesthesia | IHC | AT8 (pS202/pT205) | NCx & Hip | (79) |
| B6.C3 | 10 | Male | Not reported | WB & Gallyas | AT8 (pS202/pT205) | Entire Brain | (80) |
| B6.C3 | 11 & 18 | Not reported | Anesthesia | IHC | AB1518 (not reported) | Hip | (81) |
| B6.C3 | 11 | Not reported | Anesthesia | IHC | PHF1 (pS396/pS404) | NCx & Hip | (82) |
| B6.C3 | ~7.5 | Not reported | Anesthesia (?) | WB & IHC | AT8 (pS202/pT205)<br>PHF1 (pS369/pS404) | NCx & Hip | (83) |
| B6.C3 | ~7 | Male | Anesthesia | WB | pS519, pS202, pS235, pS396, pS404 | FrCx & Hip | (84) |
| C57BL/6 | 6 | Female | Cervical dislocation | WB | pT205, pS396, pS404 | Hip | (85) |
| C57BL/6J | 22 | Female | Anesthesia | WB & IHC | pS199, pS396 | NCx & Entire Brain | (86) |
| C57BL/6J | 7 | Male | Not reported | IF | pS199 | NCx & Hip | (87) |
| C57BL/6J | 3-12 | Not reported | Not reported | WB | PHF1 (pS396/pS404) | FrCx | (88) |
| C57BL/6J | 12 | Male | Cervical dislocation | Proteomics & Gallyas | N/A | NCx & Hip, Olf. Bulb, Brainstem | (61) |
| C57BL/6 | ~9 | Female | Anesthesia | WB | pS235, pT205 | Entire Brain | (89) |
| C57BL/6J | ~7.5 | Male | Not reported | WB | pS262 | NCx & Hip | (90) |
| C57BL/6 | 7 | Male | Decapitation | WB | PHF1 (pS396/pS404) | Hip | (91) |
| C57BL/6 | ~12 | Female | Anesthesia | WB | pS235, pT205 | NCx & Hip | (92) |
| C57BL/6 | 3 & 6 | Male | Not reported | WB | pS199, pT205, pS396, pS404 | Hip | (93) |
| C57BL/6J | ~7.5 | Male | Not reported | IF | pT181 | NCx & Hip | (94) |
| C57BL/6J | 6 & 9 | Male | Anesthesia | IF | pS199, pS202, pS262, pT181 | NCx | (95) |
| C57BL/6 | 6-7 | Not reported | Anesthesia | IHC | AT8 (pS202/pT205) | Amygdala | (96) |
| C57BL/6J | 12 | Not reported | Anesthesia | IHC/IF | AT8 (pS202/pT205)<br>PHF1 (pS369/pS404) | NCx | (5) |
| Not reported | ~12 | Male | Anesthesia | WB | pS235, pT205 | Entire Brain | (97) |
| Not reported | 3-12 | Not reported | Anesthesia | WB | pS199, pT205, pS396, pS404 | Entire Brain | (98) |
| Not reported | 6 | Not reported | Anesthesia | WB | AT8 (pS202/pT205) | NCx | (99) |
| Not reported | 7 | Not reported | Anesthesia | WB | pS396 | Brain hemisphere | (100) |
| Not reported | ~7 | Not reported | Anesthesia | WB | AT8 (pS202/pT205) | NCx | (101) |
| Not reported | >18 | Not reported | Anesthesia | IHC | pS199, pT231 | Cerebellum | (102) |
| Not reported | ~2-3 | Male | Anesthesia | WB | PHF1 (pS396/pS404) | NCx & Hip | (103) |
| Not reported | 2 | Male | Anesthesia | WB | AT8 (pS202/pT205)<br>PHF1 (pS396/pS404) | Subventricular zone, NCx & Hip | (104) |

## DATA FILE S1

**REGULATED PROTEINS 3 MONTHS: TG vs. WT**

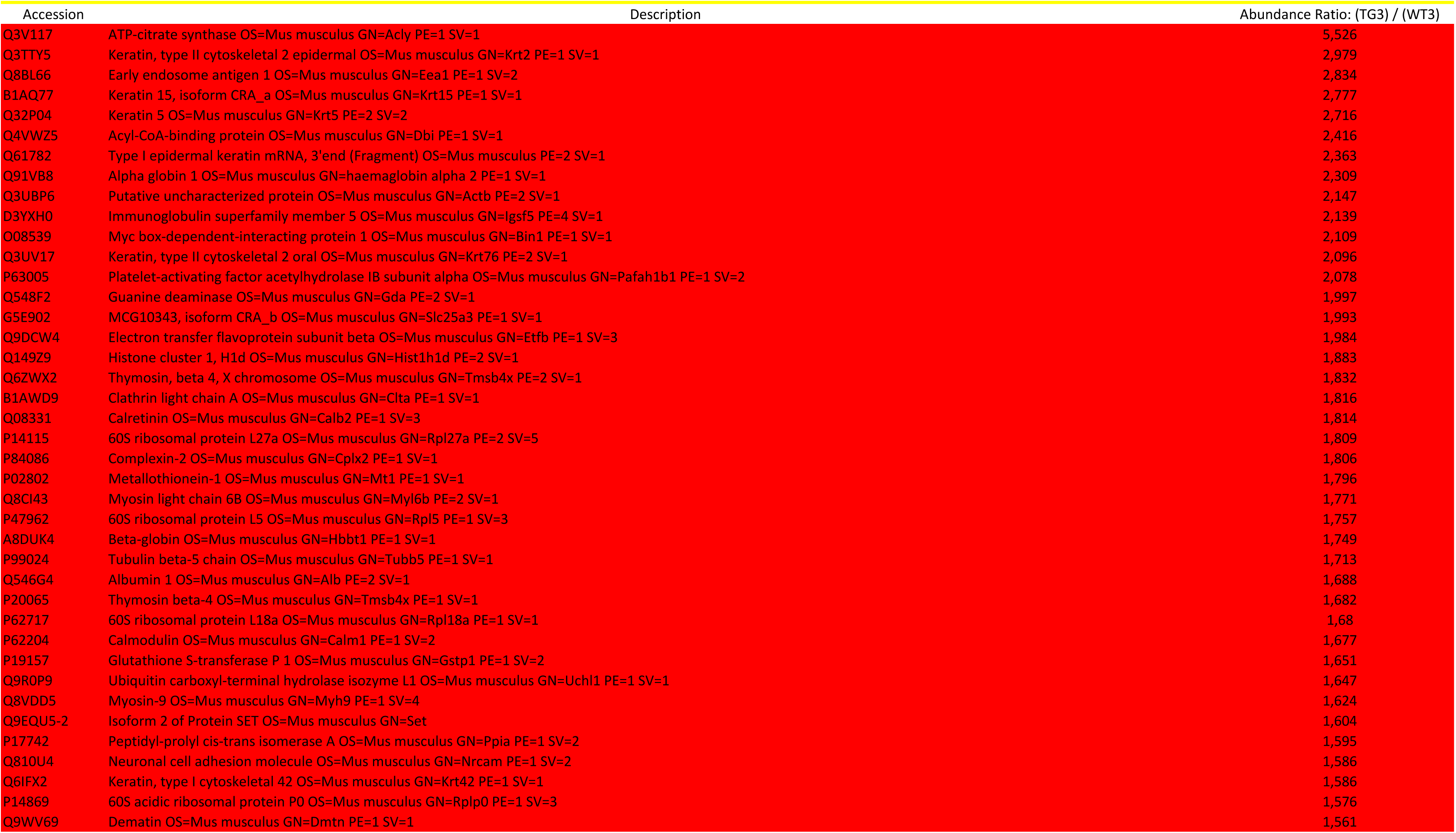

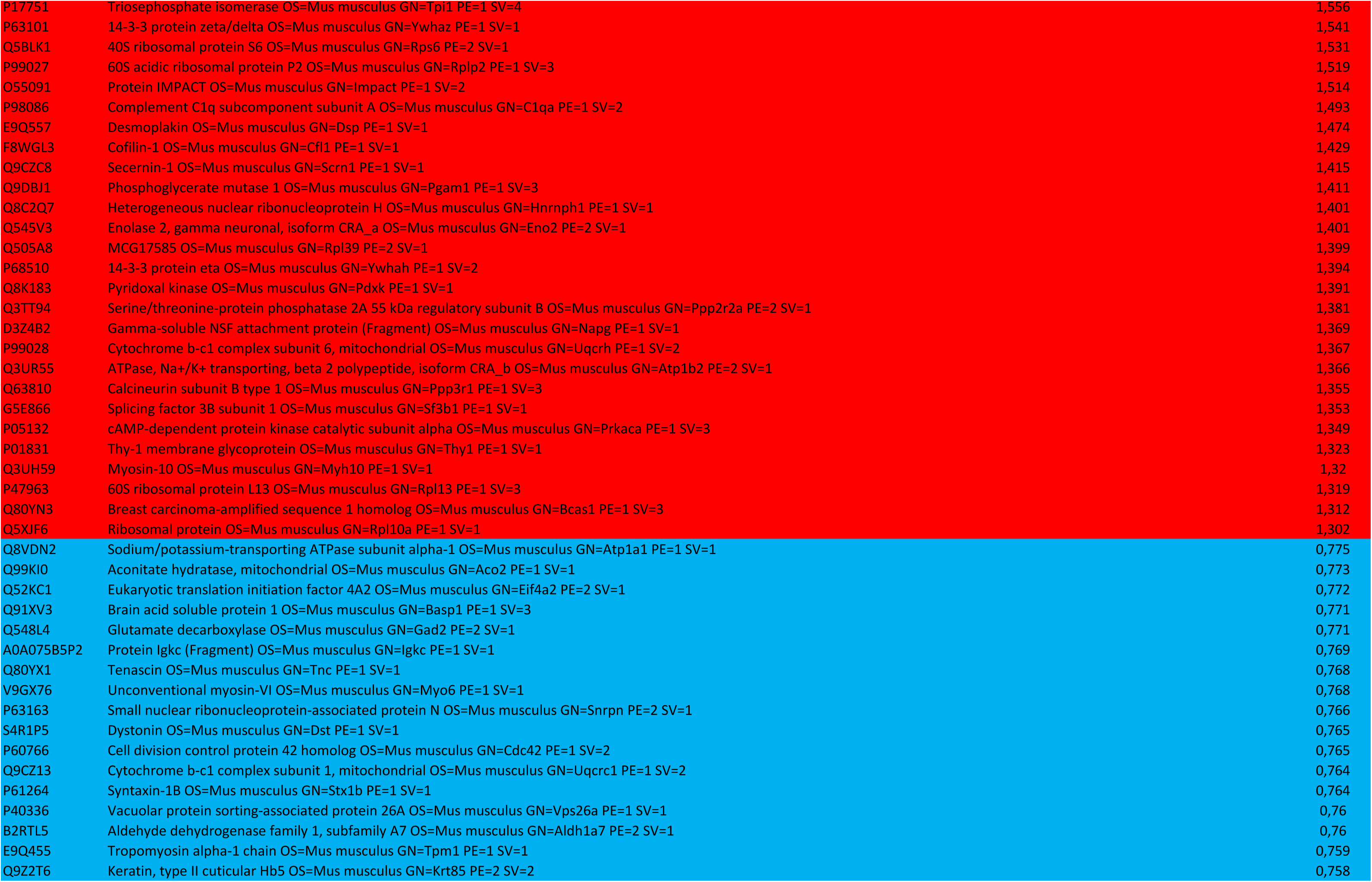

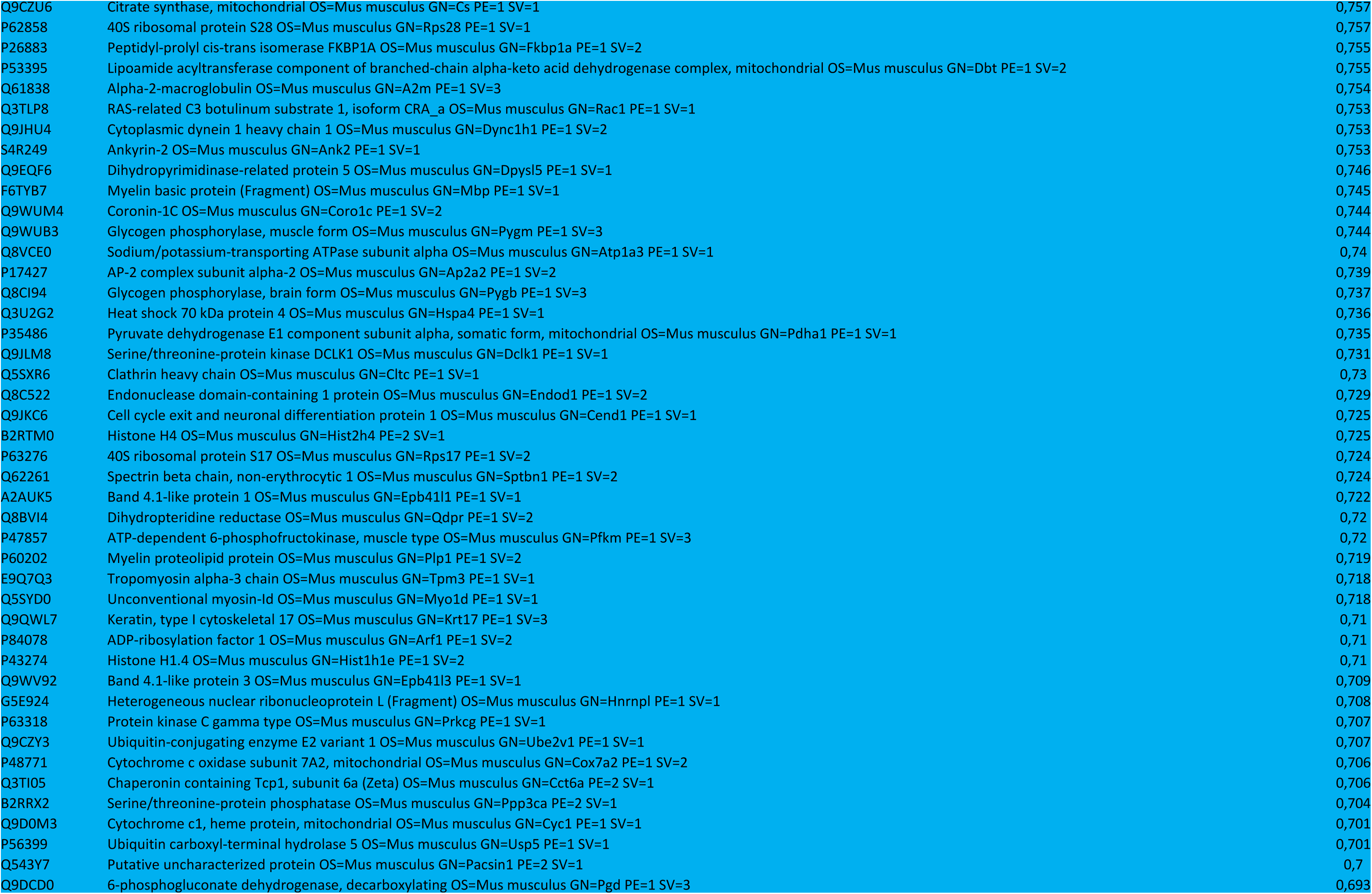

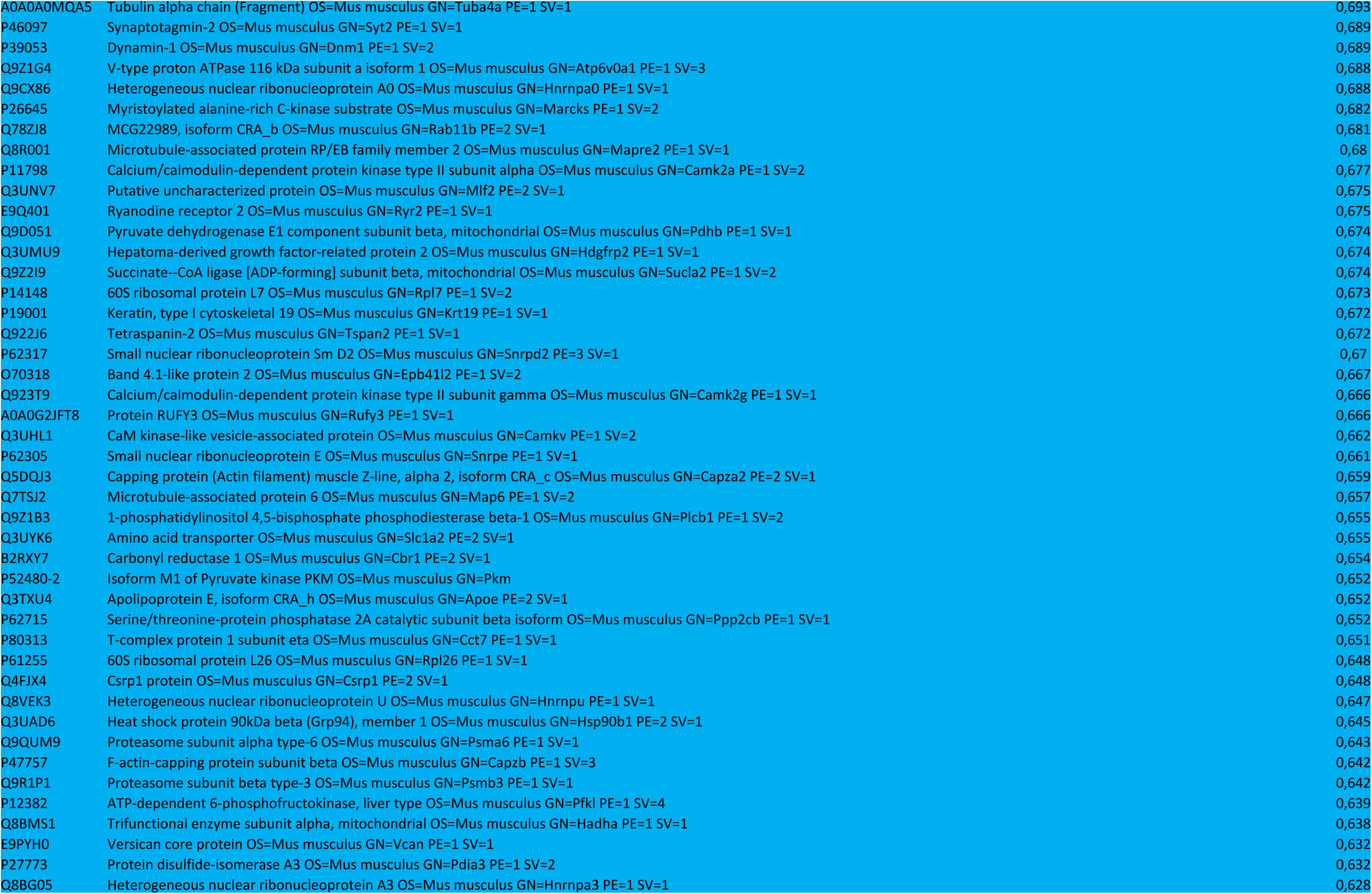

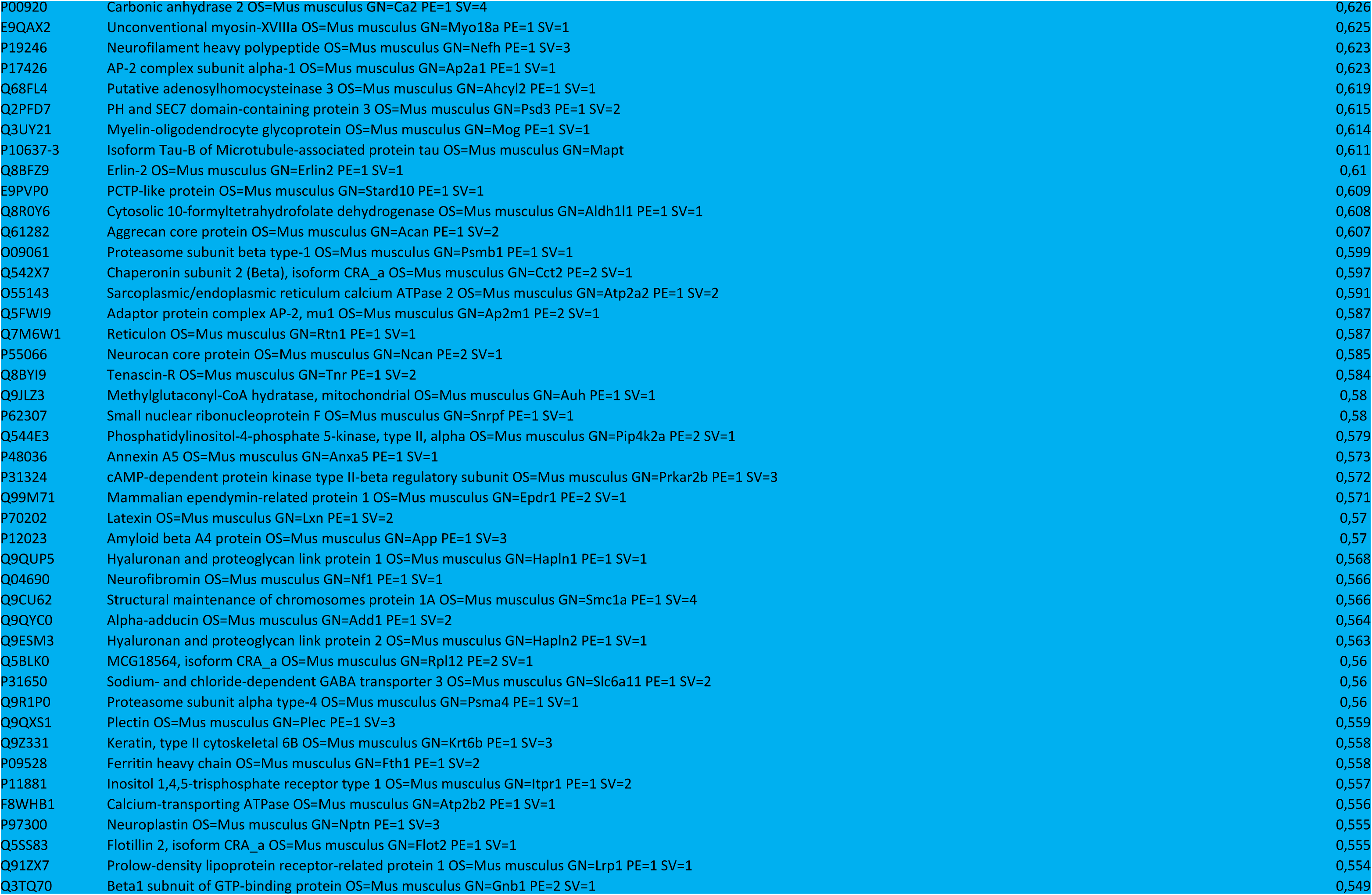

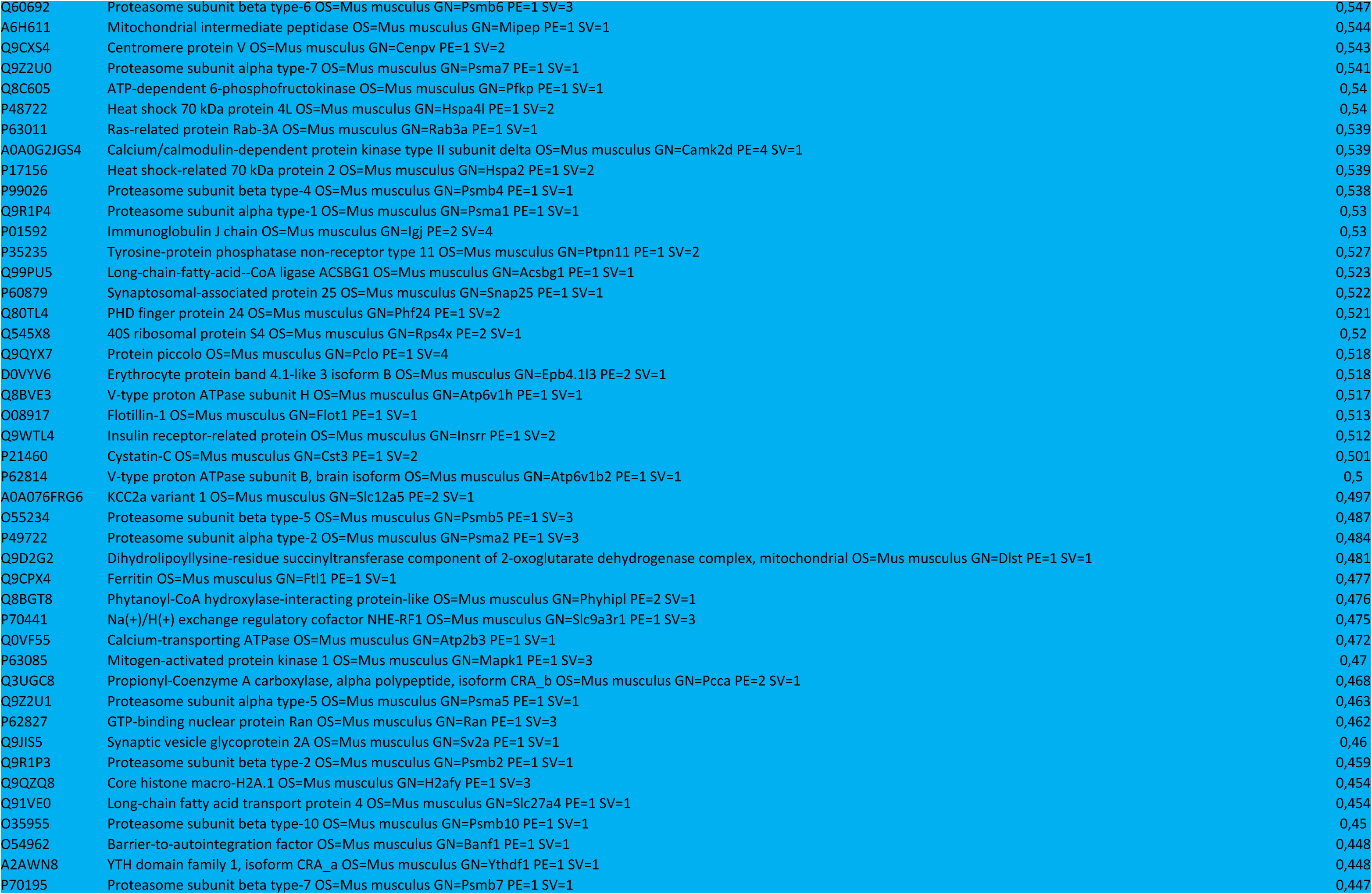

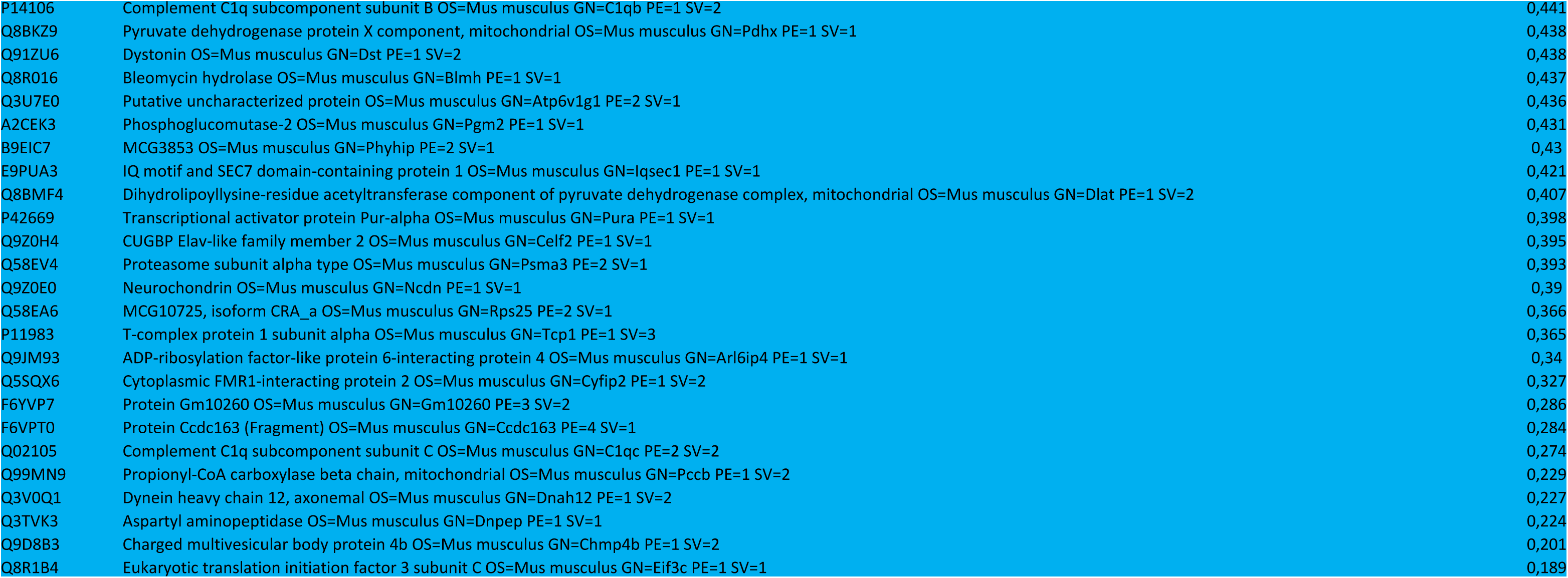

## DATA FILE S1

**REGULATED PROTEINS 24 MONTHS: TG vs. WT**

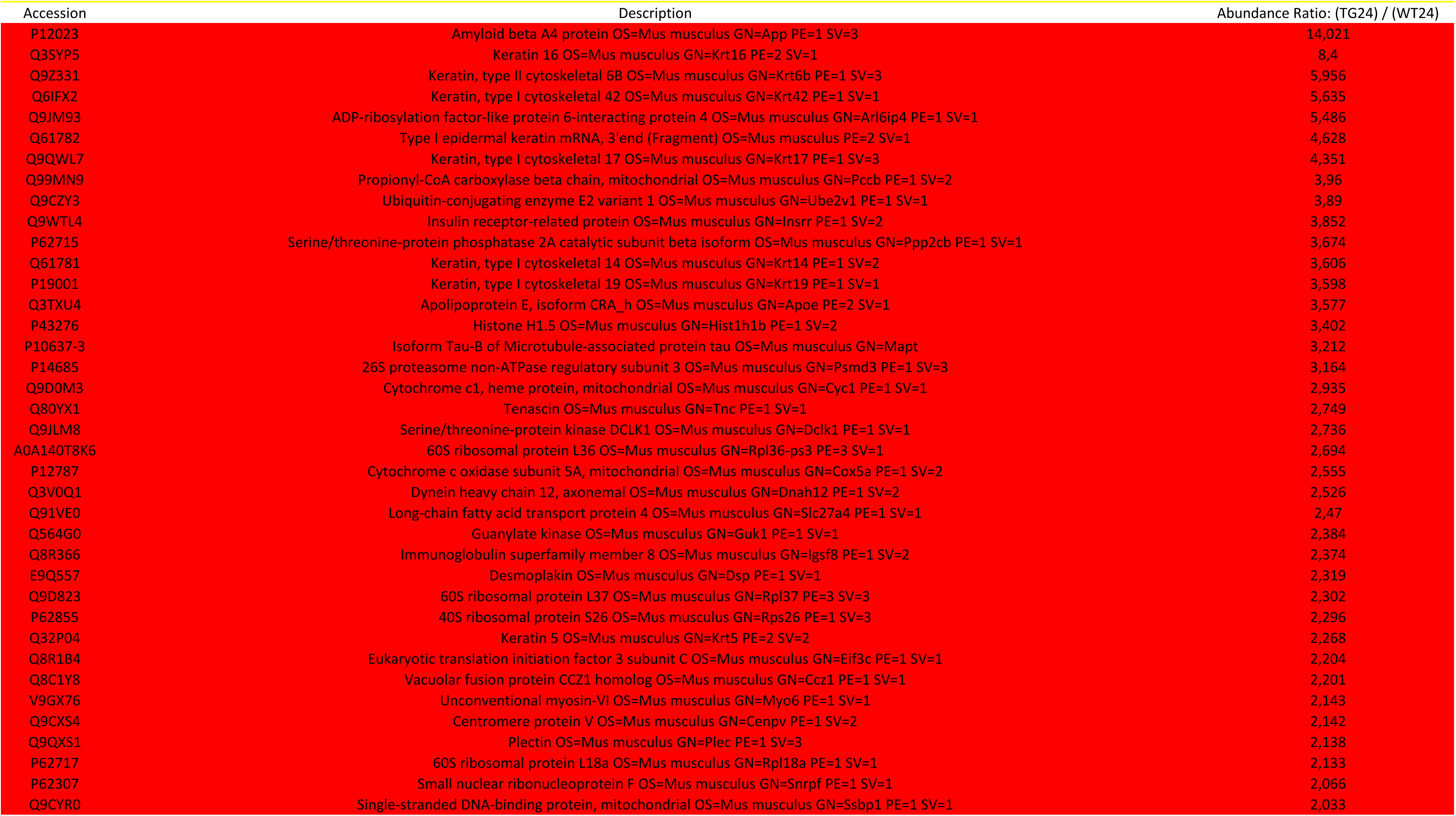

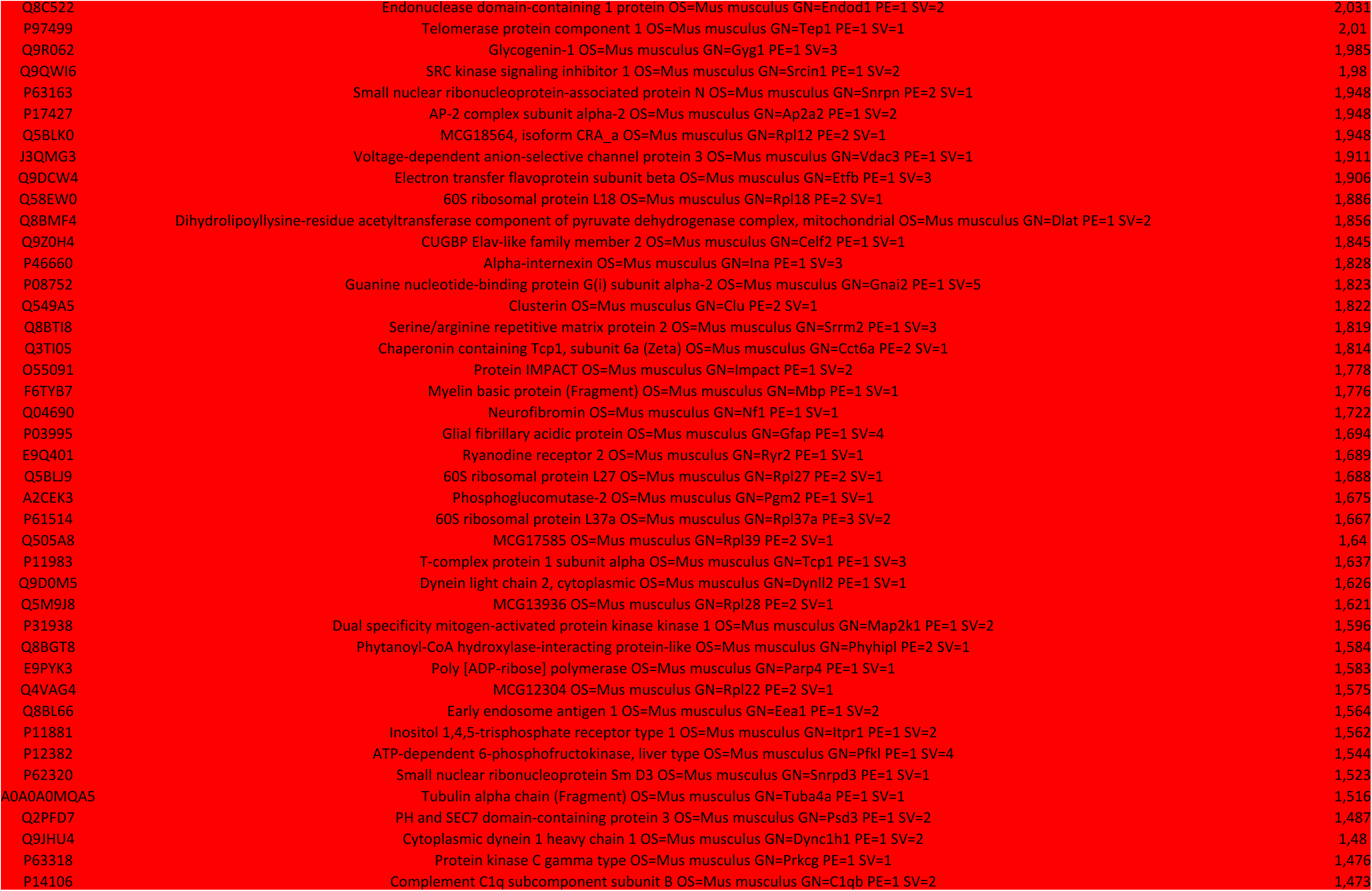

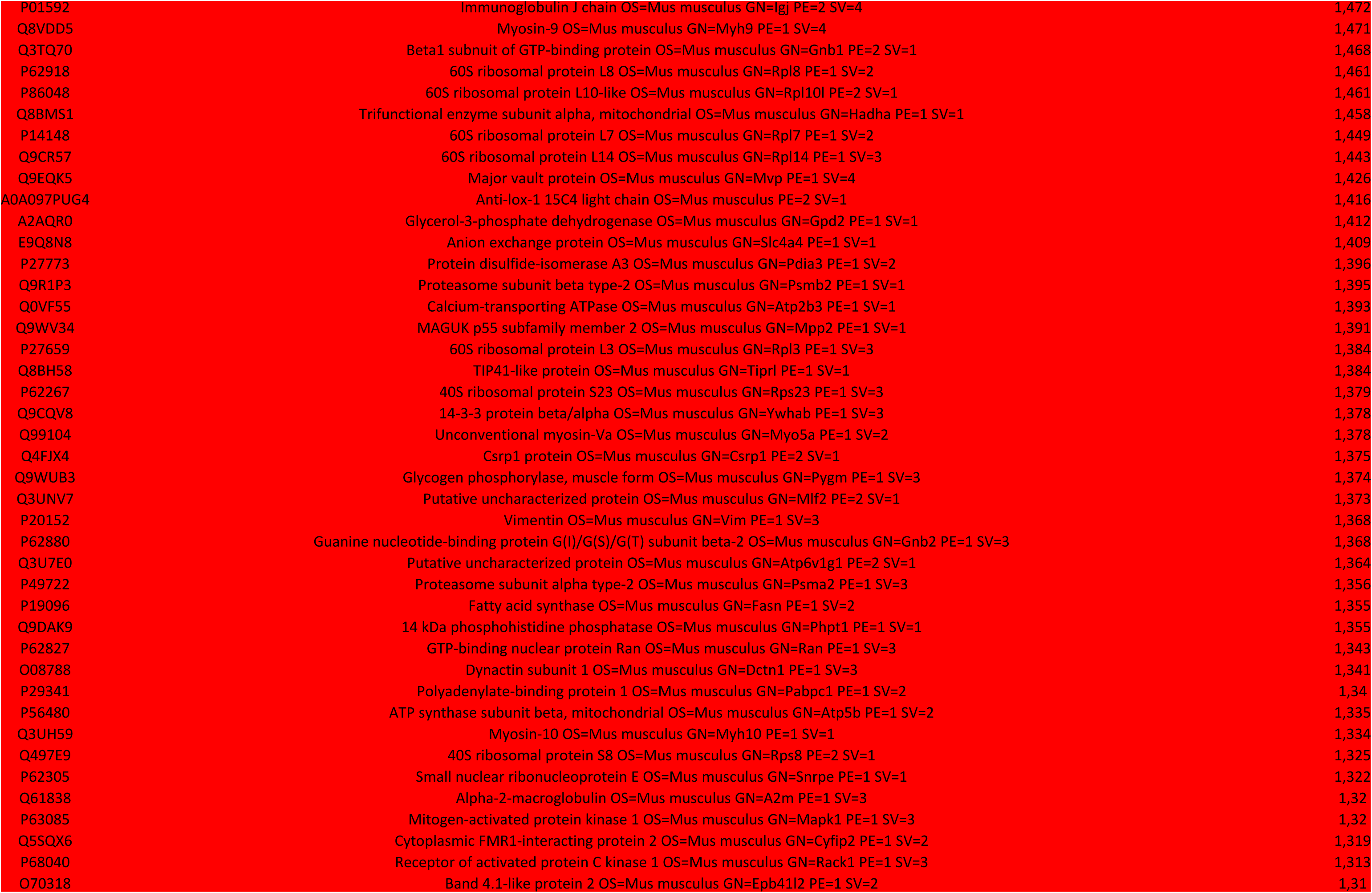

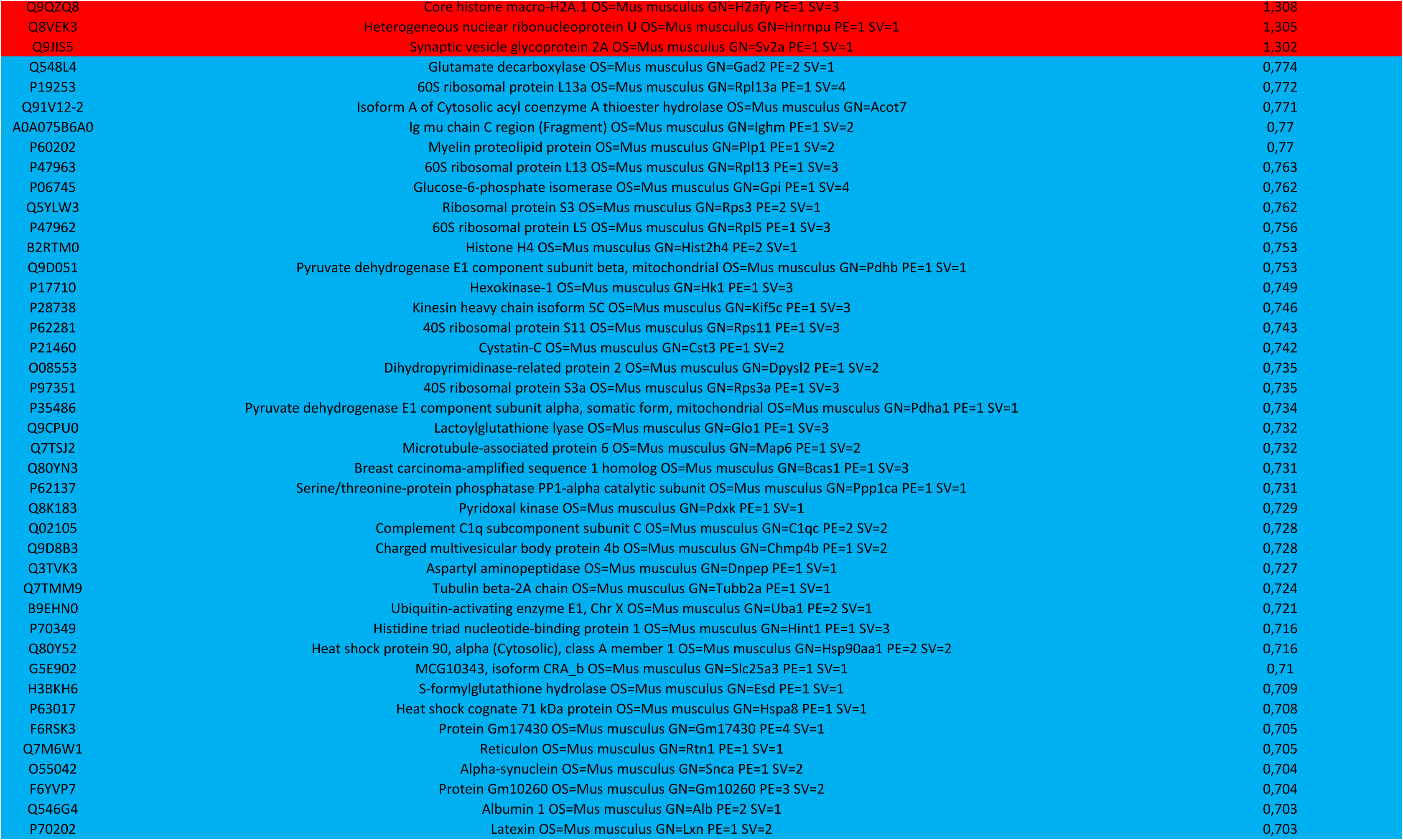

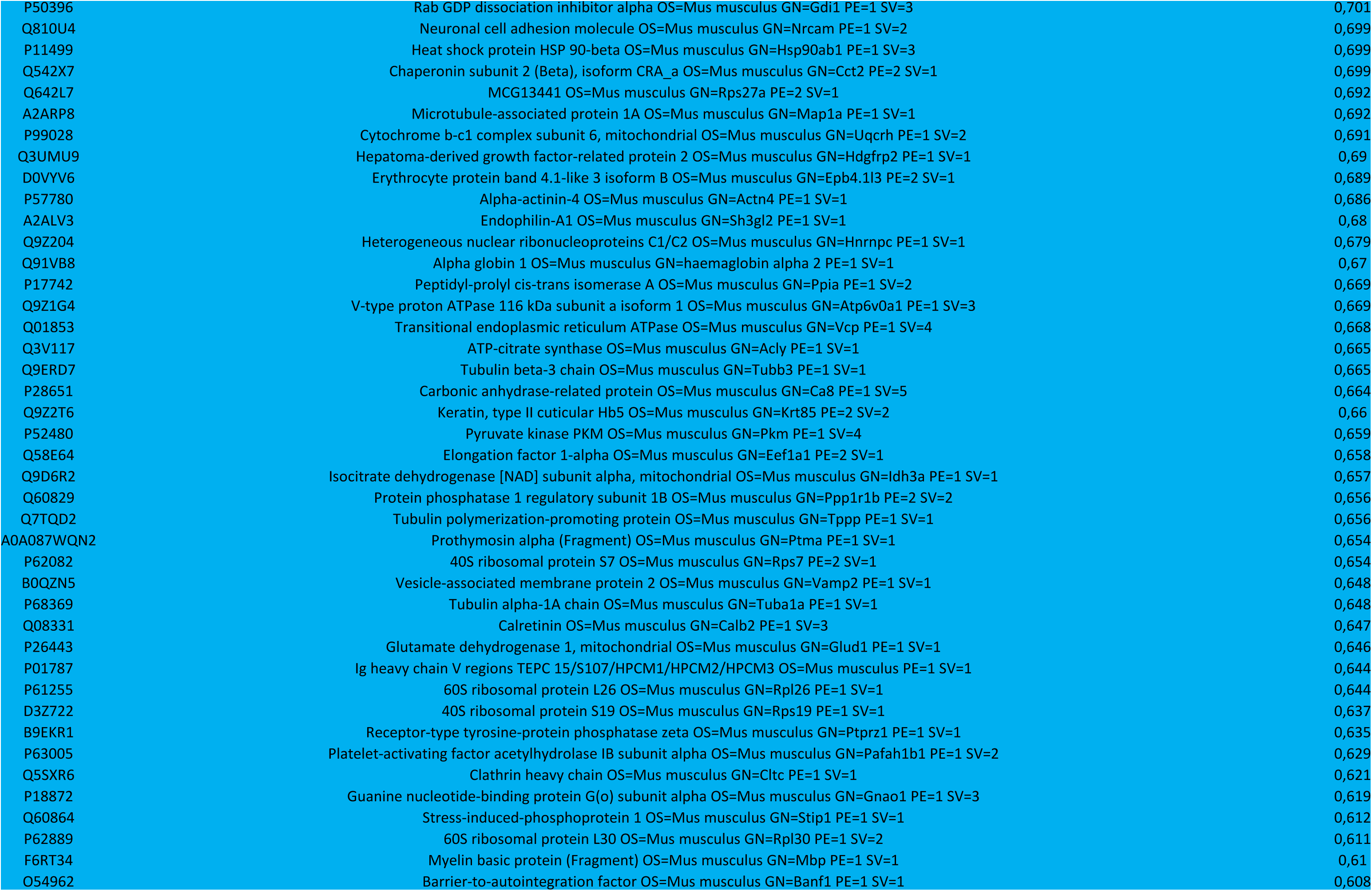

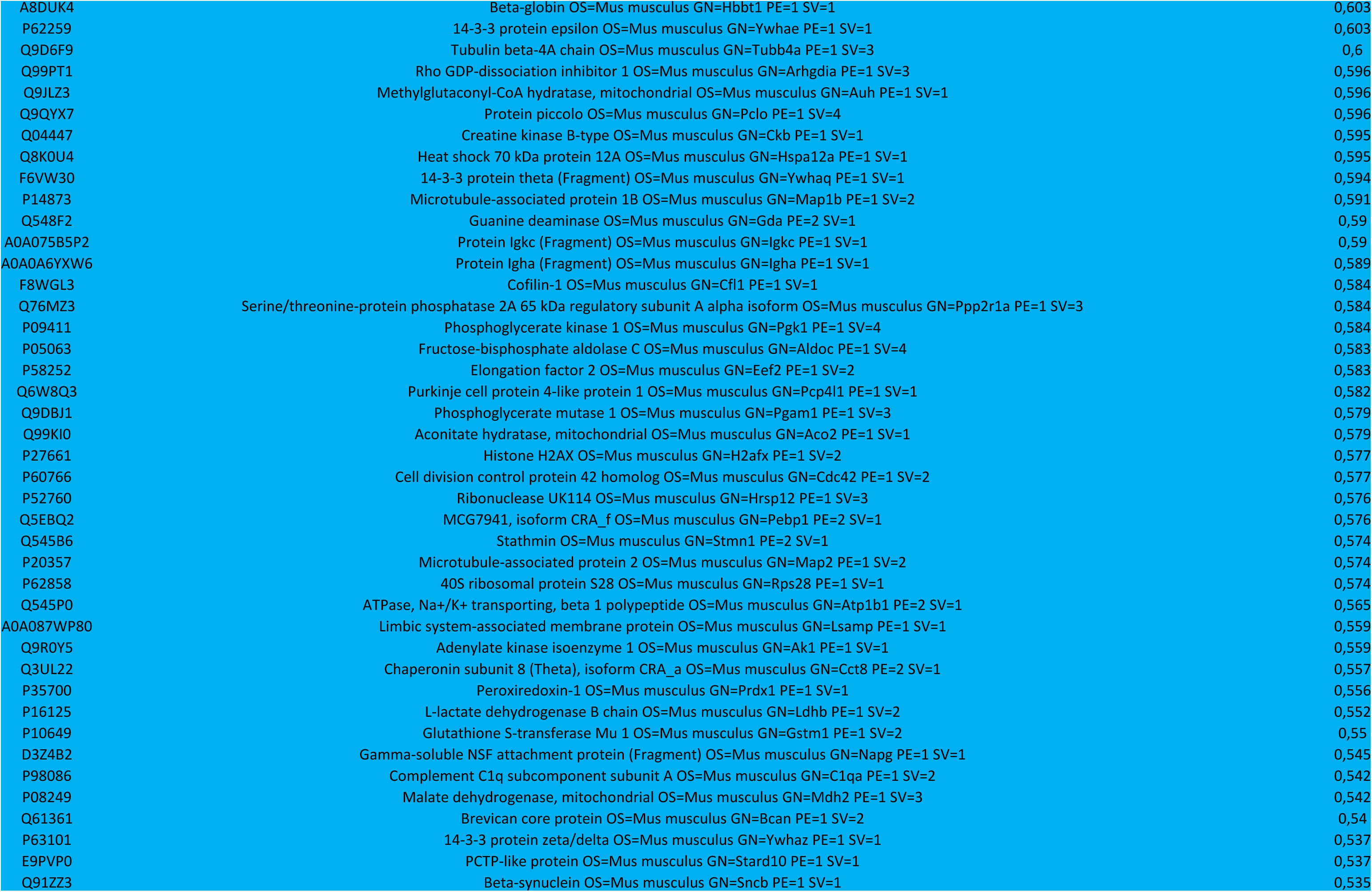

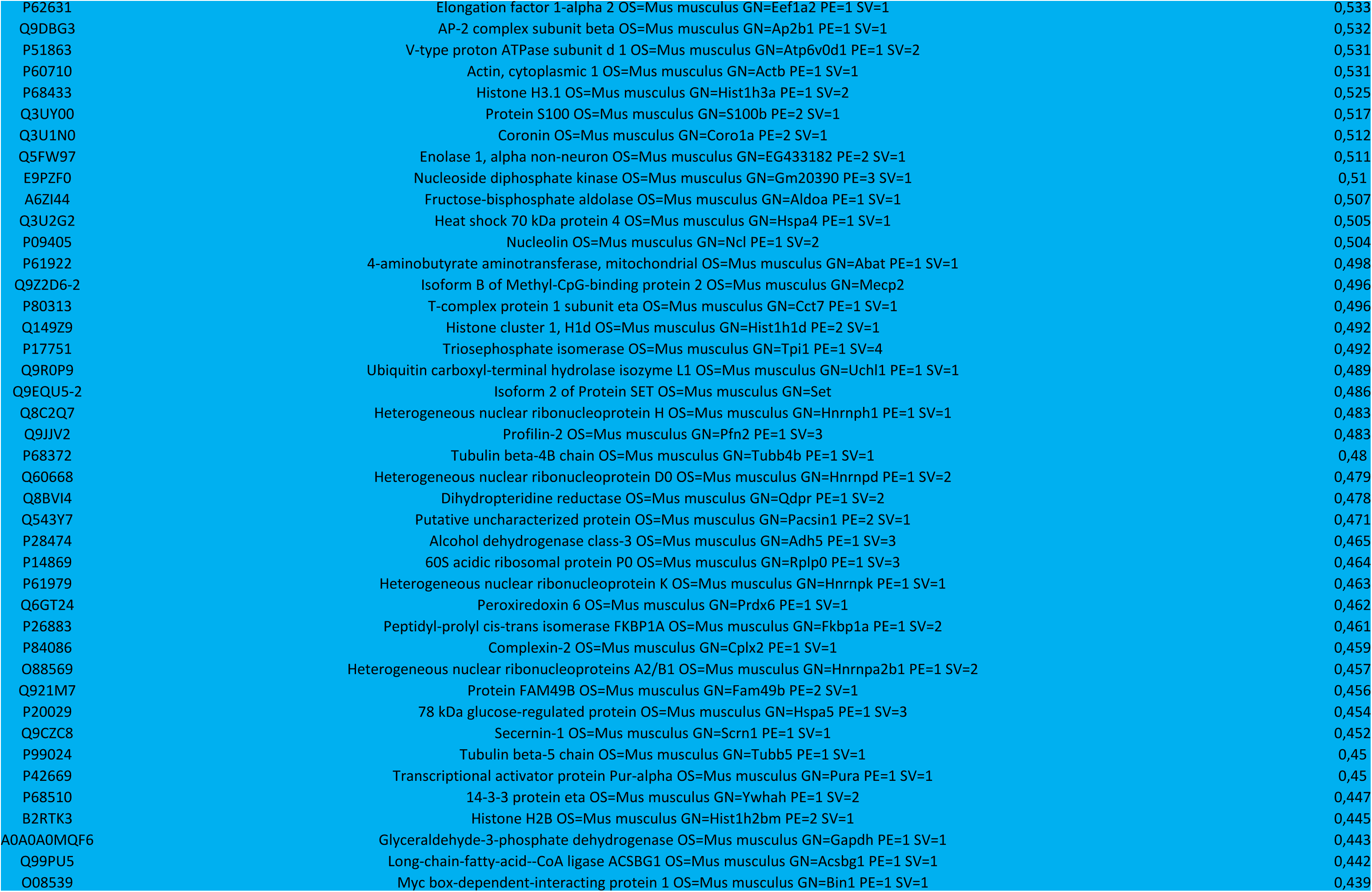

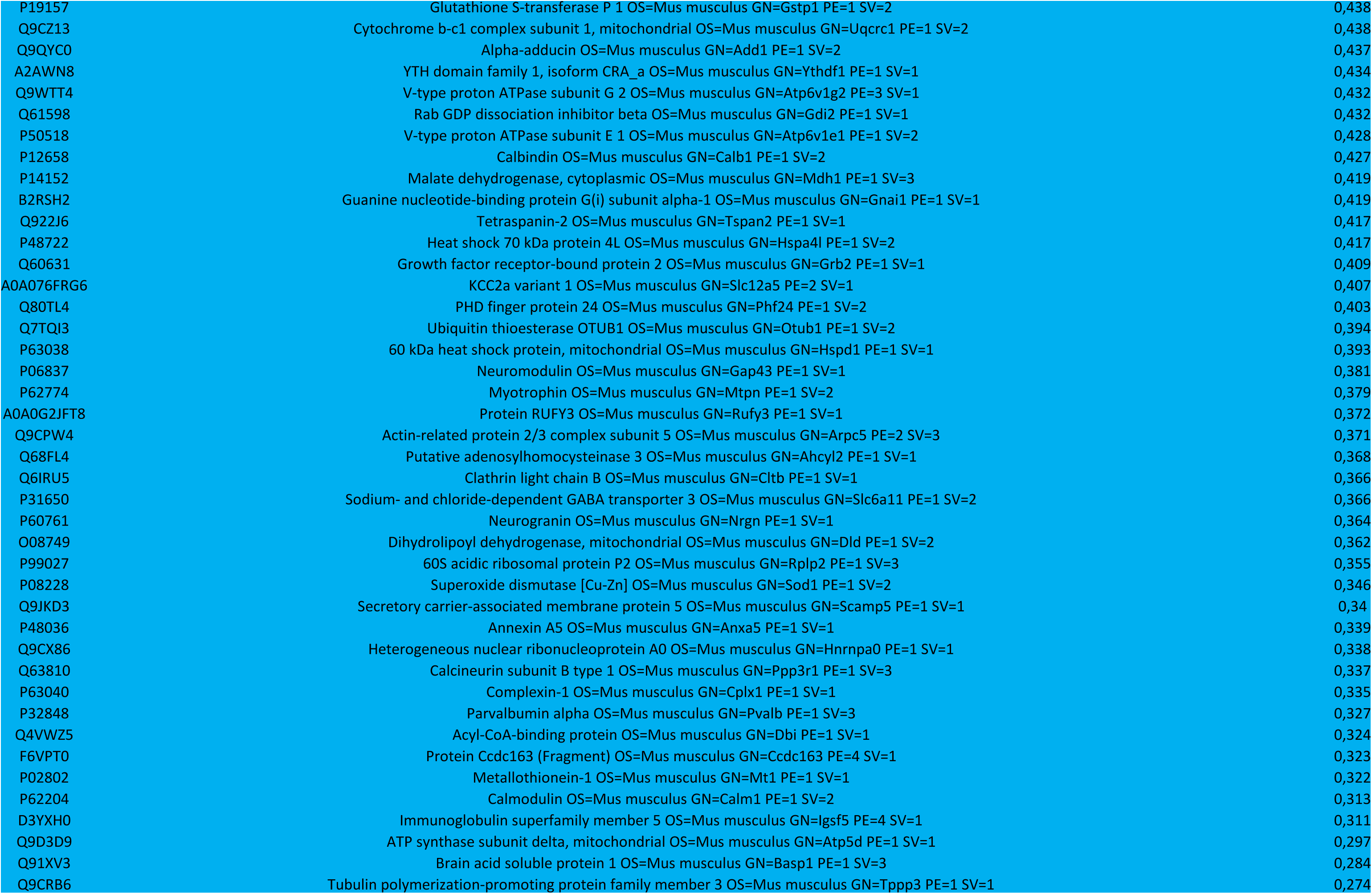

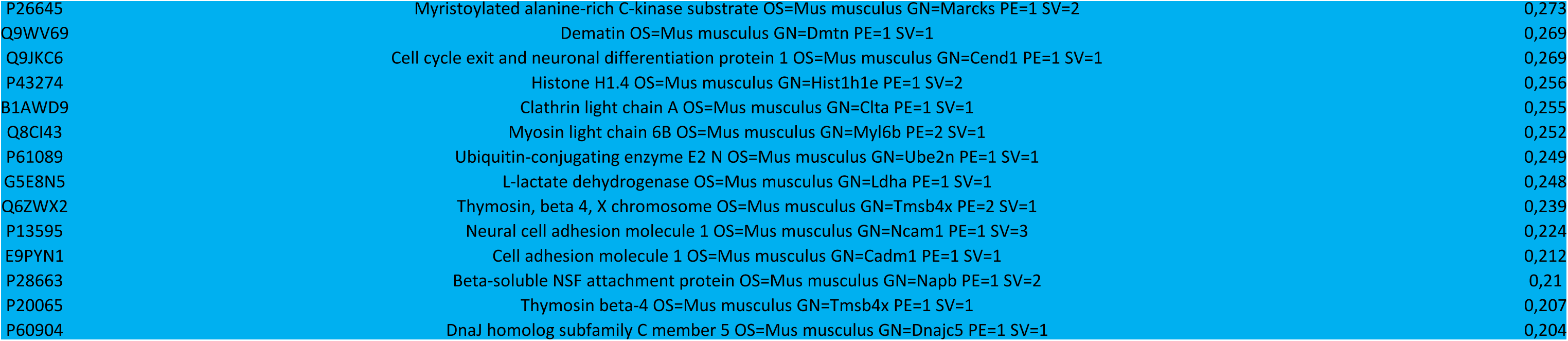

## DATA FILE S1

**REGULATED PROTEINS TG: 24 vs. 3 months**

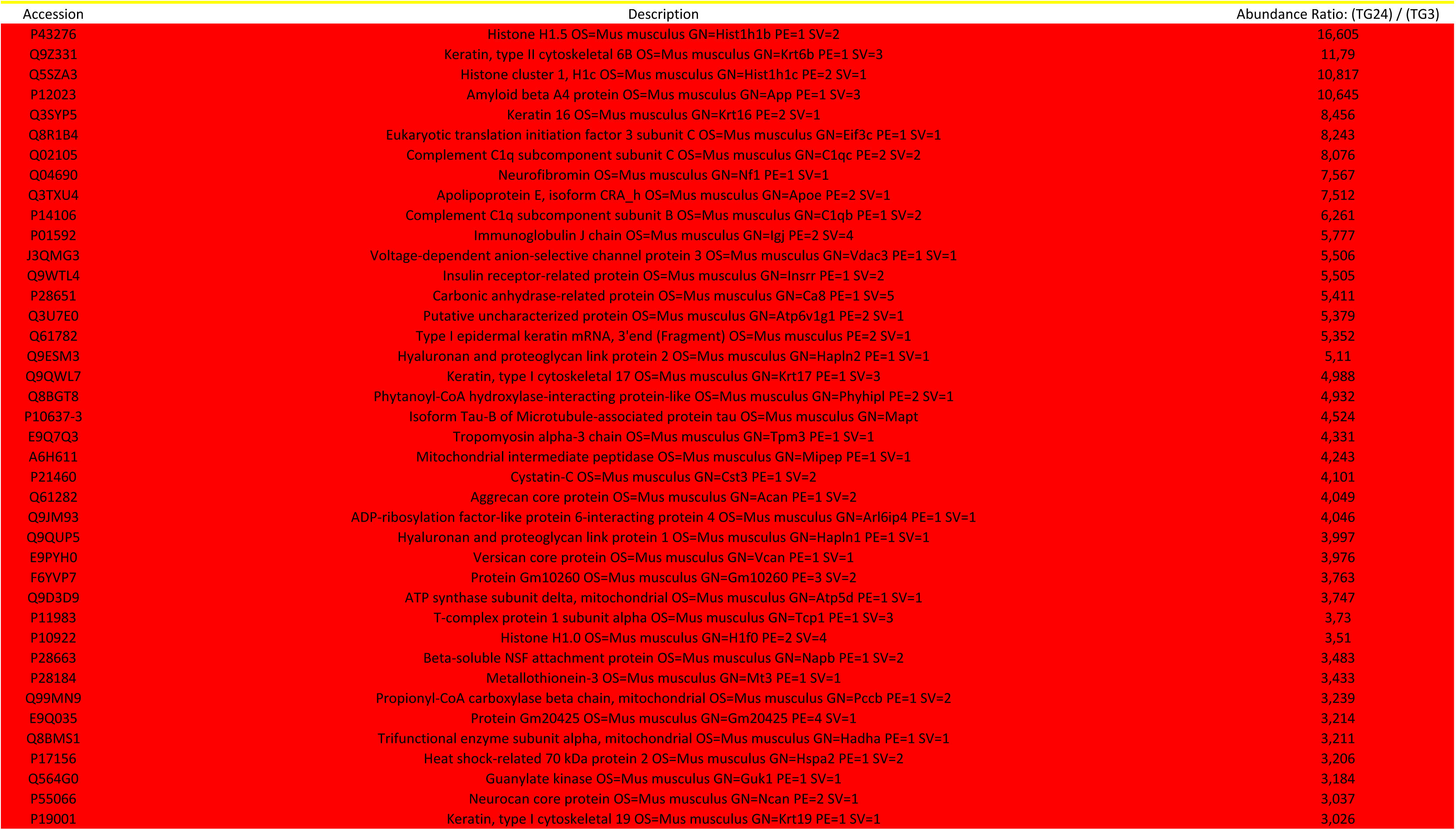

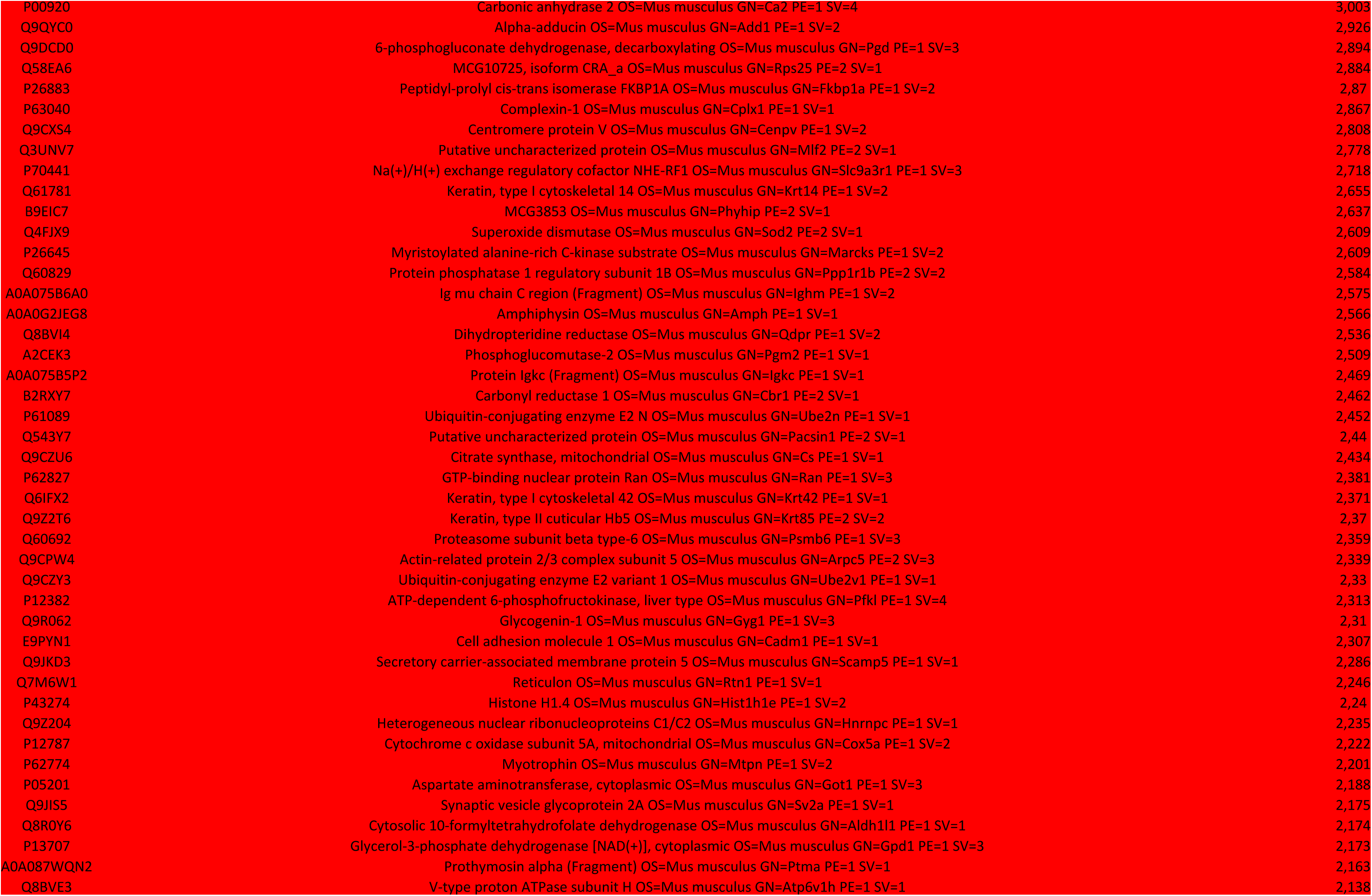

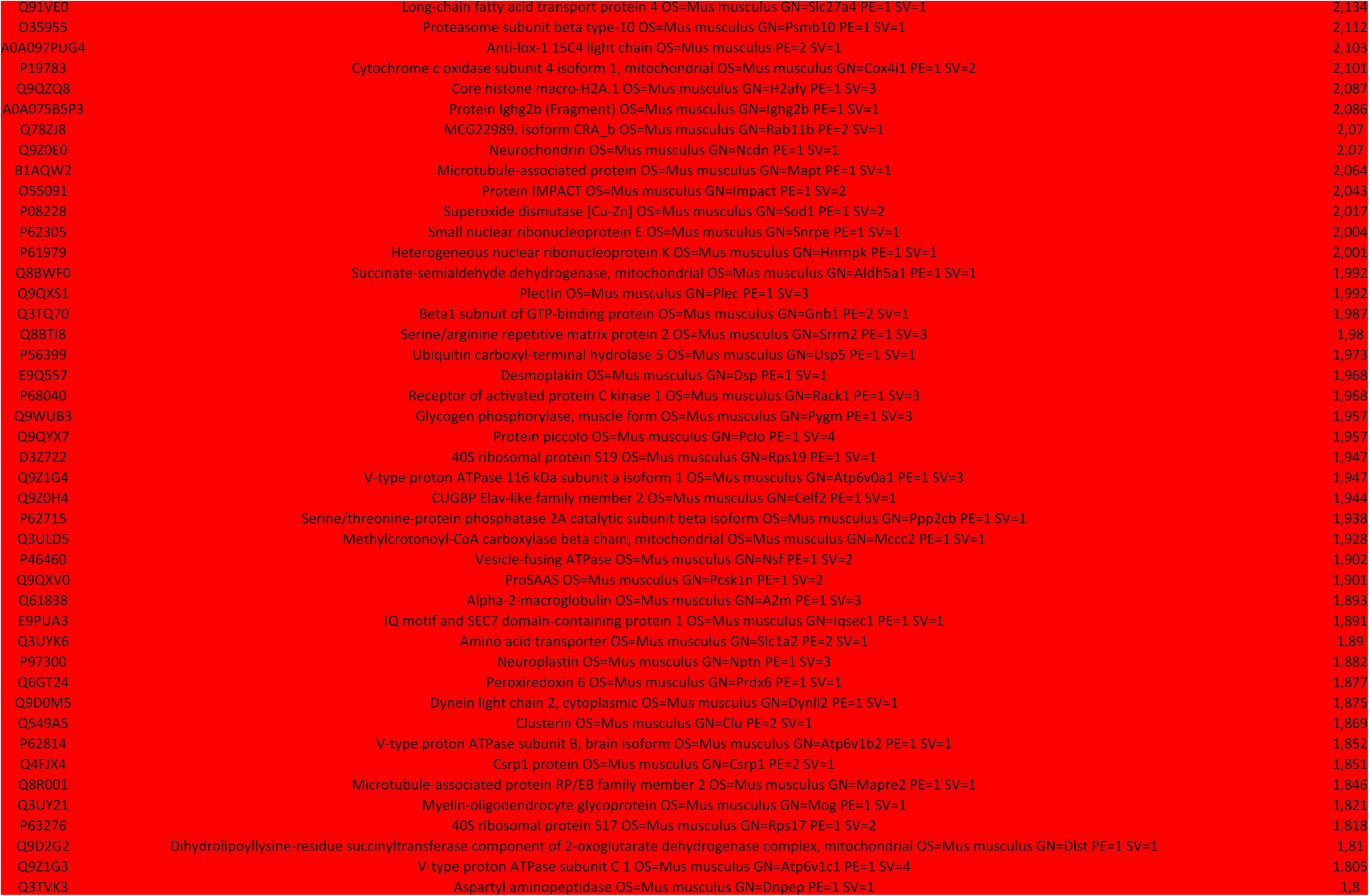

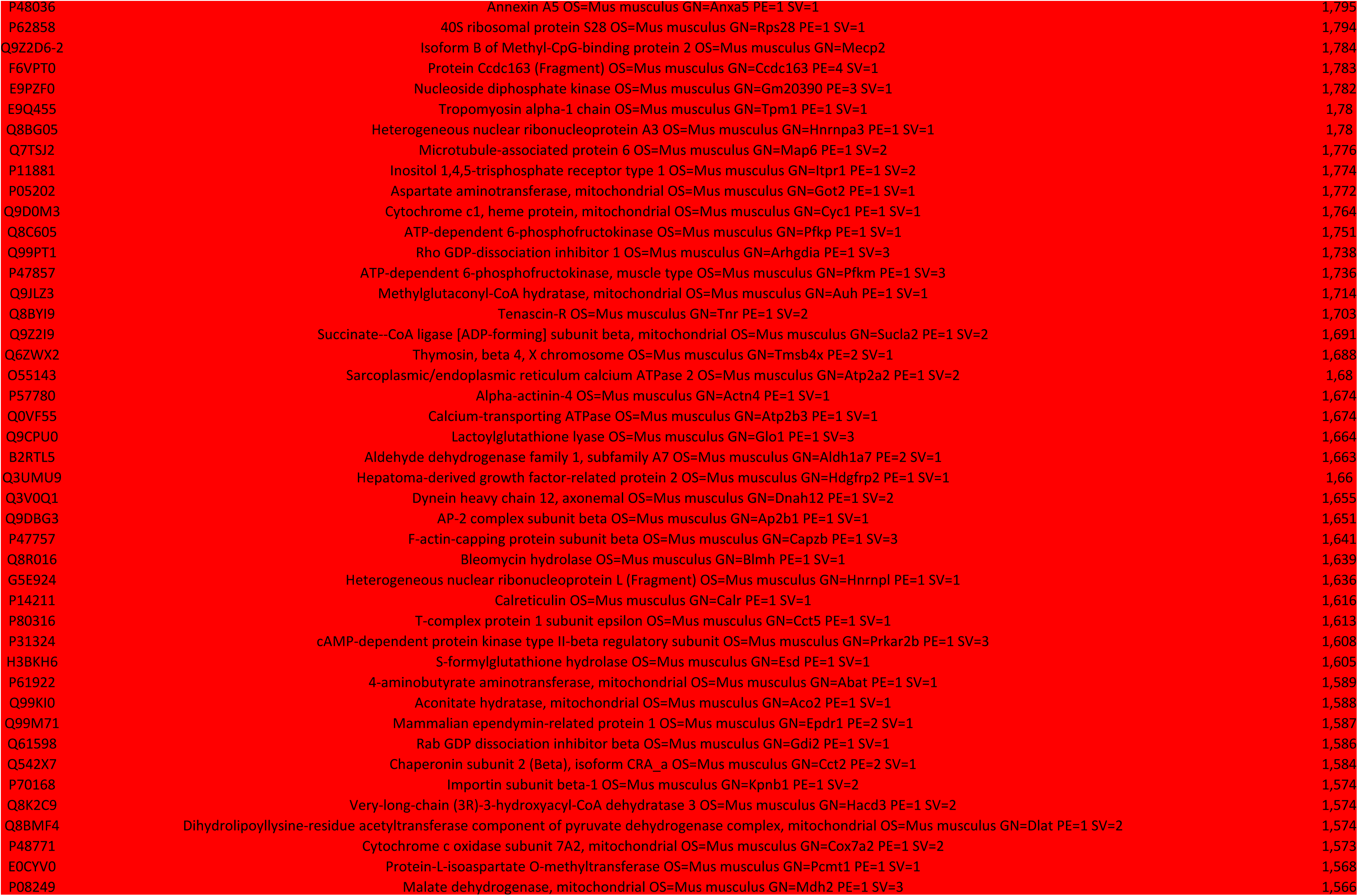

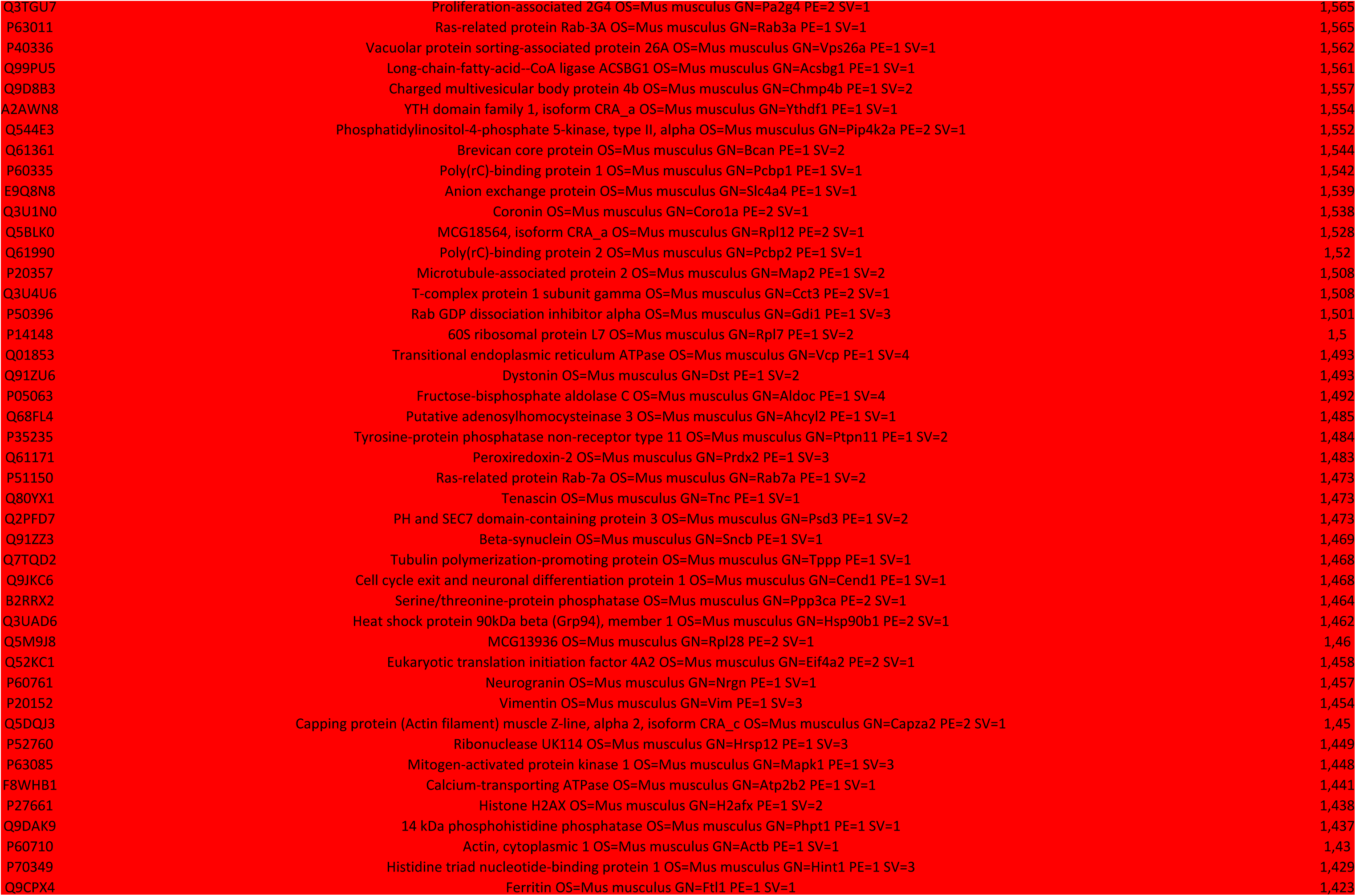

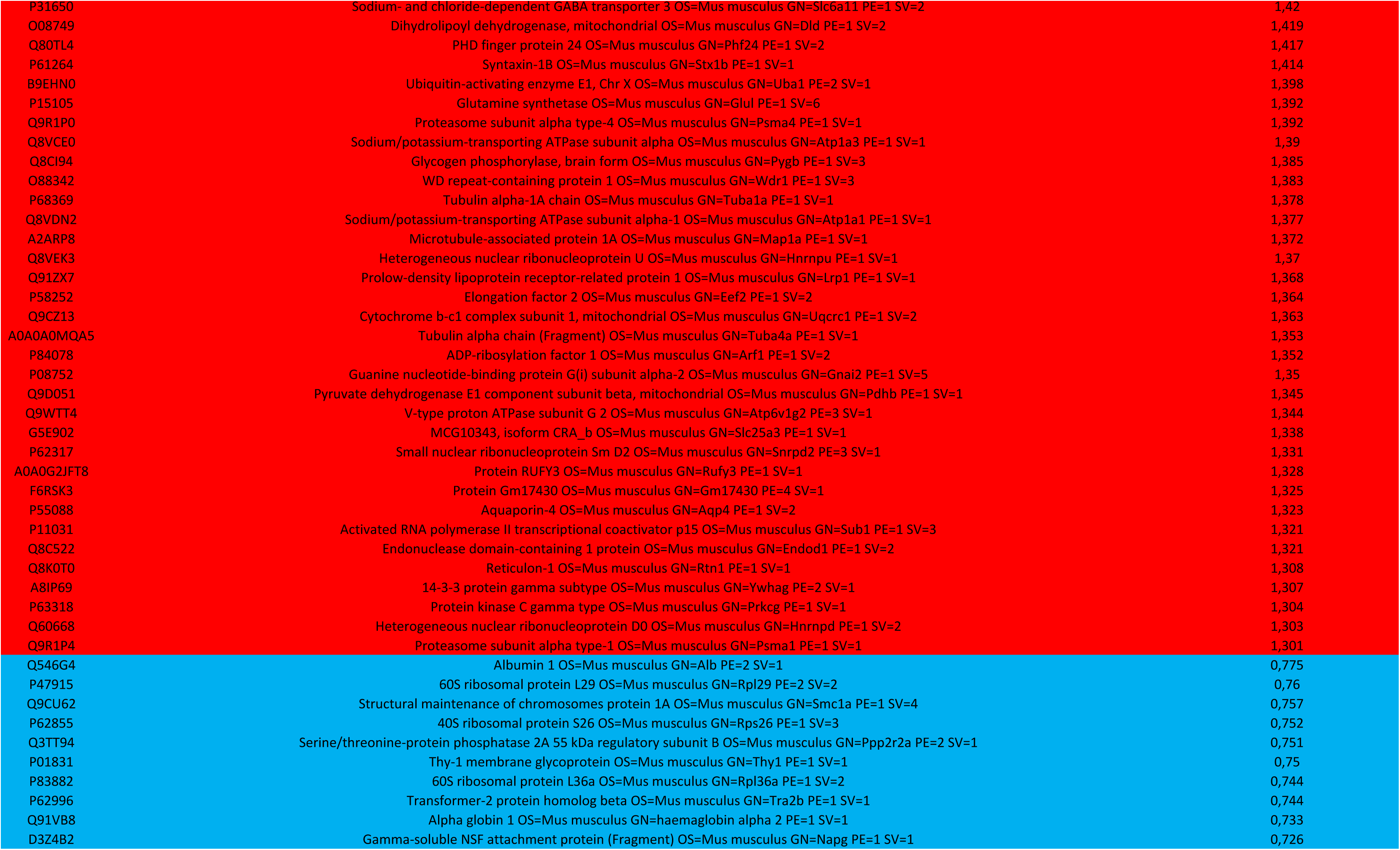

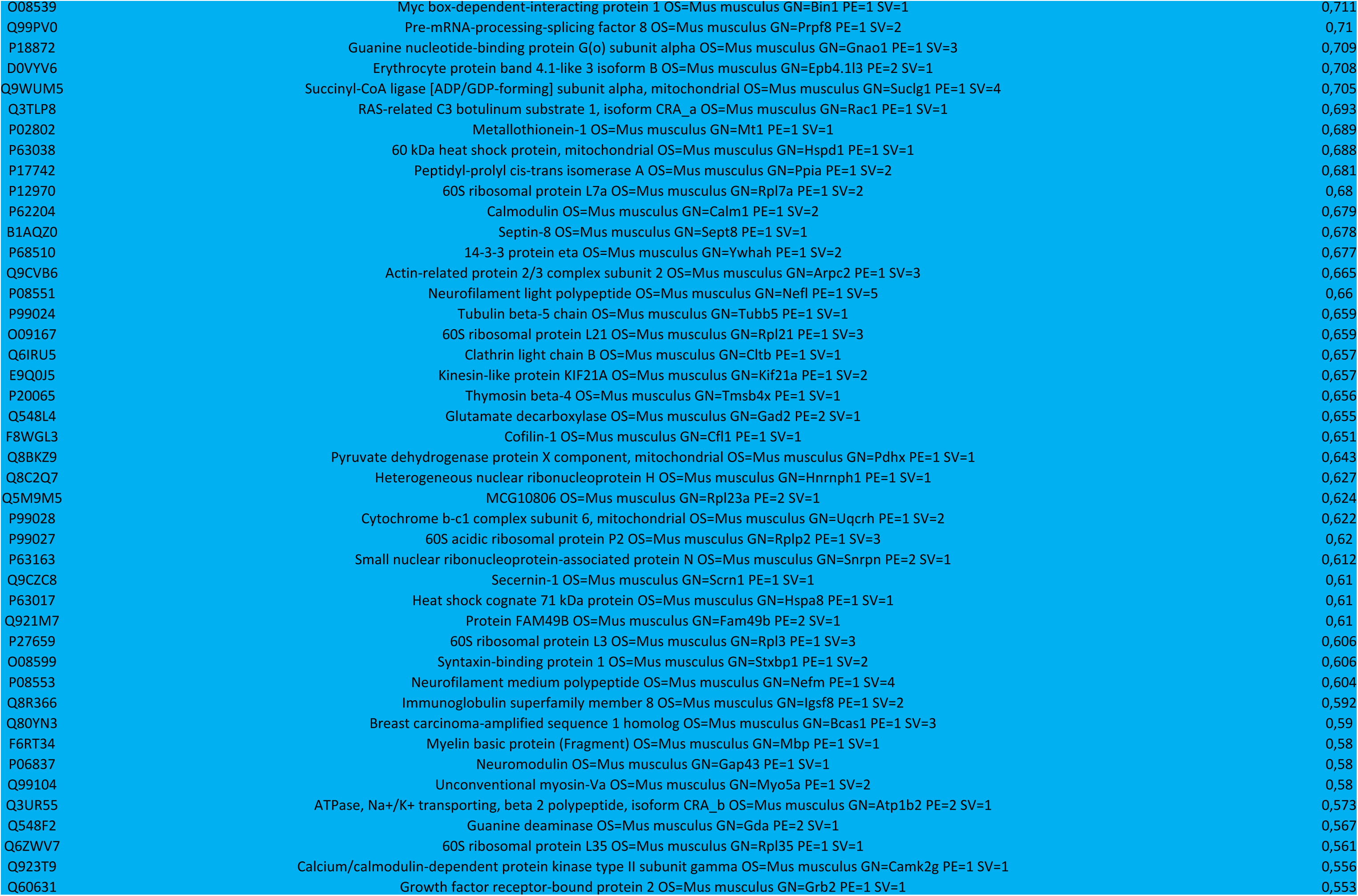

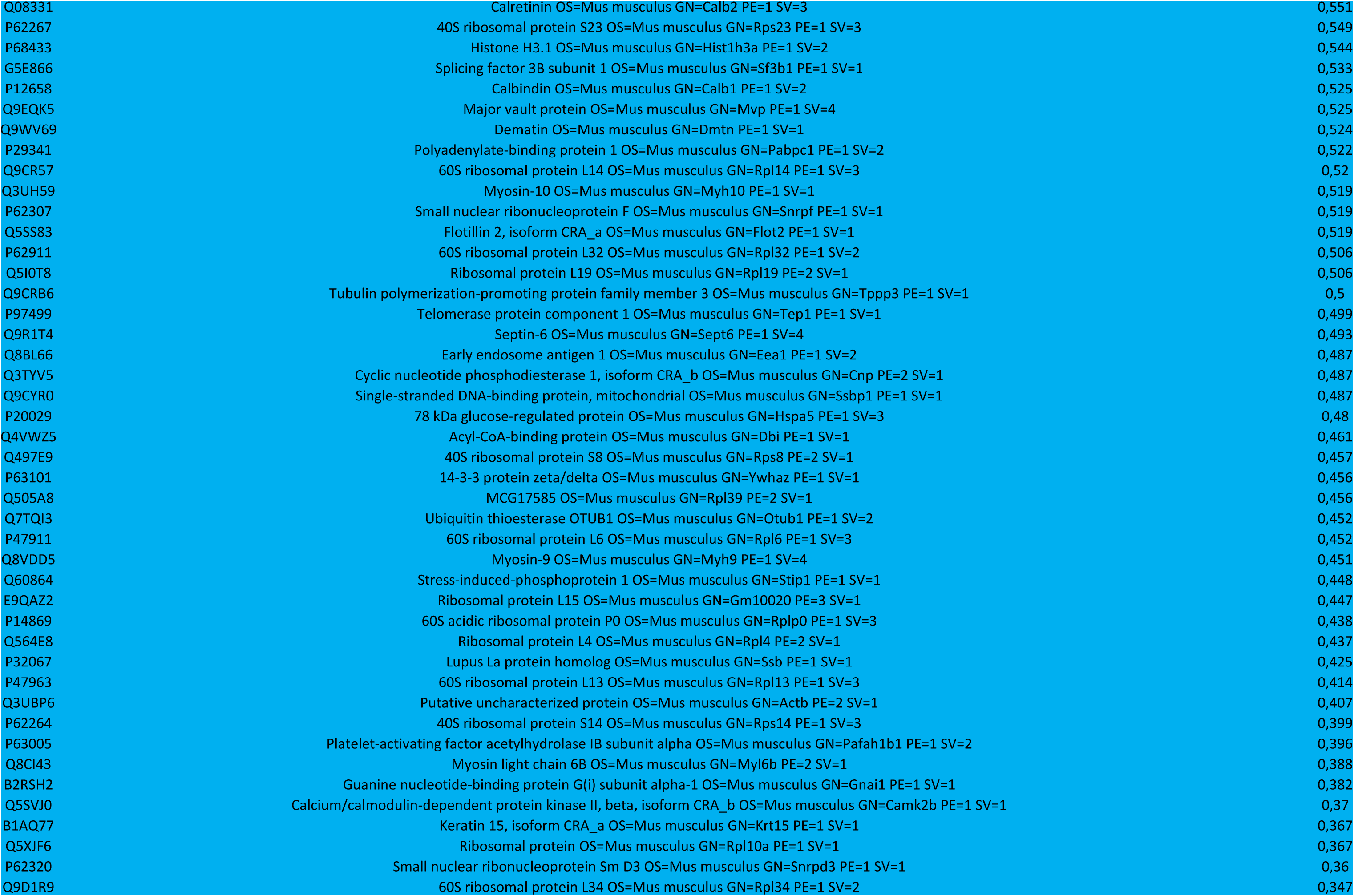

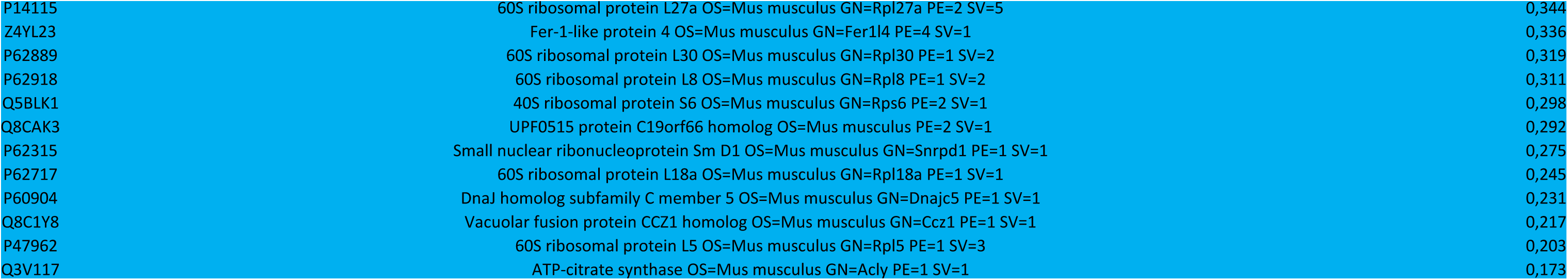

## DATA FILE S1

**REGULATED PROTEINS WT: 24 vs. 3 months**

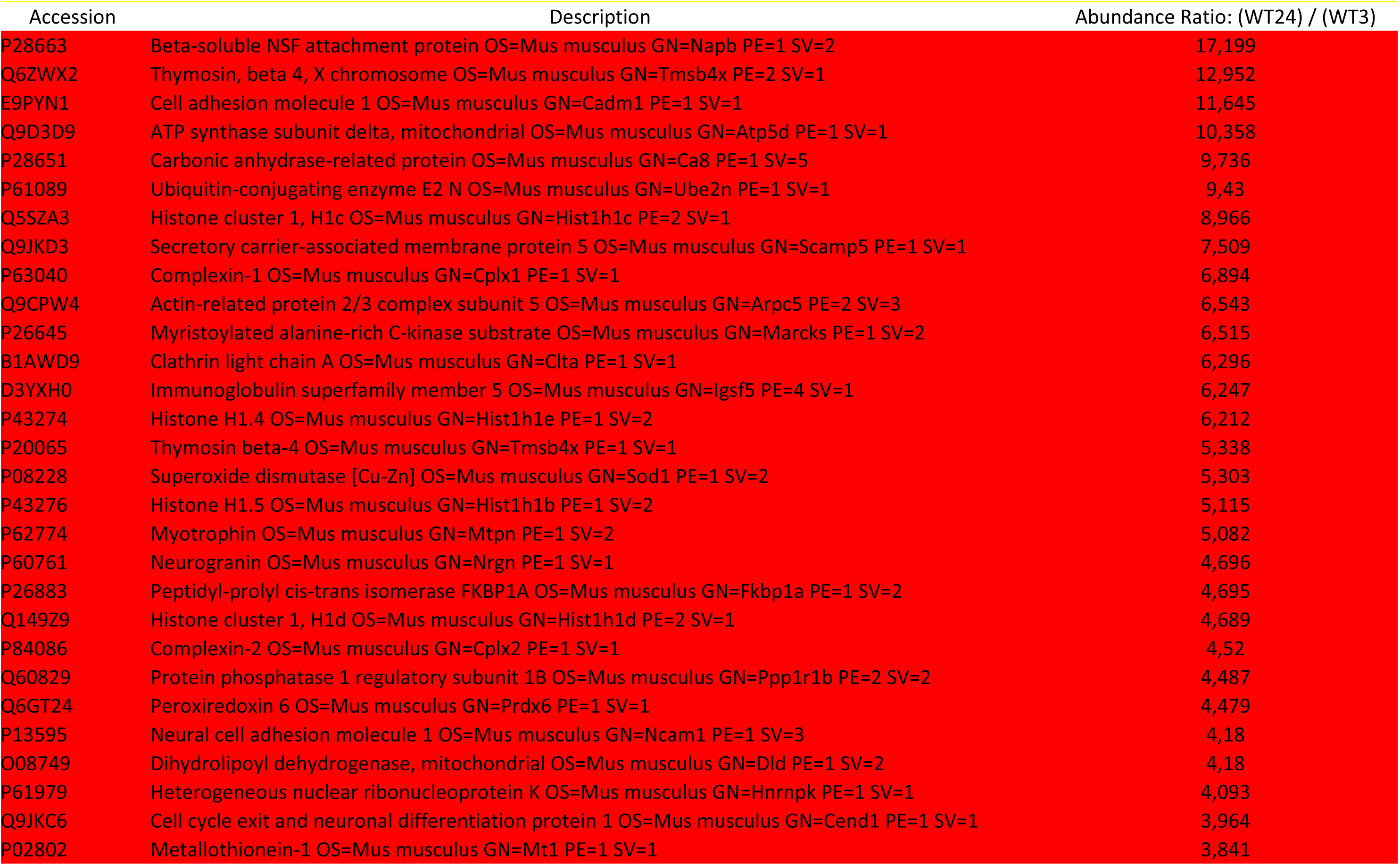

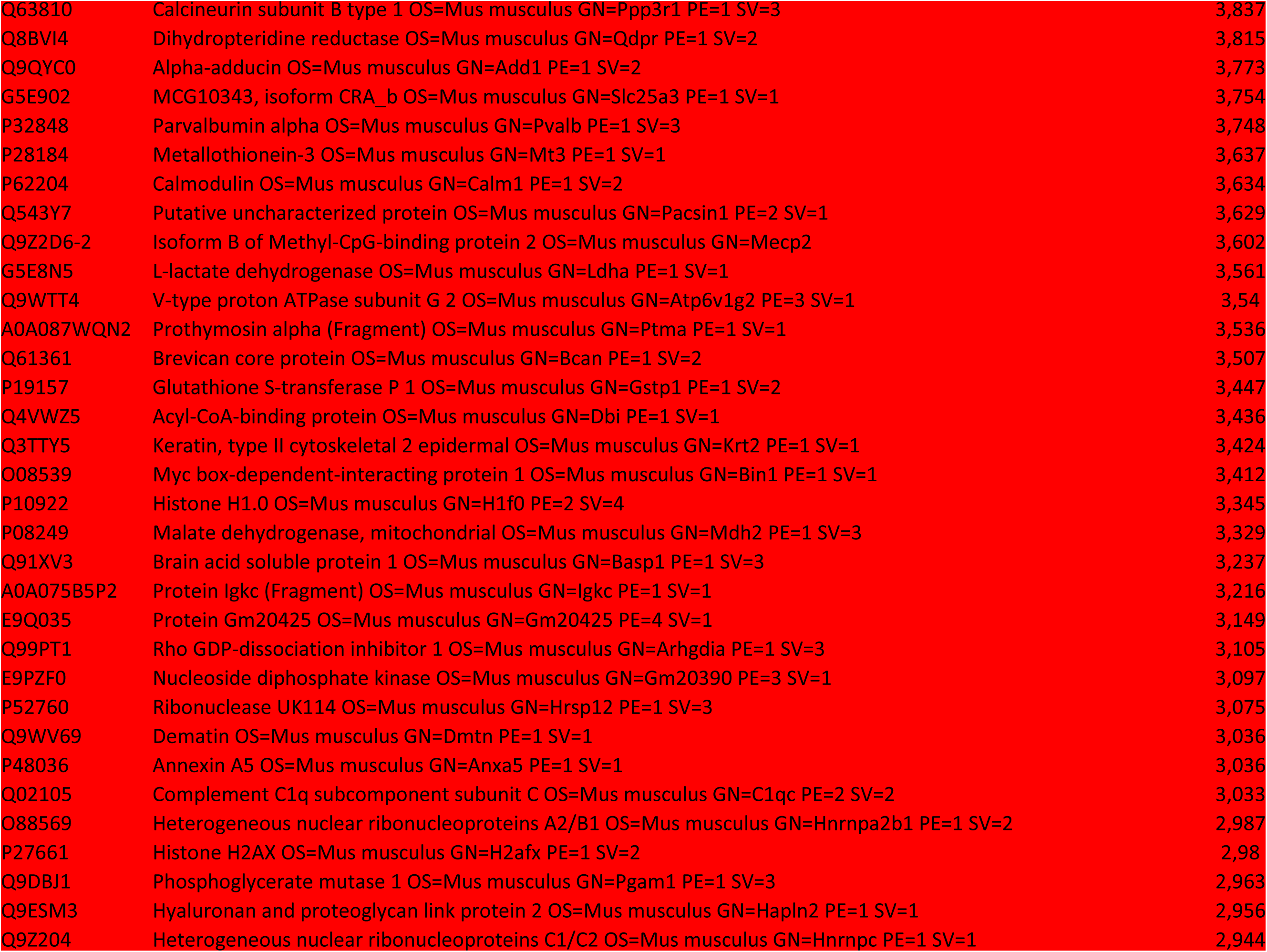

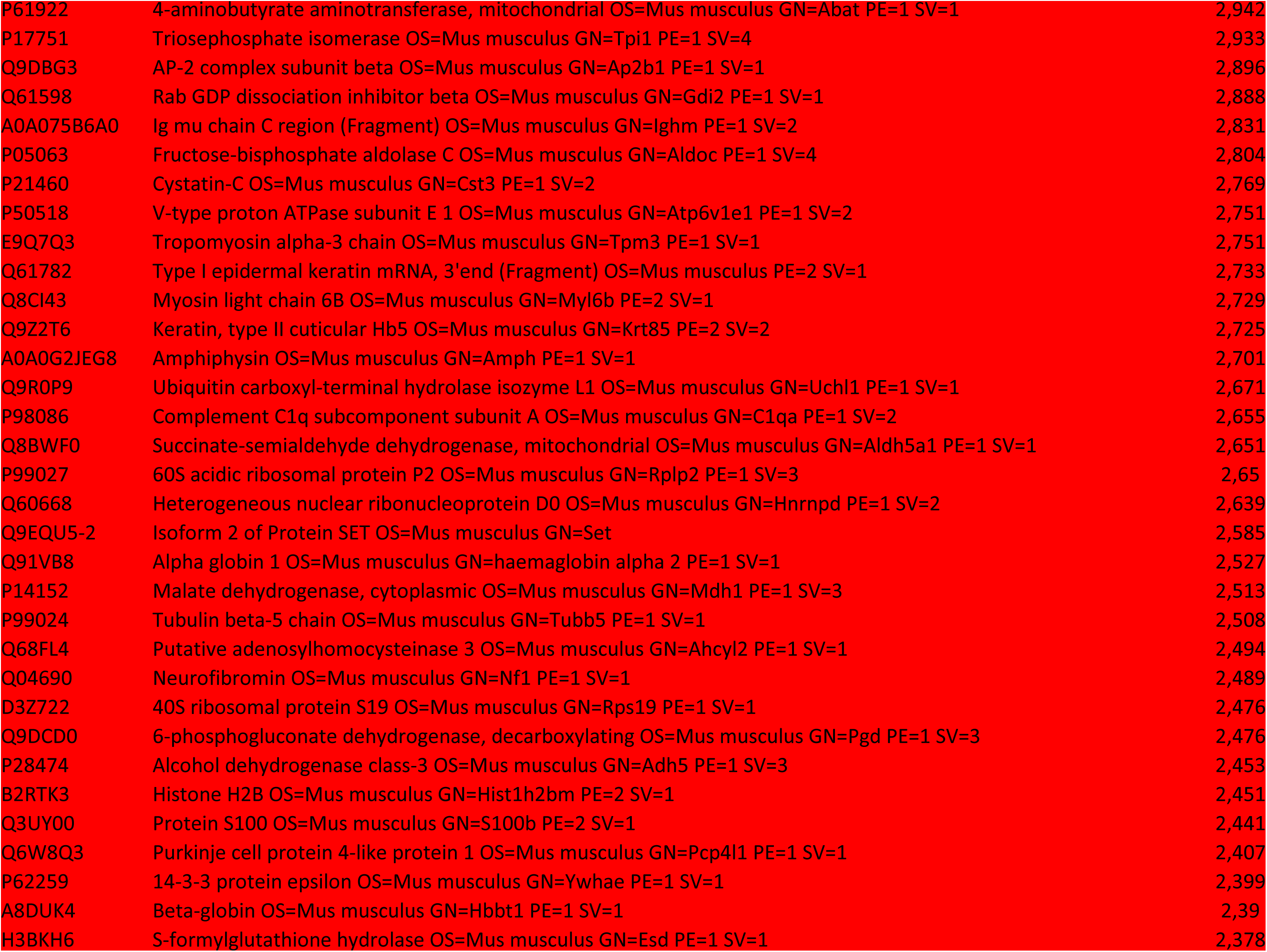

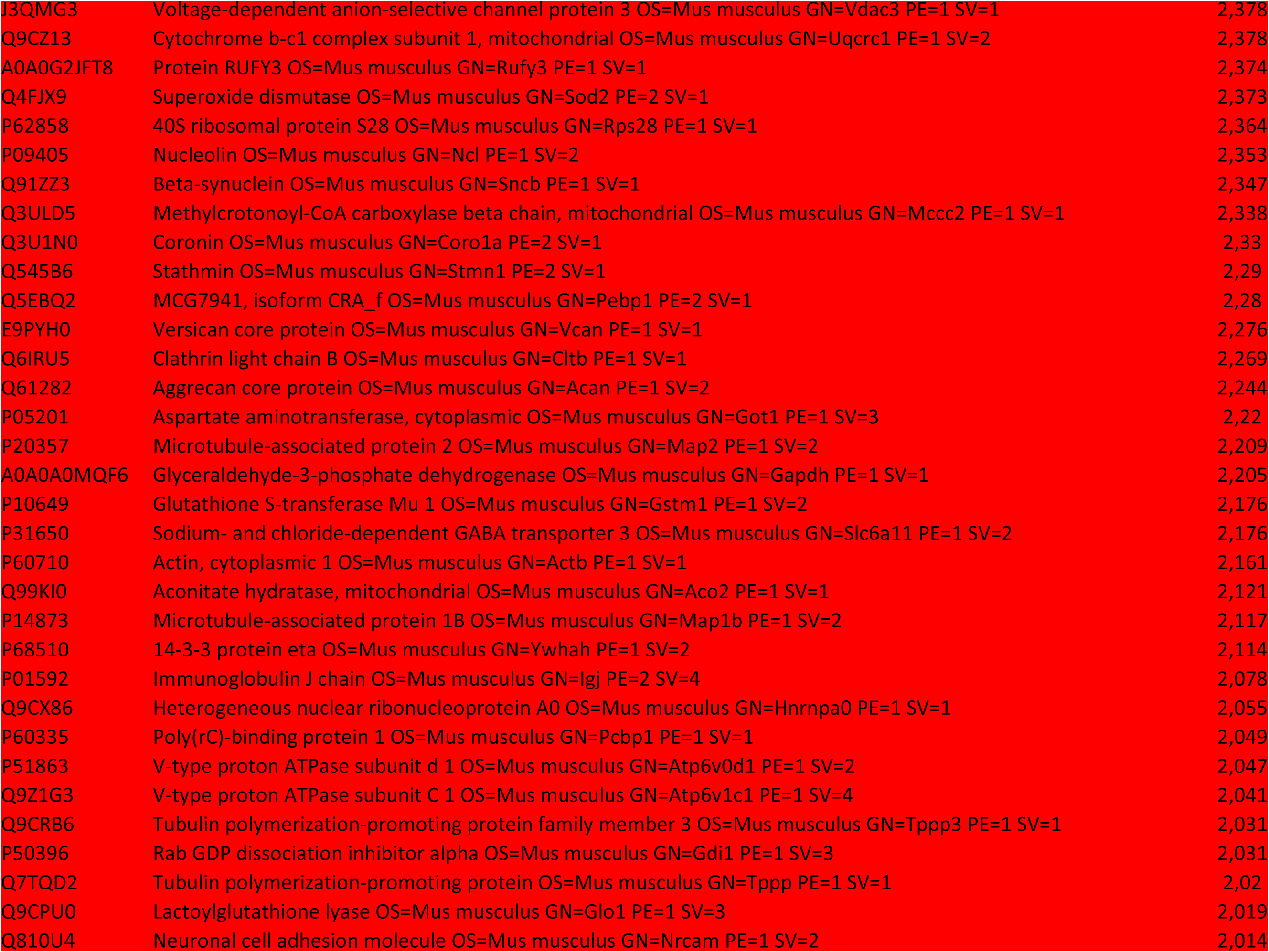

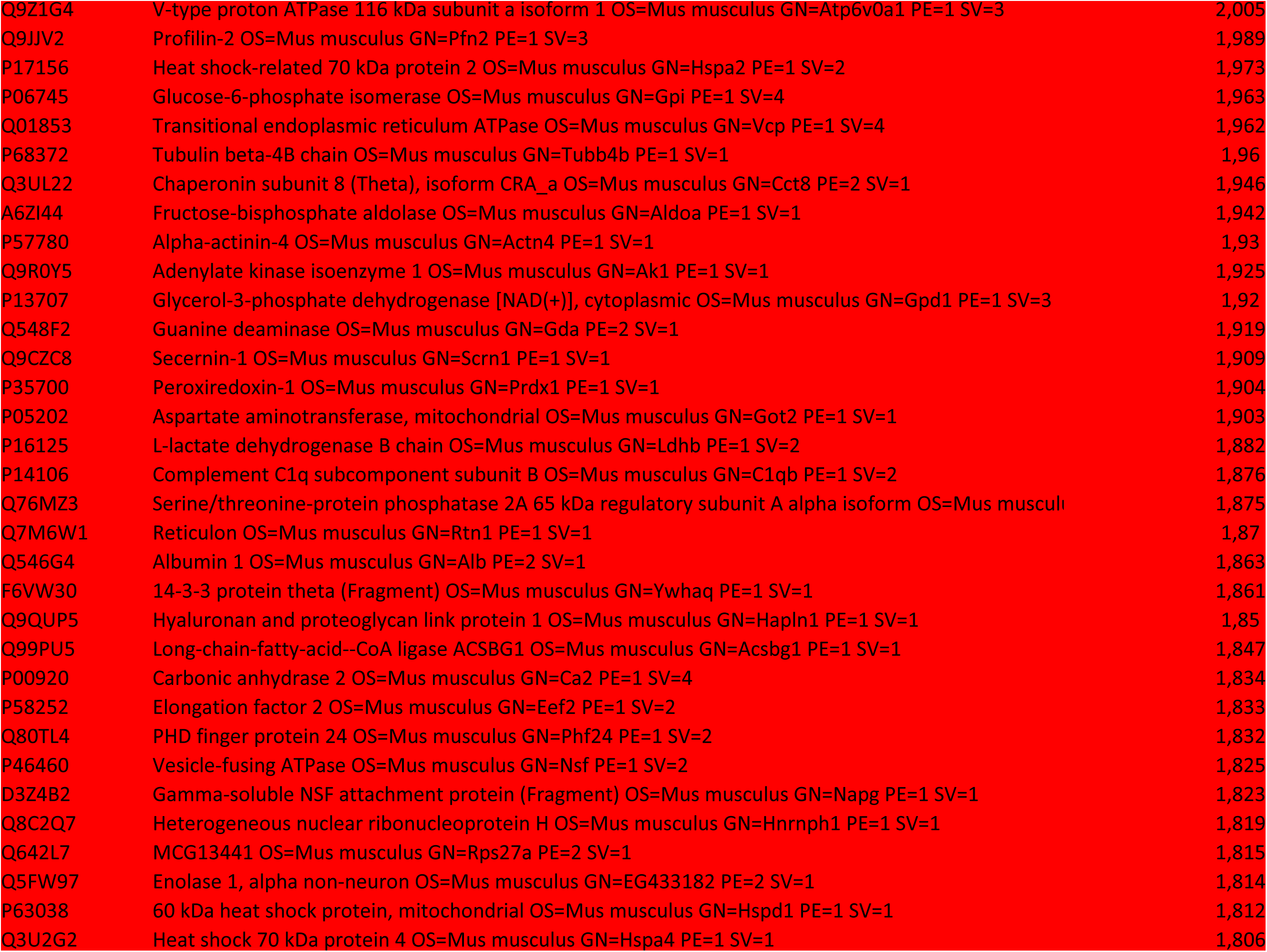

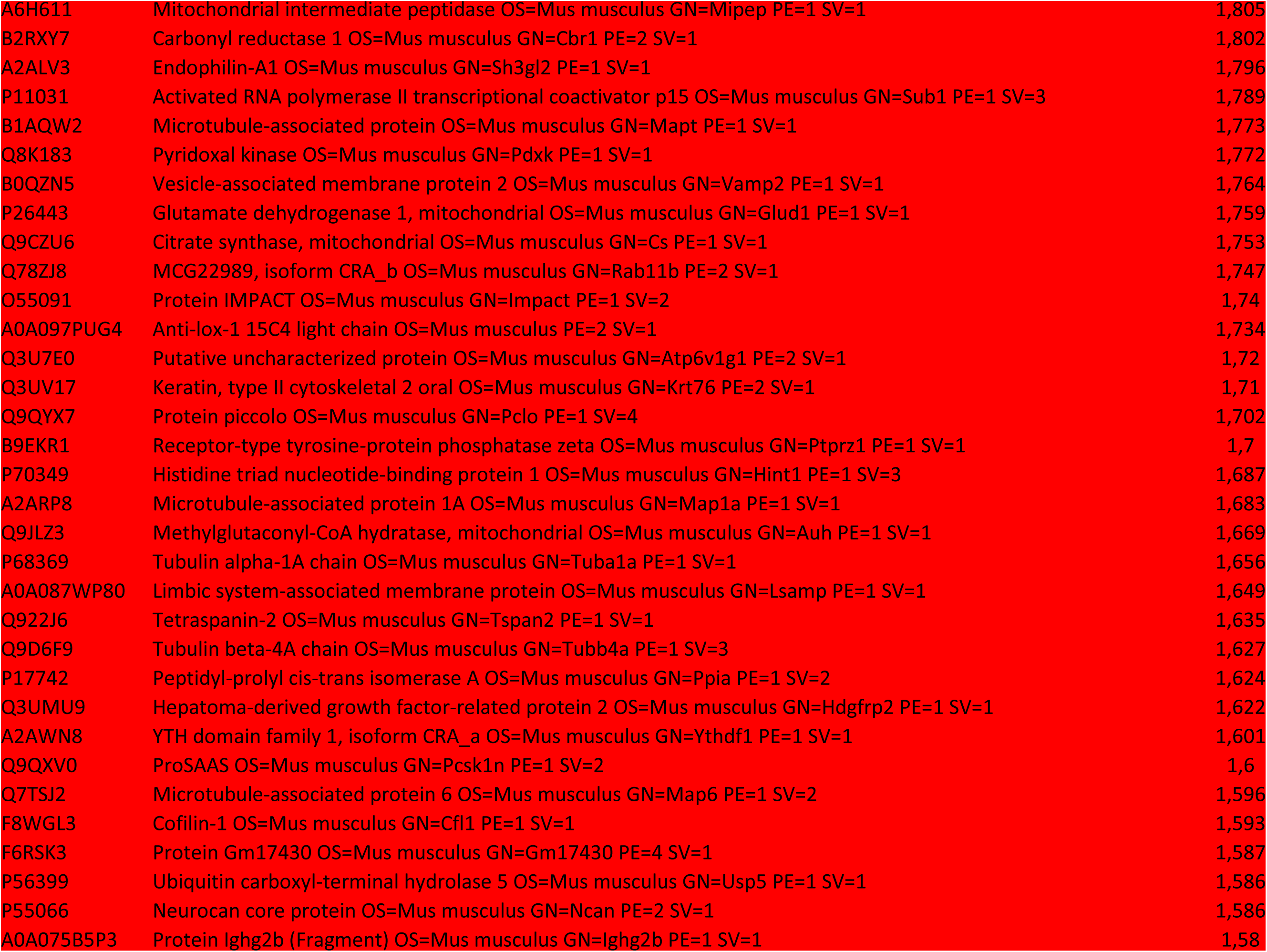

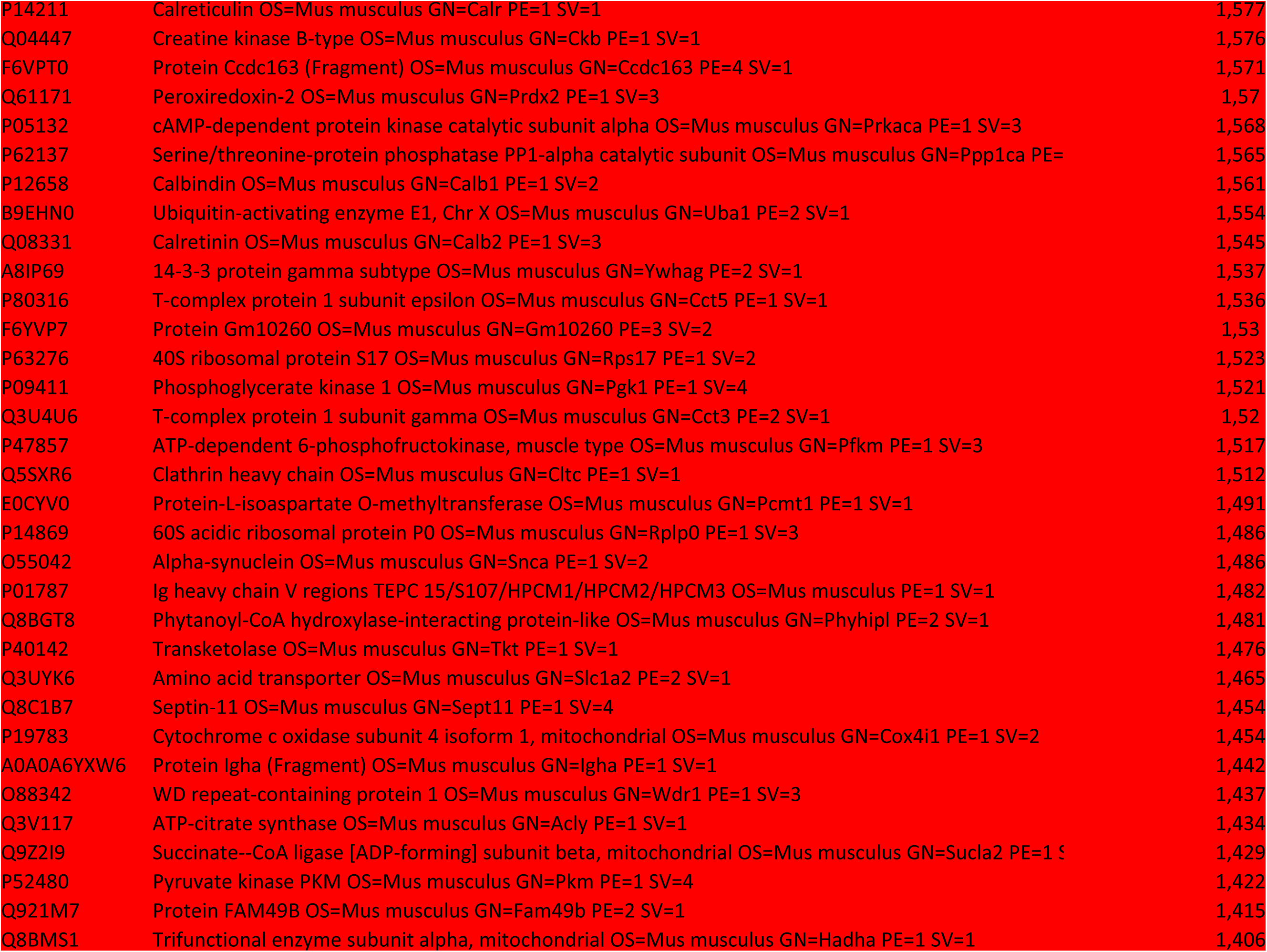

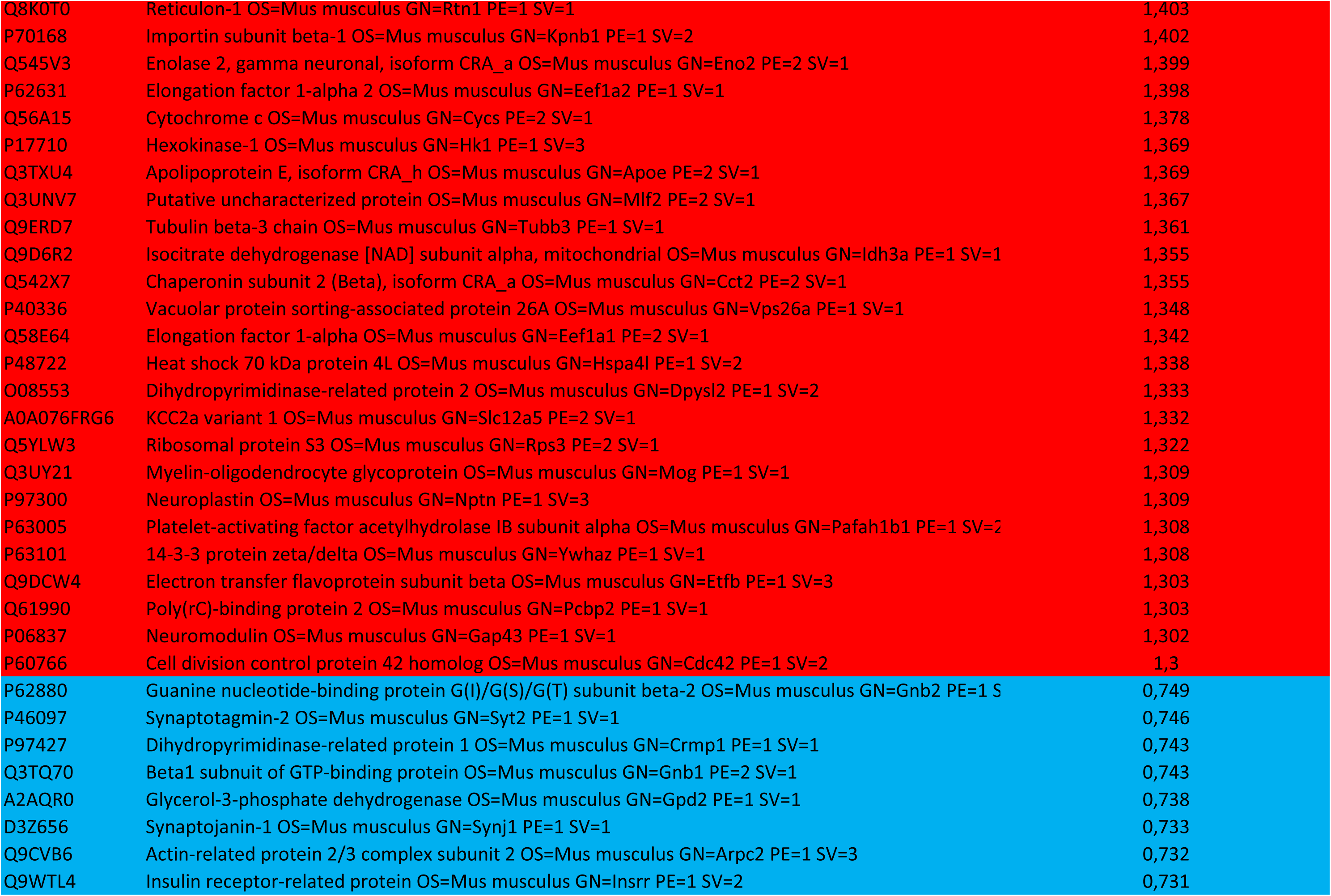

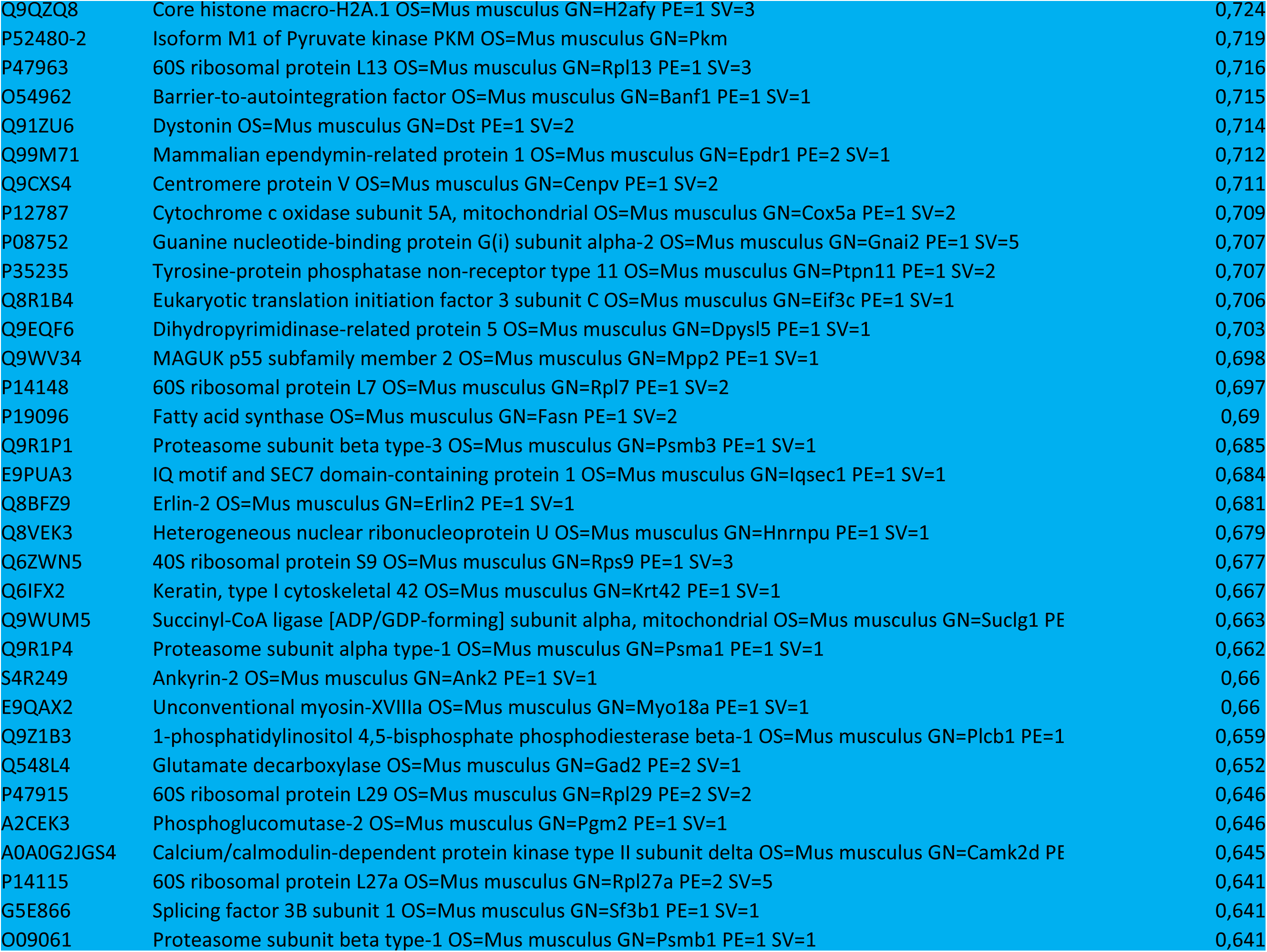

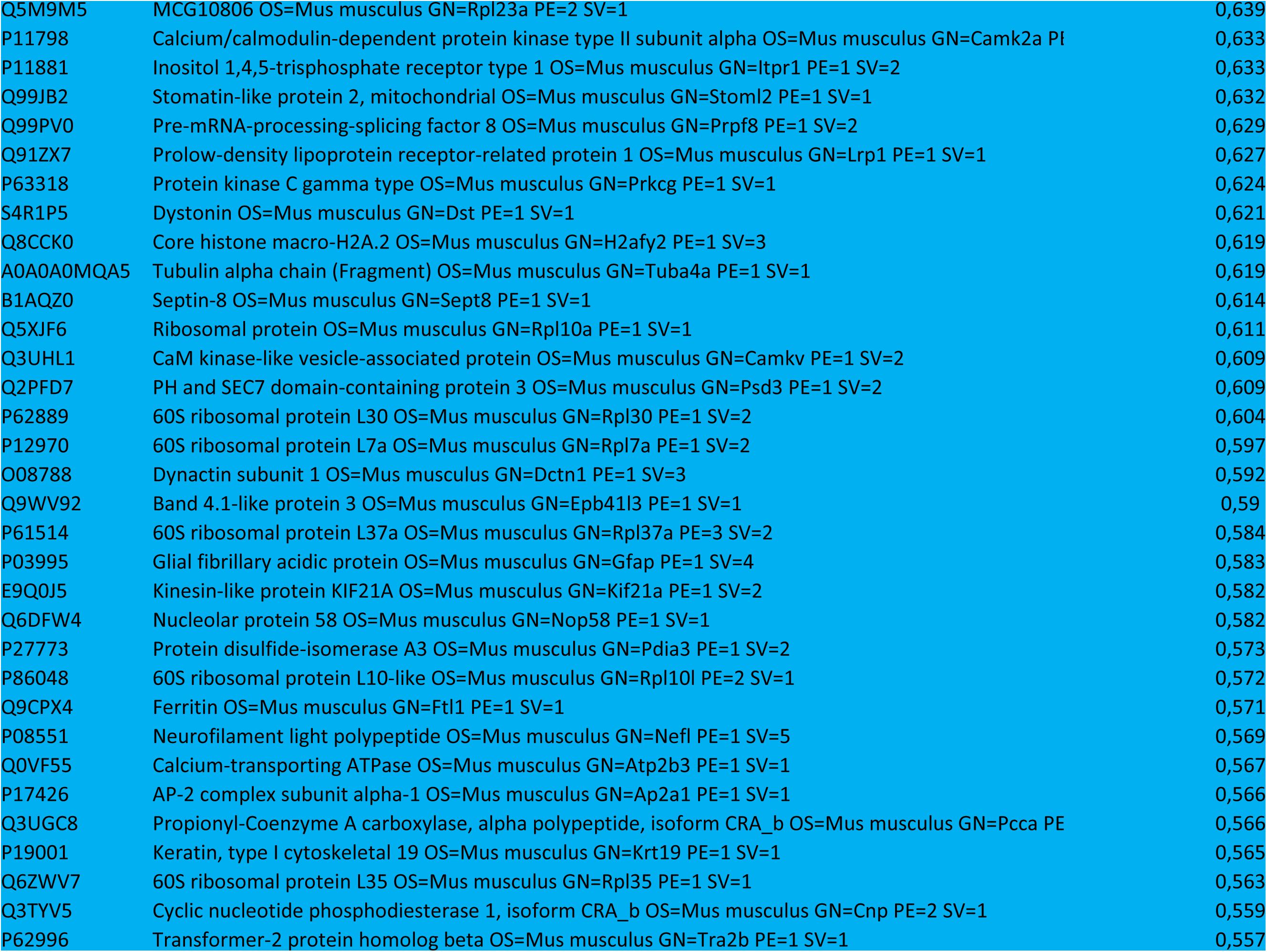

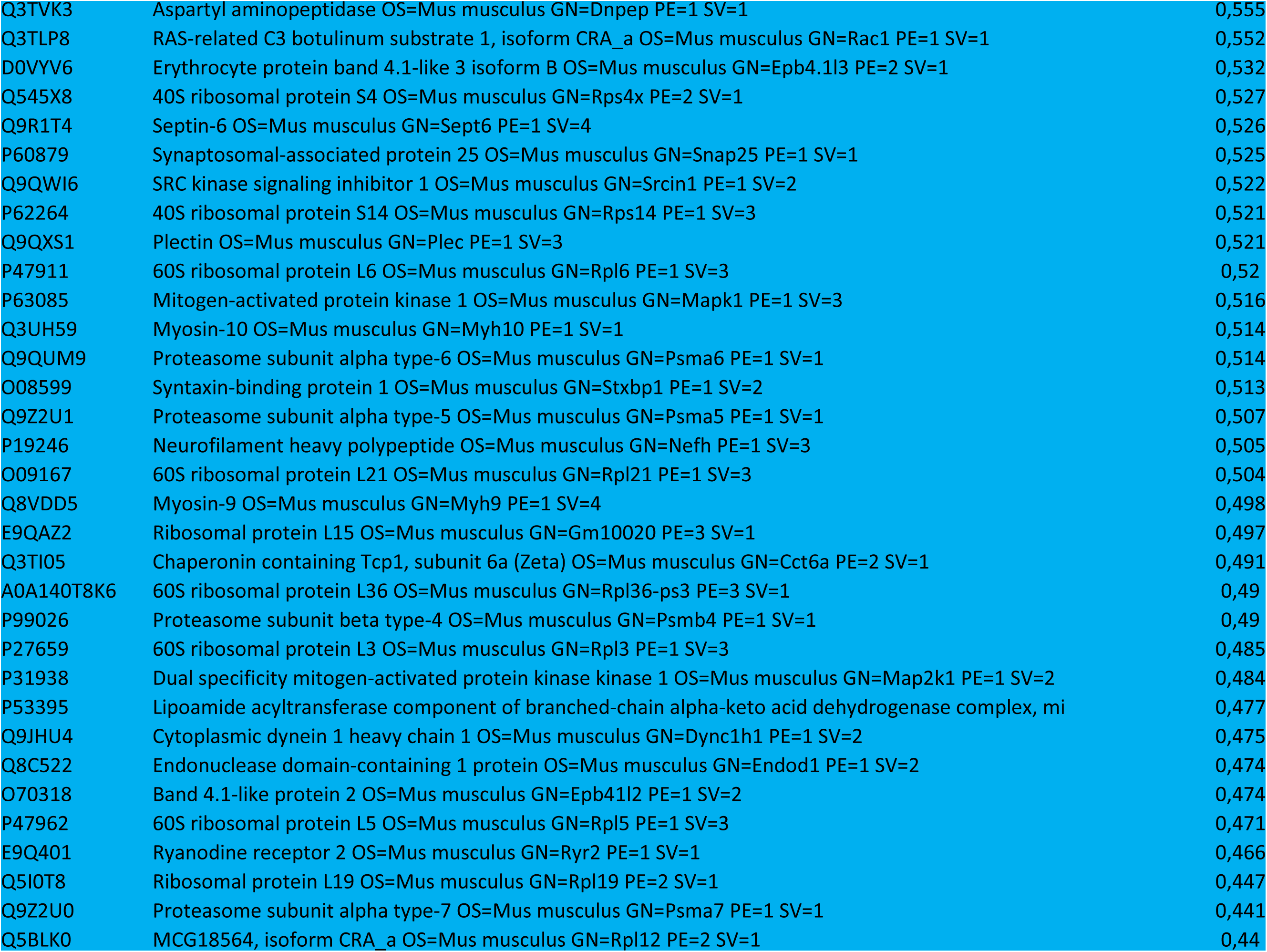

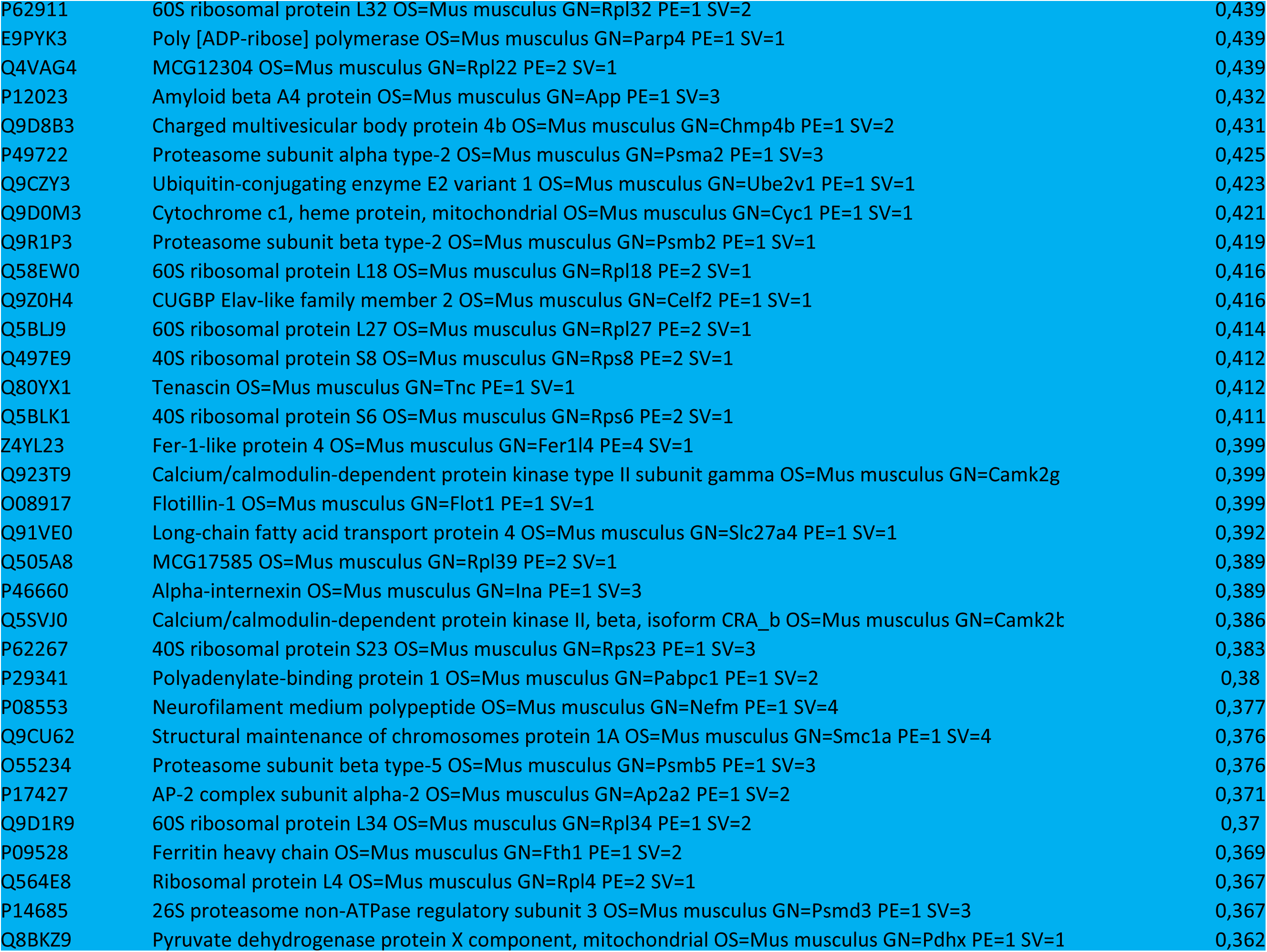

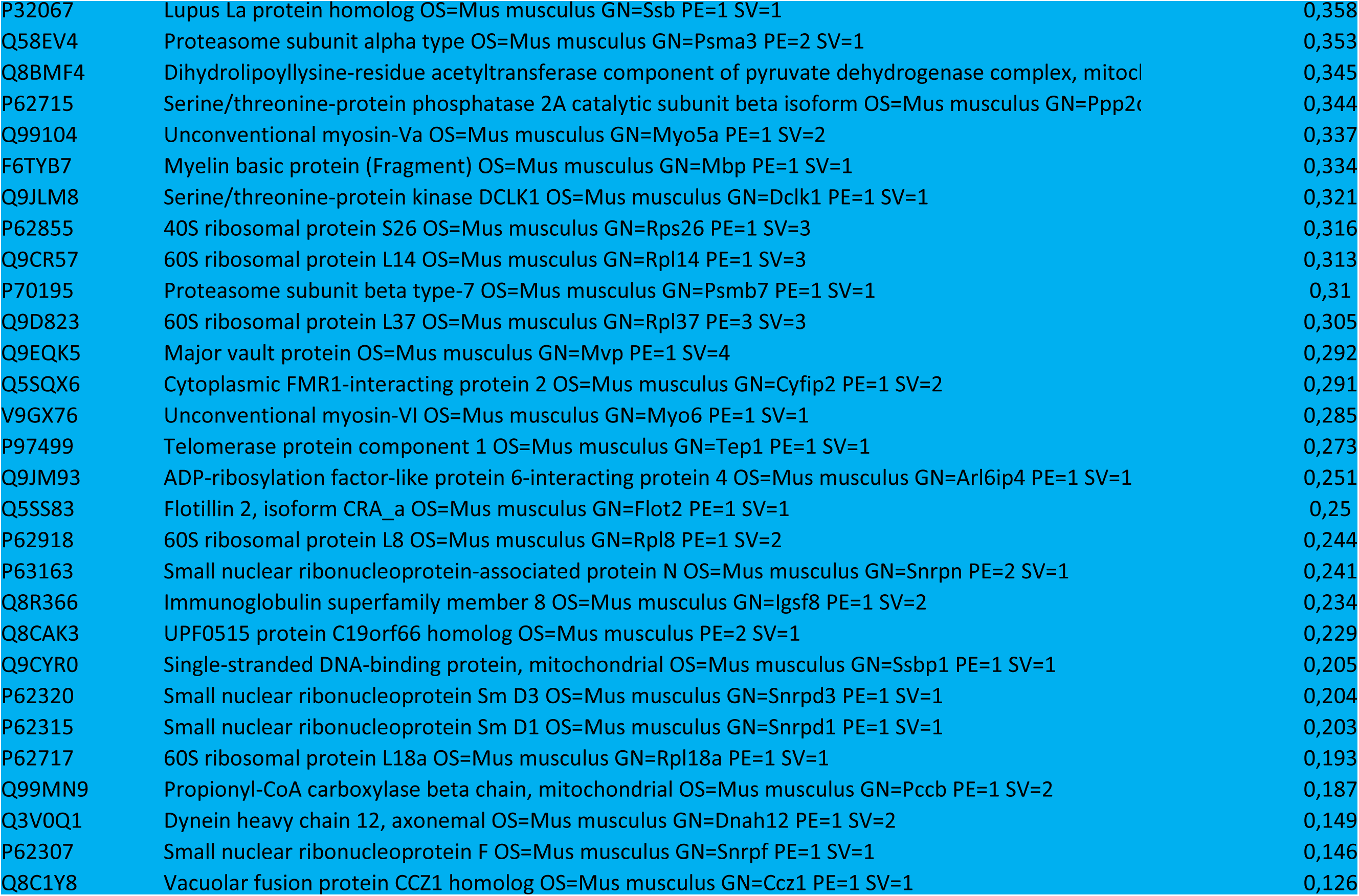

## DATA FILE S1

**REGULATED PROTEINS AD vs. non-AD**

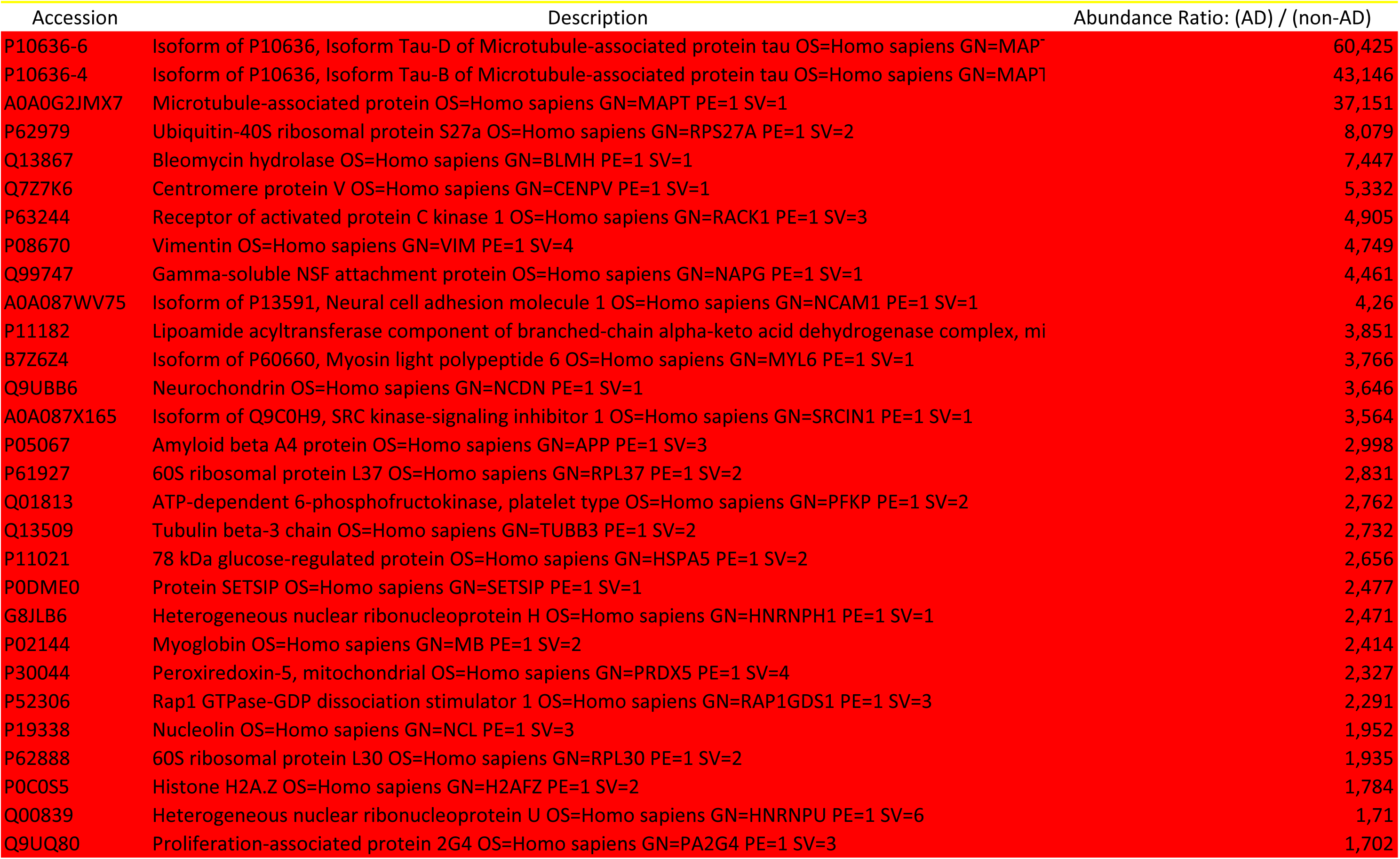

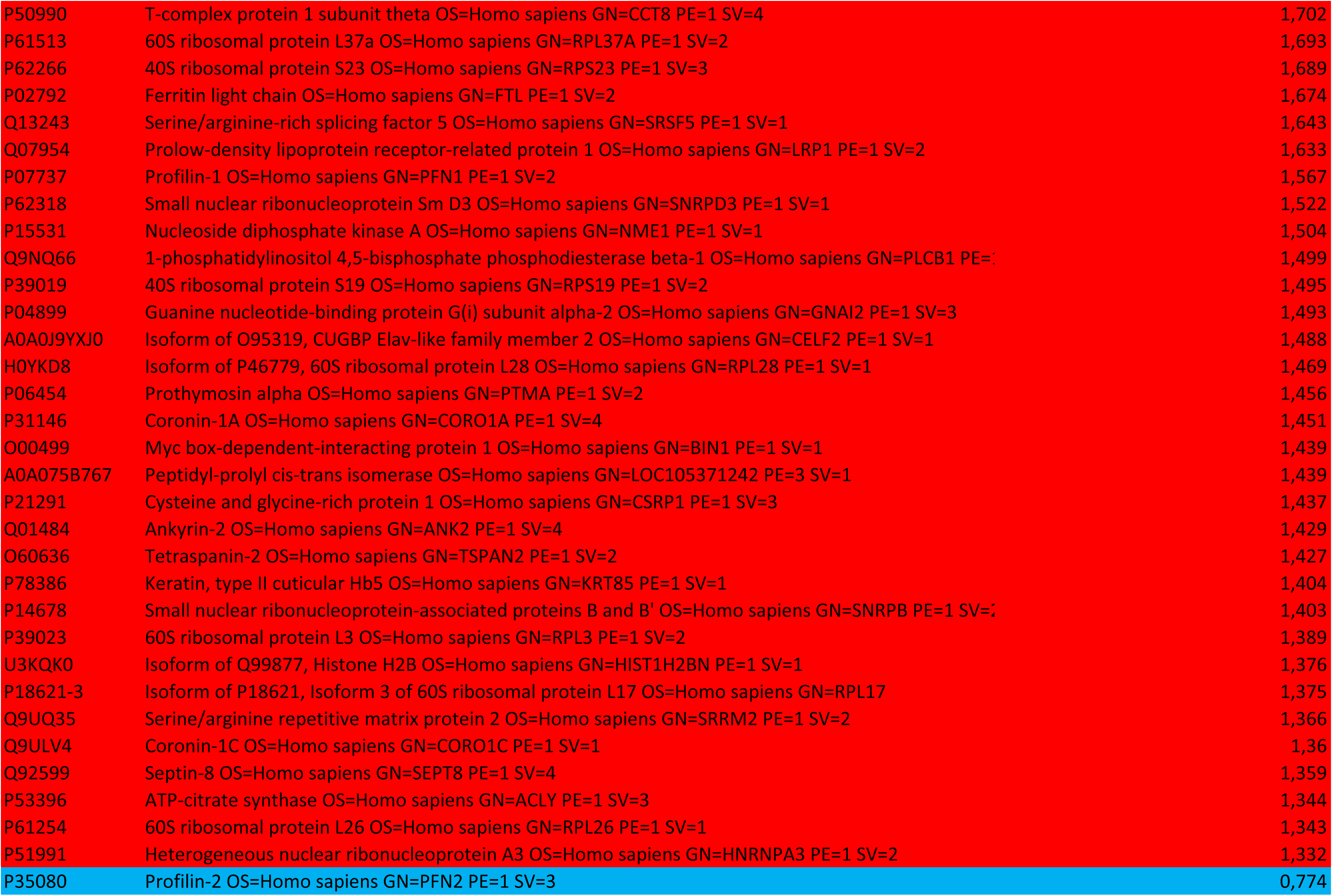

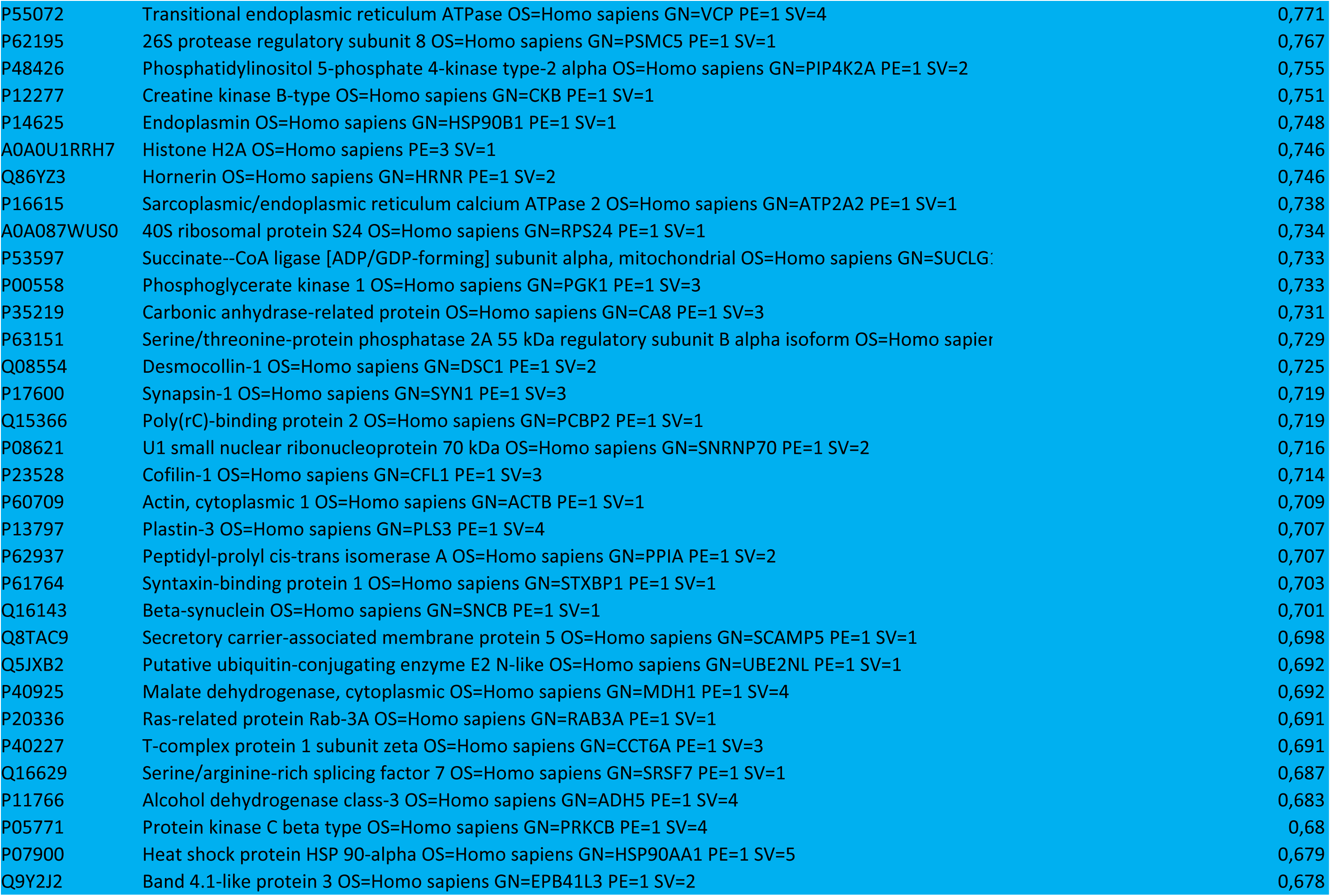

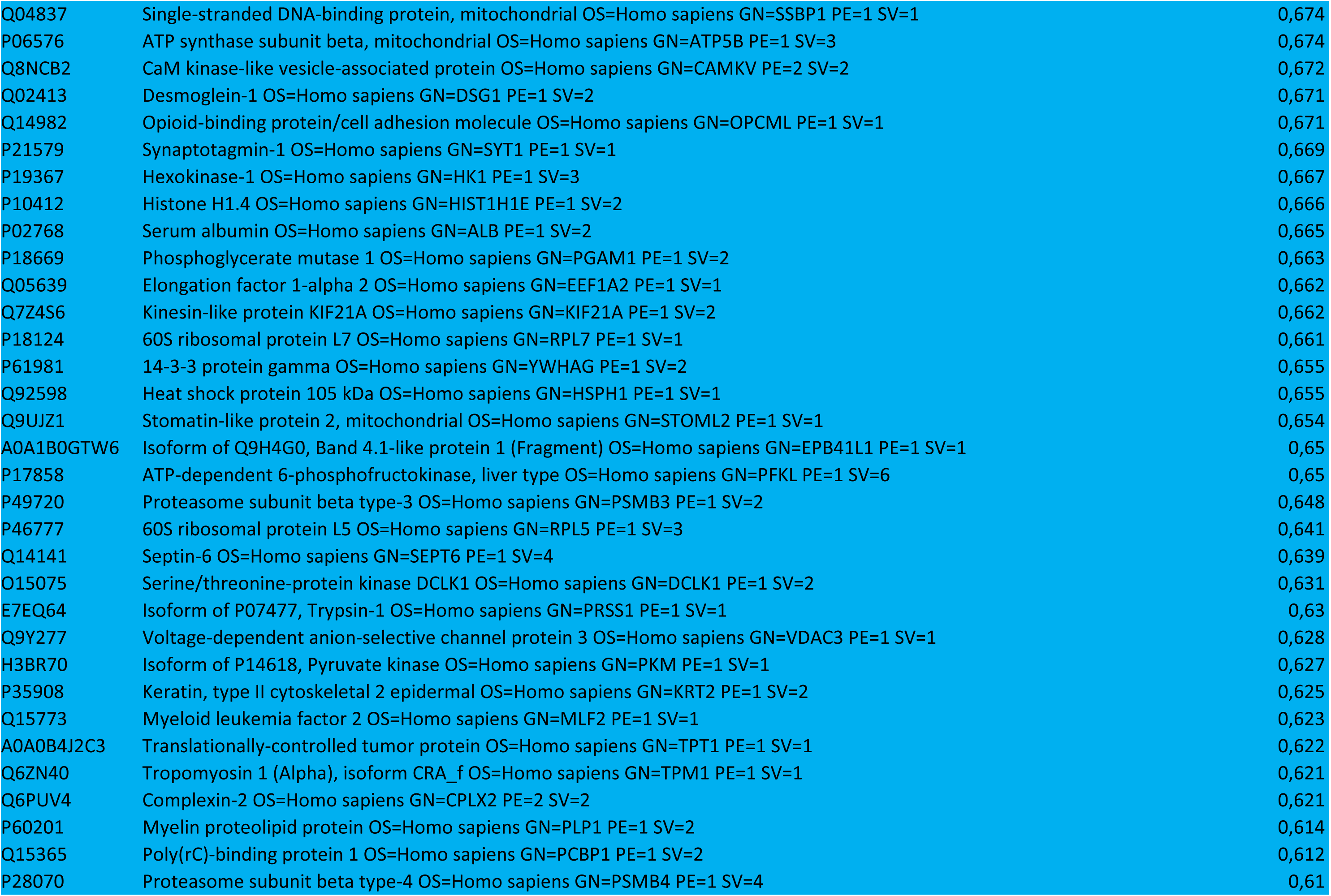

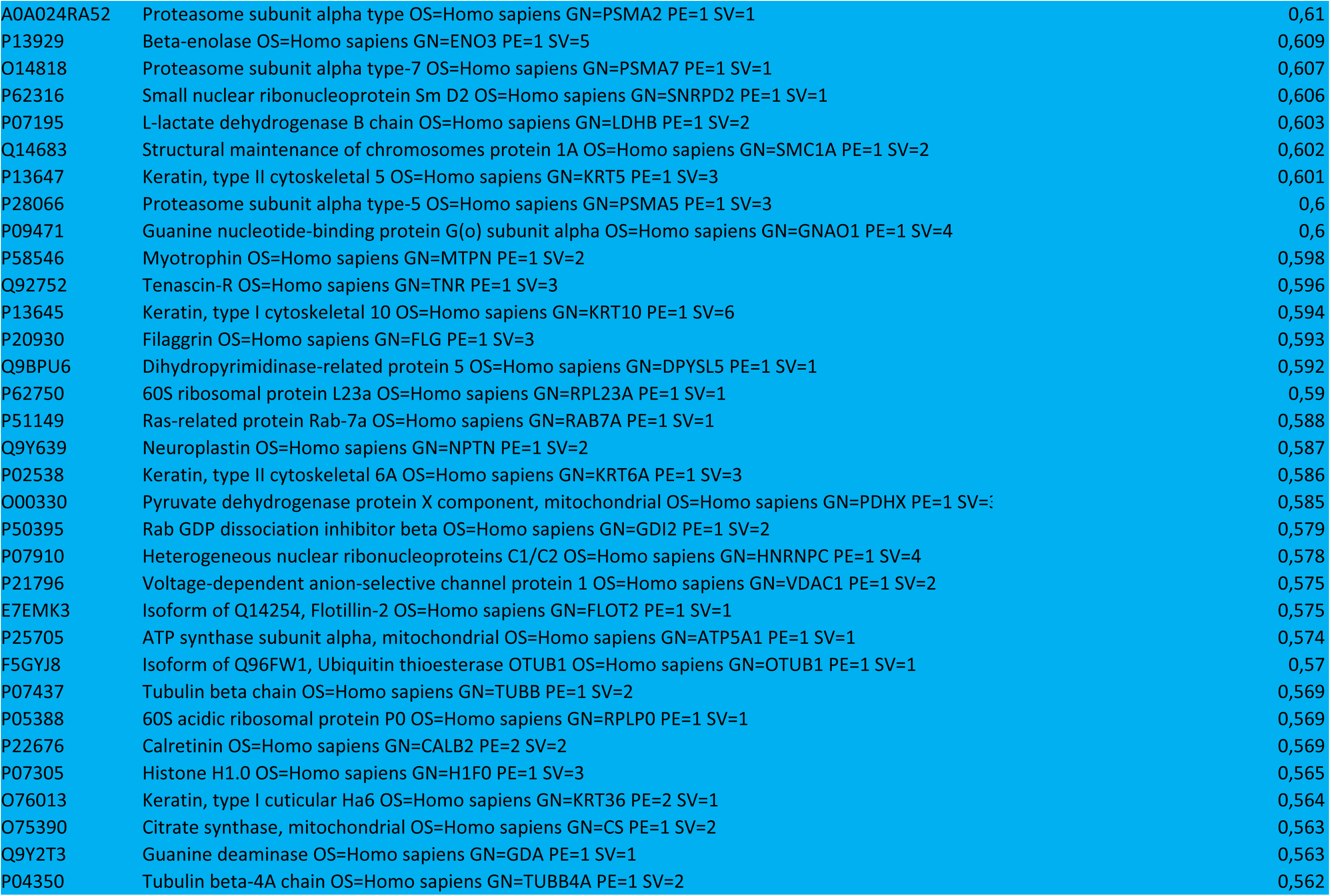

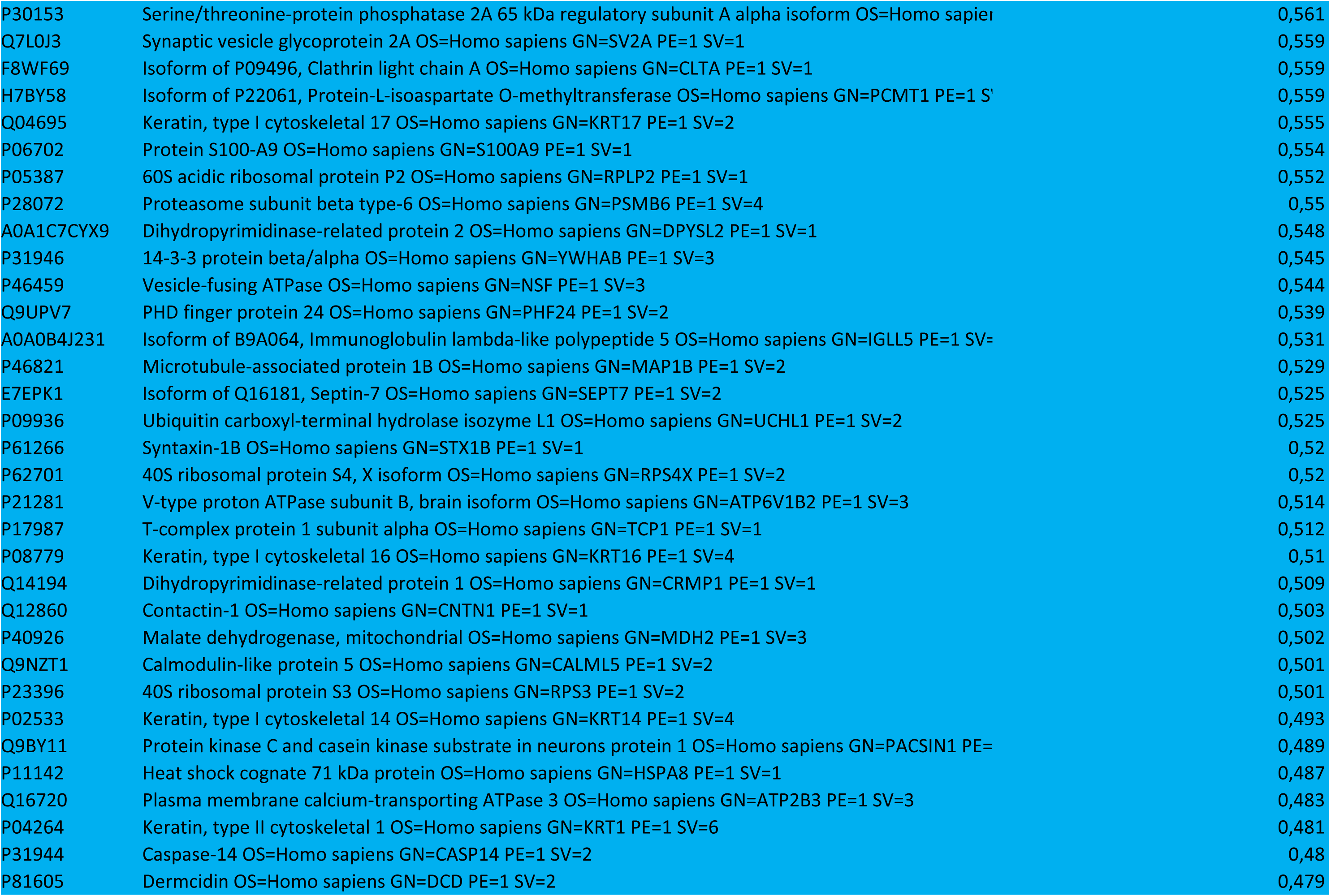

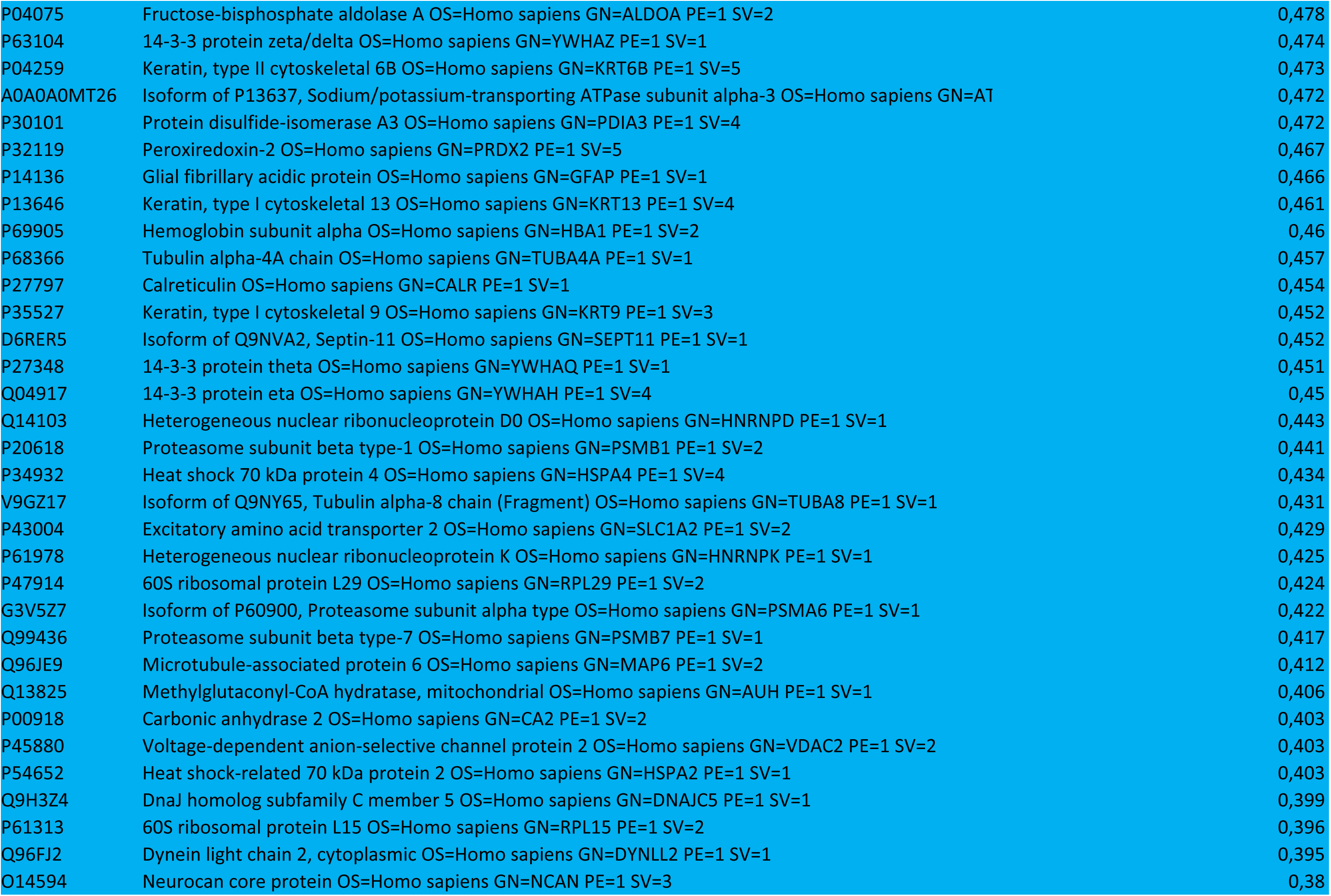

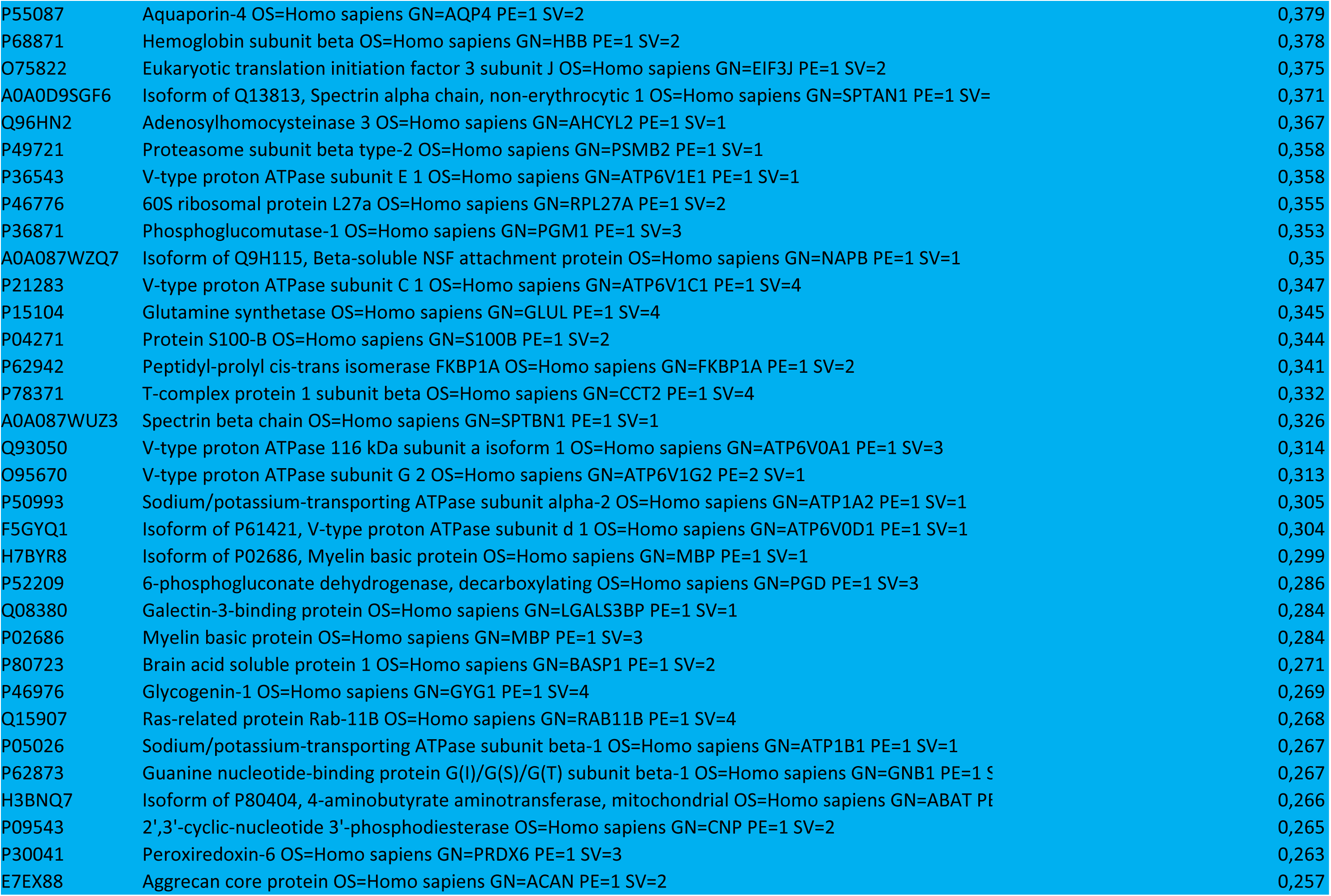

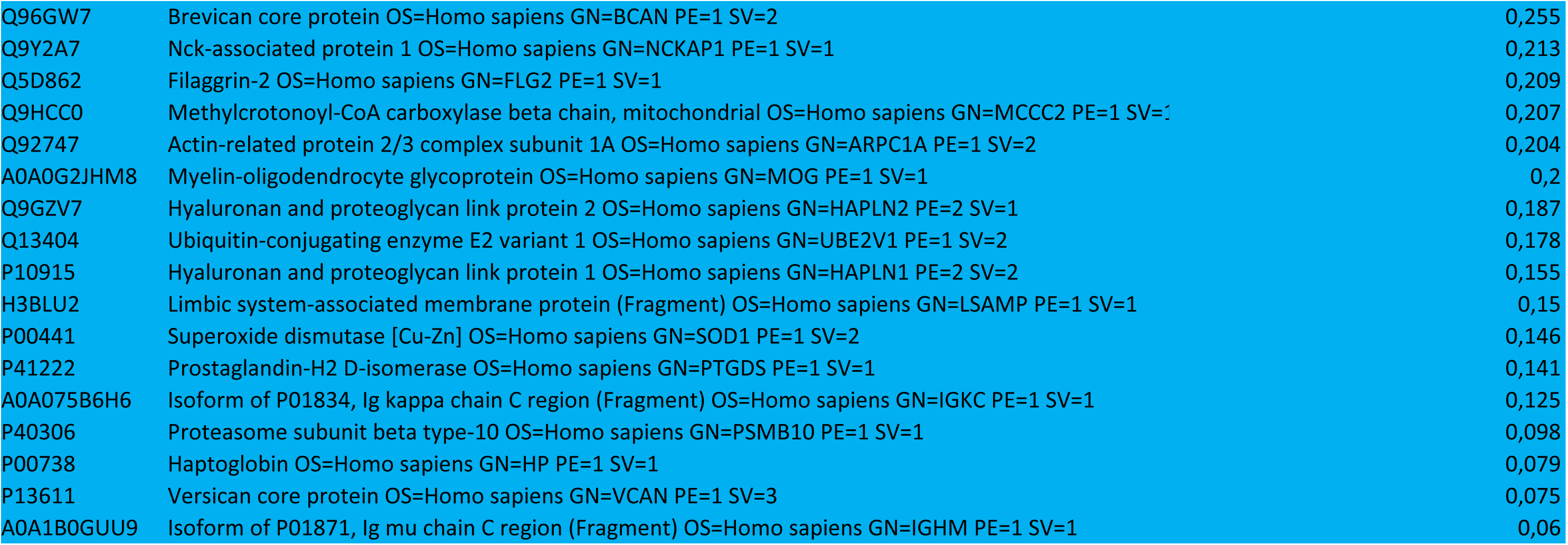

